# A Phylogenetic Model of Established and Enabled Biome Shifts

**DOI:** 10.1101/2024.08.30.610561

**Authors:** Sean W. McHugh, Michael J. Donoghue, Michael J. Landis

## Abstract

Where each species actually lives is distinct from where it could potentially survive and persist. This suggests that it may be important to distinguish established from enabled biome affinities when considering how ancestral species moved and evolved among major habitat types. We introduce a new phylogenetic method, called RFBS, to model how anagenetic and cladogenetic events cause established and enabled biome affinities (or, more generally, other discrete realized versus fundamental niche states) to shift over evolutionary timescale. We provide practical guidelines for how to assign established and enabled biome affinity states to extant taxa, using the flowering plant clade Viburnum as a case study. Through a battery of simulation experiments, we show that RFBS performs well, even when we have realistically imperfect knowledge of enabled biome affinities for most analyzed species. We also show that RFBS reliably discerns established from enabled affinities, with similar accuracy to standard competing models that ignore the existence of enabled biome affinities. Lastly, we apply RFBS to Viburnum to infer ancestral biomes throughout the tree and to highlight instances where repeated shifts between established affinities for warm and cold temperate forest biomes were enabled by a stable and slowly-evolving enabled affinity for both temperate biomes.

## Introduction

Biomes are environmentally consistent spatial assemblages of ecologically similar co-occurring species (Jiang et al. 2017; Hunter et al. 2021). Species ranges are often constrained by biomes, with most species occupying only a few of all resident biomes within a region. To occupy and persist in a biome, species must both successfully compete within the species assemblage and physiologically tolerate local environmental conditions (Moncrieff et al. 2016; Mucina 2019; Lu et al. 2022). Those with long-term persistence within a biome are typically identified by biologists as having a phenotypic “affinity” to it (Palmer 1990; Schrire et al. 2009; Jansson et al. 2013; Dale et al. 2020), consistent with pre- or post-adaptation to the biome.

Historical evidence strongly suggests that historical “shifts” in the set of biome affinities for a species are common and important in shaping current global distributions of biodiversity (Crisp et al. 2009; Donoghue and Edwards 2014; Antonelli et al. 2018; Willink et al. 2024). These shifts may manifest as expansions or contractions on the set of biomes affinities of a species, where the shifts are frequently accompanied by other important historical events, including ecological phenotypic shifts, long distance dispersal, and/or paleoclimatic shifts (Donoghue 2008; Donoghue and Edwards 2014; Jara-Arancio et al. 2014; Cardillo et al. 2017; Zizka et al. 2020).

However, phylogenetic patterns of biome affinities vary greatly among species throughout the tree of life. For some clades, most species only occupy individual biomes, and exhibit rare historical shifts between biomes that are often accompanied by other major evolutionary shifts in key ecological traits (Crisp et al. 2009; Gagnon et al. 2019). Other clades appear to be less constrained, with species that either are generalists across multiple major habitat types or appear to have repeatedly shifted among biomes (Holstein and Renner 2011; Cardillo et al. 2017; Ceccarelli et al. 2019; Magalhaes et al. 2019). Knowing what conditions promote the development of specialized versus generalized biome affinities would help us better understand community assembly, historical biogeography, adaptation, diversification, and extinction. Biologists are therefore especially interested in reconstructing when, where, and how species have adapted to different biomes throughout their evolutionary histories.

Motivated by this, biologists have applied phylogenetic models to a broad assortment of clades and biomes to reconstruct how species gain and lose affinities with different biomes over evolutionary timescales (Wiens and Graham 2005; Ree and Smith 2008; Goldberg et al. 2011; Landis et al. 2021a,b). These models typically assume that biome affinities are recorded as presence-absence characters that change along the branches of a time-calibrated phylogeny. Such reconstructions assume that a species is only considered to have an affinity with a biome if it is known (observed) to naturally occur within it. Yet, species do thrive in biomes where they have not historically occurred, as evidenced by invasive species that have recently colonized new biomes in new regions (Gallagher et al. 2010; Escobar et al. 2016; Banerjee et al. 2017; Wiens et al. 2019) and species that occupy anthropogenic habitats (Ellis and Ramankutty 2008; Martin et al. 2015; Winchell et al. 2020). Standard phylogenetic models fitted using biome occupancy data might capably infer which biomes ancestral species occupied, but they are not designed to infer the broader set of biomes ancestral species could have ecologically tolerated.

Ecologists interested in species ranges address this gap by distinguishing between the fundamental and the realized niche of a species. The fundamental niche is defined as the set of environmental factors in which populations grow at a net positive rate until reaching stable sizes, when disregarding biotic interactions and dispersal barriers within the environment (Grinnell 1917; Hutchinson 1957; Soberon and Peterson 2005; Sobeŕon 2007; Holt 2009; Peterson and Sobeŕon 2012; Jiménez and Sobeŕon 2022). The realized niche is defined as the set of environmental factors that the species stably occupies (Gaston 2003); typically it includes what remains of the fundamental niche after it has been truncated by biotic and dispersal factors (Sobeŕon 2007; Araújo and Luoto 2007; Pollock et al. 2014; Anderson et al. 2017). This distinction has proven useful when considering how ecological traits and species distributions may change over time, especially in the context of major environmental transition (Pearman et al. 2008; Schwartz 2012; Early and Sax 2014).

In an analogous manner, it should be important for phylogenetic models to distinguish among established biome affinities, enabled biome affinities, and non-affinities. A species has an established affinity for a biome if that biome contains environmentally suitable habitat patches that are actually occupied by the species as part of its realized niche. A species possesses an enabled affinity for a biome if that biome contains environmentally suitable habitat patches within the fundamental niche of the species, even though it does not occupy that biome owing to geographic or biotic inaccessibility. Lastly, a species has no affinity for a biome if the biome contains too little local habitat overlapping with the fundamental niche of the species for it to persist.

Distinguishing established from enabled biome shifts when using naive models that encode no such distinction is difficult (Sobeŕon and Townsend Peterson 2011; Schnitzler et al. 2012; Tingley et al. 2014; Wiens et al. 2019; Bates and Bertelsmeier 2021). For example, two scenarios might underlie the established biome shift of a species. In one case, a species might acquire a novel adaptation to occupy the new biome, implying both an enabled and established shift (Schnitzler et al. 2012). On the other hand, a species might be pre-adapted to a biome, and then simply establish itself therein, with no shift in enabled affinity required (Tingley et al. 2014).

What could we learn from a phylogenetic model that distinguishes between these biome shift scenarios? It may become possible to differentiate whether a species adapted to establish itself in a new biome, or whether it was pre-enabled and simply moved (Marazzi et al. 2012; Donoghue and Edwards 2014). Such a model could also help resolve how biome affinities changed during speciation, where several mechanisms can act differently on the realized versus fundamental niches. For example, ecological speciation may shift both enabled and established biome affinities, while geographical speciation driven by dispersal limitation may only shift established affinities (Mayr 1963; Endler 1977).

In this article we introduce a new phylogenetic method, *RFBS*, for modeling biome shifts among realized and fundamental niche states, such as established and enabled biome affinities. For a given set of biomes, a species can possess non-affinities, enabled affinities, or established affinities, as has been done in recent host repertoire models that distinguish between ”fundamental” and ”realized” host-use states (Braga et al. 2020). Our model is also inspired by the Dispersal-Extinction-Cladogenesis (*DEC*) model (Ree et al. 2005; Ree and Smith 2008), where discrete portions of a geographic range are gained and lost along phylogenetic branches, and may be asymmetrically inherited following cladogenesis. Under *RFBS*, species transition among affinity states according to anagenetic and cladogenetic events that reflect how phylogenetic lineages shift among biomes. We implement our model in a Bayesian framework using the programming language Julia. Using simulations, we validate our model, testing how accurately rates and ancestral states for biome affinities can be recovered even when data on enabled affinities is scarce. We also apply *RFBS* to the flowering plant clade *Viburnum*, which has moved and evolved among a variety of mesic forest biomes throughout its global radiation (Spriggs et al. 2015; Lens et al. 2016; Landis et al. 2021a,b).

## Methods

### Terminology

The description of *RFBS* as a model depends upon clear operational definitions for realized and fundamental niches, established and enabled biome affinities, and established and enabled biome shifts. We stress that our definitions of enabled and established biome affinities are related to, but distinct from, the definitions of fundamental and realized niches so as to account for the difference in scale between local niches and regional biomes. This section provides additional context for the brief definitions we gave in the introduction.

The niche of an individual species is a function of a vast set of abiotic and biotic factors and, thus, rarely aligns with all conditions within just one biome. Rather, species niches may span across many local habitat patches that exist within and among multiple biomes (Godsoe et al. 2017). In addition, biomes are defined at coarse regional scales, and are composed of multiple habitat types with differing environmental conditions (Williams et al. 2004; Godsoe et al. 2017; Mucina 2019).

Species-level variation in biome affinities are often observed at regional and phylogenetic scales, even though affinities are gained and lost at the scale of local populations. As stated above, a species has an established affinity for a biome if that biome contains environmentally suitable habitat patches that are actually occupied by the species as part of its realized niche. A species possesses an enabled affinity for a biome if that biome contains environmentally suitable habitat patches within the fundamental niche of the species, even though it does not occupy that biome owing to geographic or biotic inaccessibility. A species gains an enabled affinity with a novel biome without gaining an established affinity when its niche adapts to environmental conditions found both in the resident and the novel biome. For instance, many mesic biomes contain patches of rocky bare soil, which creates the opportunity for a species to adapt to the local conditions that resemble those of arid biomes. Circumstances such as this could allow a species to expand its fundamental niche to include arid conditions, thus gaining an enabled affinity to an arid biome elsewhere (Axelrod 1972; Lichter-Marck and Baldwin 2023). Later, the enabled species could secondarily gain an established affinity with an arid biome, e.g. following dispersal or paleoenvironmental changes in the distribution of biomes. Unsurprisingly, species with fundamental niches that cannot be realized within any present biomes are in a precarious position. They risk almost certain extinction (by definition) unless they establish an affinity within a present biome by expanding their fundamental and realized niches to encompass the necessary conditions for population-level persistence (Holt and Gaines 1992; Pulliam 2000; Holt 2009).

We define biome shifts as changes in enabled or established biome affinities. An enabled biome shift occurs when, for whatever underlying reason, a species gains or loses potential access to new environmental conditions through evolutionary changes in traits (Donoghue and Edwards 2014; Bates and Bertelsmeier 2021). An established biome shift occurs when a species actually gains or loses its presence in a biome (Tingley et al. 2014). Enabled and established biome shifts could occur in a step-wise fashion, with an enabled shift preceding an established one (as in the rocky outcrop example above). But, they could also occur more-or-less in unison, in a “lockstep” fashion (Edwards et al. 2017) – i.e., when a species is gradually exposed to novel conditions in a new biome, through repeated dispersal events or climatic transitions, and expands its fundamental niche as it comes to occupy the biome. During anagenesis (between speciation events), species shift between non-affinities and established affinities either directly or through an intermediate-enabled affinity (Braga et al. 2020). During cladogenesis (at speciation events), biome affinities are inherited either asymmetrically or equally between daughter lineages as a result of several biologically realistic event types. For example, geographic allopatric speciation might cause daughter lineages might “split” ancestral established affinities without dividing the enabled affinities. Ecological speciation, on the other hand, might cause daughter lineages to diverge both among established and enabled affinities during speciation.

### Model Description

This section defines *RFBS*, a phylogenetic model to characterize biome shifts among realized and fundamental niche states. With *RFBS*, the biome affinities of a species may take any of three states with respect to a particular biome: (i) a species may have no affinity or a non-affinity with a biome (encoded as “0”), meaning that there is no evidence or expectation that the species is suitably adapted to that biome; (ii) a species may have a enabled affinity with a biome (encoded as “1”), meaning that it is sufficiently adapted to survive and persist in a particular biome, regardless of whether or not the biome is within the species range; or (iii) a species may have an established affinity with a biome (encoded as “2”), meaning that the species is currently established within and is adapted to a given biome. This (0, 1, 2) affinity coding notation follows (Braga et al. 2020).

Regarding the relationships among the biome affinity states, we assume that all species that have an established affinity with a biome also implicitly have an enabled affinity with that biome. That is, established biome affinities are a subset of enabled biome affinities for a species at a given time. In addition, we assume that all species always have at least one established affinity, otherwise they would be extinct. As such, we do not allow species to have an established affinity for a biome without an enabled affinity for it, as they would fail to persist.

While it is possible that individuals of a species could enter habitat patches within novel biomes that are environmentally outside of the individuals’ fundamental niche and subsequently rapidly adapt within generations (Holt and Gaines 1992; Pulliam 2000; Holt 2009), we do not consider such instances as established affinities being gained before enabled affinities. At the macroevolutionary timescales we model, such events would appear as outcomes of a rapid “lock-step process”, wherein established and enabled affinities are gained simultaneously (Edwards et al. 2017). In the discussion, we expand on this concept, and argue that we must also consider lineage extinction to explicitly account for cases in which species occupy unsuitable biomes.

To proceed with the formal model definition, assume a system with three biomes (<u>t</u>ropical, <u>w</u>arm temperate, <u>c</u>old temperate). We represent the affinities of a species with each biome using an uppercase character for an established affinity (*T*, *W*, *C*), a lowercase character for an enabled affinity (*t*, *w*, *c*), and no character for a non-affinity. Thus the notation *Tw* Tw indicates that the species has an established affinity with the tropical forest biomes (*T*), enabled affinities with tropical and warm temperate biomes (*t* and *w*), and a non-affinity with cold temperate biomes. Assuming the biomes follow the order of their introduction, the affinities *Tw* are equivalently represented by the vector 210.

Our model assumes that biome affinities change differently during anagenesis (within a species over time) versus cladogenesis (when one species splits in two) (Fig. 1). Below, for simplicity, we provide examples of the anagenetic rate matrix, *Q*, the three-dimensional cladogenetic transition probability matrix, *P*, and all associated model parameters encoded for a two-biome system with only <u>h</u>ot and <u>c</u>old biomes, for which species evolve among five possible sets of biome affinities: (*H, C, Hc, hC, HC*). For a three-biome system with 19 states, see Supplement 1.

**Figure 1:**
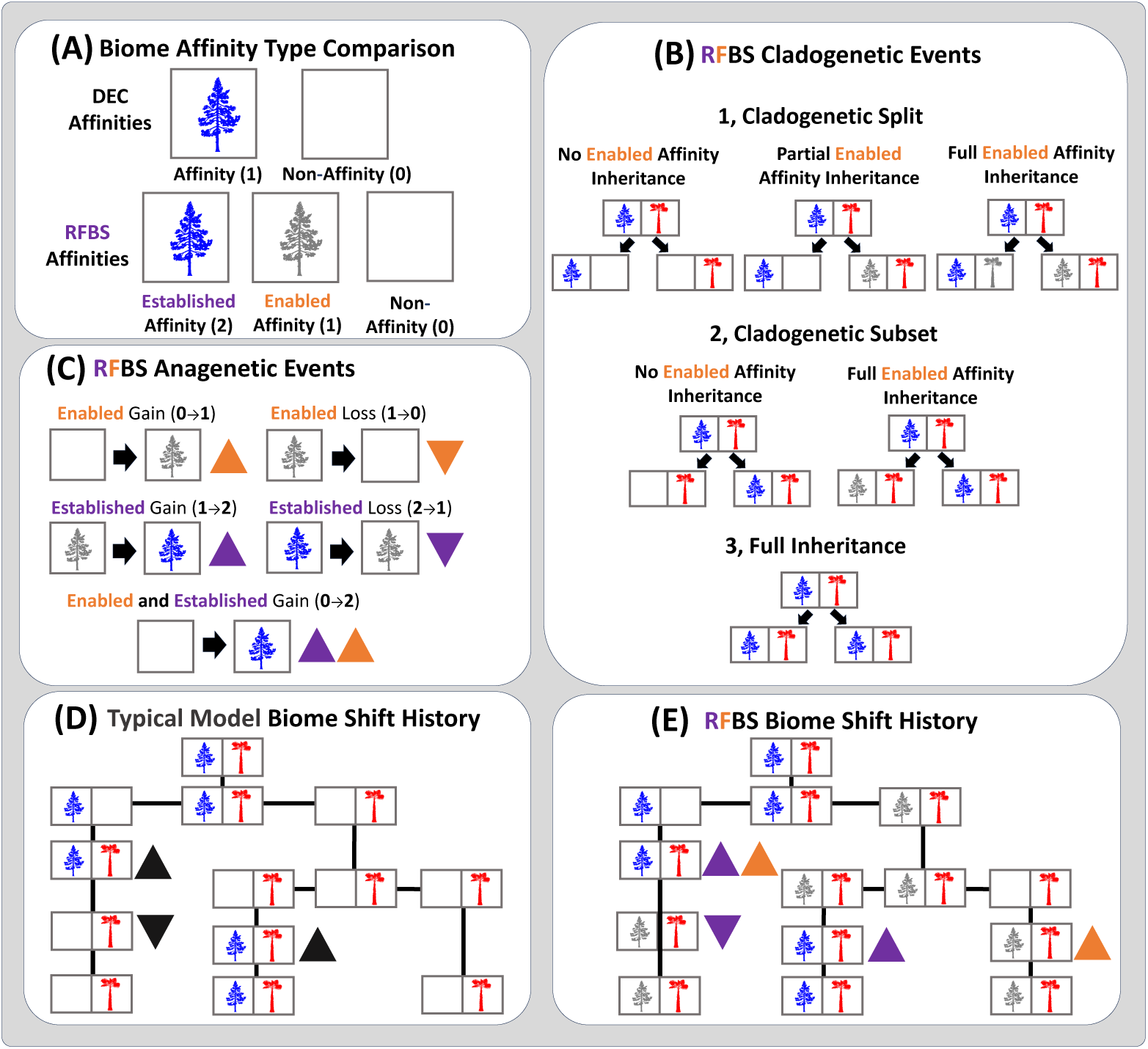
Cartoon of *RFBS* model events. This example shows an example character history from a typical biome shift model considering only biome established affinities and non-affinities, and a possible *RFBS* character history given the same established affinities on a toy phylogeny with three species and two biomes (cold: blue conifer in left cells; warm: red palm in right cells). A) A species has either an established affinity (purple text/arrows and colored tree), an enabled affinity (orange text/arrows and grayed-out tree), or a non-affinity, (blank tree) for each biome in the system. B) During cladogenesis, if a species has two or more biome affinities, they are inherited by daughter lineages in any of five ways, with the established affinity set either splitting or one daughter inheriting the full range while the other retains only a subset. Such cladogenetic events can act either only on the established affinities, with enabled affinities still being retained, or down to the enabled affinity, too. C) During anagenesis, species gain and lose affinities, as shown using a single cold temperate affinity. (D) Most models used to model biome shifts only consider biome occupancy, resulting in histories only for the gain and loss of established affinities. (E) *RFBS* may produce richer character histories with shifts in both established and enabled affinities.

During anagenesis, each event causes a species to gain or lose its established or enabled affinity with a single biome (Fig. 1a). An enabled affinity can only be lost if the species does not have an established affinity for that biome. We consider two different events where established affinities can be acquired in “step-wise” and “lock-step” scenarios. Consecutive single gain events represent “step-wise” process by which species first gain an enabled biome affinity (at rate *g*_0_*_→_*_1_) before gaining an established affinity by gaining access to the biome and persisting within it (at rate *g*_1_*_→_*_2_), as we expect under instances of biome preadaptation. Each double gain event allows for species to concomitantly adapt to biomes as they gain access to them in a rapid ”lock-step” process (Edwards et al. 2017), transitioning from a non-affinity to an established affinity (at rate *g*_0_*_→_*_2_). Species may lose established affinities (at rate *l*_2_*_→_*_1_) and enabled affinities (at rate *l*_1_*_→_*_0_) through separate events, though we forbid “double losses”, as it implies a scenario where species immediately evolve to lose an enabled affinity the moment it is no longer established within the biome.

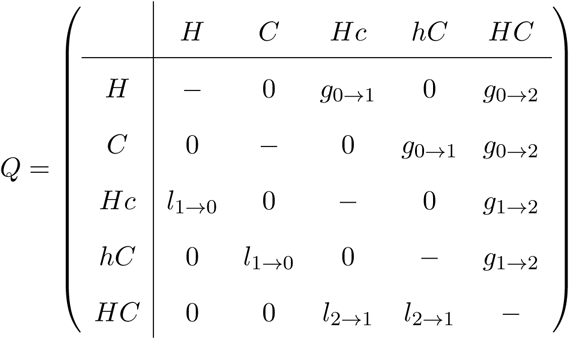

Cladogenetic events operate differently on established affinities, enabled affinities, and non-affinities (Fig. 1b). Established affinities are subsets of enabled affinities (as discussed above) that may also be truncated due to various non-evolutionary factors including competitive biotic interactions, trophic interactions, dispersal limitations, and many others. In a cladogenetic scenario involving primarily dispersal-limiting factors, such as allopatric speciation, daughter lineages do not necessarily identically inherit the established affinities of their immediate ancestor, but will likely conserve their enabled affinities (Wiens and Graham 2005). However, cladogenesis can also involve evolutionary factors acting on the fundamental niche, with different populations adapting locally to environmentally divergent habitat patches as a consequence of geographical speciation or a cause of ecological speciation (Rundell and Price 2009; Pyron et al. 2015; Rinćon-Barrado et al. 2021).

We model the probabilities of cladogenetic events as a three-dimensional probability matrix, *P*, with the size of each dimension corresponding to the size of the state space. The first dimension corresponds to each possible parental state, while the other two dimensions represent the two daughter states (arbitrarily, the rows correspond to left-daughter lineage states, and columns correspond to right-daughter lineage states). The probabilities of all possible pairs of daughter states descending from any parental state sum to one. Supplement 1 contains the full representation of the cladogenetic transition probability matrix for a three-biome system. Here we provide two specific examples that characterize the general behavior of cladogenetic event patterns and probabilities.

First, take the simplest case of a cladogenetic event where the parent lineage has only a single established affinity, and may have any number of enabled affinities. Affinities will always be identically inherited when the parent lineage has an established affinity for only one biome.

Assuming a parent lineage with an established affinity for a hot biome (*H*) and an enabled affinity for cold biome (*c*), *P_Hc_* is:

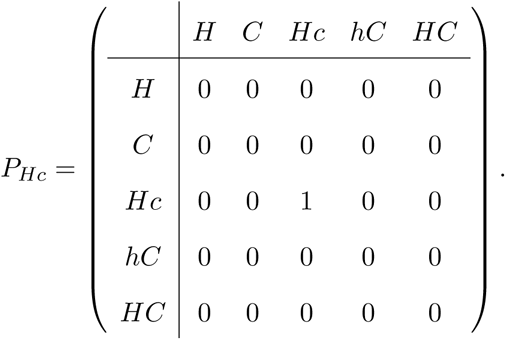

In the more complex case, a parental lineage may have multiple established biome affinities, plus any zero or more enabled affinities. Following cladogenesis, established and enabled affinities are inherited following equal-inheritance, subset-inheritance, or between-biome split-inheritance scenarios. We assume that the probabilities of the three scenarios are governed by the probabilities *p_e_* + *p_s_*+ *p_b_* = 1, respectively. To compute the cladogenetic transition probabilities, we additionally track the number of distinct cladogenetic outcomes under the three scenarios, *N_e,i_*, *N_s,i_*, *N_b,i_* for ancestral state *i*.

Under equal-inheritance cladogenesis, both daughter lineages inherit the full set of established parental biome affinities with probability 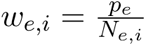 (note *N_e,i_* = 1 for all *i* in the models we explore here; see next paragraph). Subset-inheritance lets one daughter lineage inherit all established affinities in the parental state, while the second daughter lineage inherits only a subset of the parental affinities with probability 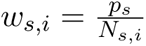. The between-biome split-inheritance scenario divides established parental affinities into non-empty sets for the two new daughter lineages, with each cladogenetic outcome having probability 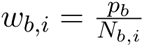. Both *N_s,i_* and *N_b,i_* can be expressed as a function of the number of established affinities in parental state *i* (*N_est,i_*) such that 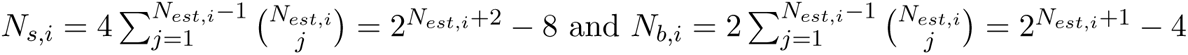, while *N_e,i_* = 1 for all *i*.

In both subset- and split-inheritance scenarios, the parental established affinities that are not inherited by daughter species may reduce in two ways. First, they may reduce down to non-affinities, implying a scenario of ecological divergence, where daughter lineages diverge in fundamental niche along one or more environmental gradients that constrain biome affinities.

Second, they may reduce down only to enabled affinities, implying a scenario where divergence is unrelated to species biome-affinities, such as geographic isolation.

As an example of these rules, the cladogenetic probability matrix for a parent lineage with two established biome affinities for hot and cold biomes, *P_HC_* is:

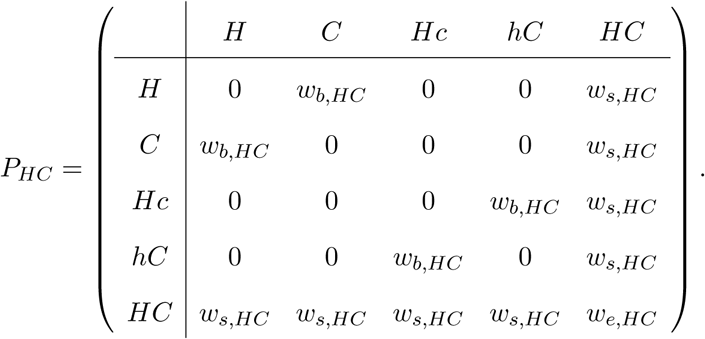

For our simulation and empirical analyses we chose these priors: *Q* matrix rate parameters *g*_0_*_→_*_1_*, g*_1_*_→_*_2_*, g*_0_*_→_*_2_*, l*_1_*_→_*_0_*, l*_2_*_→_*_1_ *∼* Exponential(*β*) where *β* is the scale parameter in units of time scaled to tree height, and *β*=0.5 in the empirical analyses and varies between 0.5 and 2.0 in simulation analyses. Cladogenetic scenario probabilities have priors *p_s_, p_b_ ∼* Uniform(0,1) with *p_e_* = 1 *− p_s_− p_b_*. We estimated model parameters using Markov chain Monte-Carlo (MCMC; Metropolis et al. (1953)) using the Metropolis-Hastings algorithm (Hastings 1970) for parameter updates. Each model parameter is updated with a standard univariate multiplier proposal, such that the newly proposed parameter value is updated as *x^′^* = *xe^λu^* where *λ* is a tuning parameter that controls the magnitude of the proposal and *u ∼* Unif(*−*0.5, 0.5).

### Resolving ambiguous biome affinity set states

*RFBS* models how biome affinities change along a phylogeny, yet all true biome affinities for any species are rarely known. Established biome affinities are perhaps the exception, which may be known with confidence through direct observation, occurrence data, and expert range maps. Assuming zero detection error, any species lacking an established affinity (an unestablished affinity) with a particular biome would have either a true enabled affinity or a true non-affinity.

Extrinsic information is needed to resolve the inherent ambiguity between enabled and non-affinities with biomes. Figure 2 illustrates the general strategies that we employ to constrain this ambiguity.

**Figure 2:**
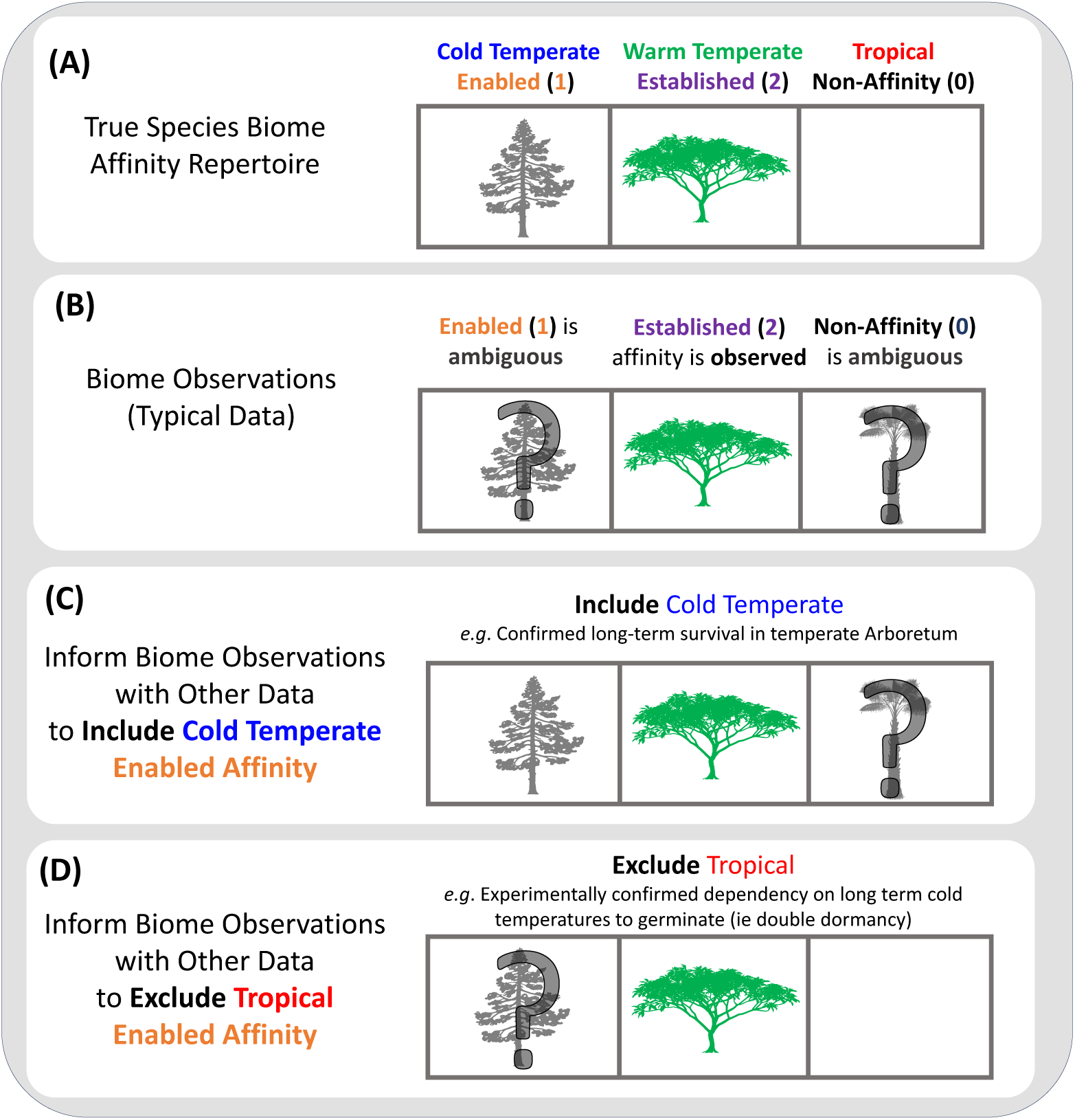
Cartoon depicting how to score enabled biome affinities. A species might have an enable, established, or non-affinity for each of three biomes: temperate (conifer), warm temperate (plumeria), and tropical (palm) forests. A) Each species will have a true set of biome affinities. B) True established biome affinities are typically scored appropriately as established affinities, however enabled and non-affinities are difficult to score as most species are rarely observed outside of their native biomes. In these instances, such affinities are left ambiguous. In example B, we would assign equal probability to states 020, 021, 120, and 121. Additional information can be used to include or exclude enabled affinities for some biomes. C) This example uses the occurrence of the plant in a cold temperate arboretum to include a cold temperate biome affinity. The possible states under this scenario are 120 and 121. D) This example uses experimental data showing the plant requires prolonged freezing periods to germinate (i.e. double dormancy). The possible states under this scenario are 020 and 120.

As the baseline, *RFBS* scores unestablished biome affinities as ambiguous for enabled (1) or non-affinity (0) with equal probability. In practice, we assign the same likelihood of observing each distinct affinity combination that is compatible with both the established affinities we do observe and any additional constraints imposed on biome affinities. For example, in Figure 2b, a species with a true established affinity for biome 1 and true enabled affinities for biomes 2 and 3 (state 211), would be ambiguously coded as having equal likelihoods for observing four states (211, 210, 201, and 200) of the 19 total states in a three-biome system. The worst-case scenario assumes complete biological ignorance, but may be improved through knowledge of species environmental tolerances or the climatic conditions among biomes, which may support including or excluding certain enabled affinities from the set of ambiguous states (Fig. 2c-d). Enabled affinities may be included when there is evidence of population persistence in conditions outside of the native range. Similarly, enabled affinities may be excluded based on observations of intolerance of specific abiotic conditions (e.g., a tropical species that is not frost tolerant almost certainly cannot survive in a cold temperate environment), experimental studies (e.g., failure of seed germination in the lab without exposure to prolonged cold temperatures), or knowledge on the adjacency of biomes along an abiotic gradient (e.g., a species with an established affinity for a tropical and for a cold temperate biome almost certainly must have an enabled affinity for intermediate warm temperate biomes).

The next sections explain how we design and apply strategies to constrain ambiguous affinities, and how we measure the sensitivity of *RFBS* to the presence of ambiguous biome affinities when constrained by different strategies.

### Simulation Experiments

We conducted simulation experiments to validate parameter estimation and ancestral state reconstruction accuracy when fit to data simulated under *RFBS* with varying levels of missing enabled affinity data and compare parameter and ancestral state estimation accuracy relative to *DEC* models when fit to data simulated under *RFBS*.

We first measured parameter estimation accuracy and precision under multiple tree size and prior distribution assignments. For each dataset, we sampled *RFBS* rate parameters from an exponential prior distribution with one of two scale parameters, 0.5, and 2.0, that represent prior treatments for slow and fast biome shift rates. Cladogenetic event probability parameters *p_s_*and

*p_b_* were each sampled under a uniform distribution (0.0 to 1.0) while setting *p_e_* = 1 *− p_s_ − p_b_*. In total, we fit *RFBS* to 100 independently simulated datasets under each combination of the two tree sizes (50 and 500 tips) and two prior scale parameter treatments (0.5 and 2.0), for a total of 400 simulated datasets. *RFBS* model parameters were fit using MCMC for 500,000 iterations, with proposal tuning parameters set to 1.5 for *Q* matrix parameters and 0.2 for the *P* cladogenetic probability matrix parameters. We evaluated estimation performance only on MCMC chains which exhibited sufficiently large effective sample sizes (*≥* 200). We evaluated posterior accuracy by measuring the frequency of data generating parameter values that fell within the 95% highest posterior density (HPD). Assuming coverage successes follow a binomial distribution, we expect that 90 to 99 of 100 (95 *±* 2 sd) HPDs will contain the true simulating value with 95% confidence.

Second, we tested *RFBS* under multiple scenarios where the data to resolve unestablished biome affinities is incomplete. To do so, we fit *RFBS* to six treatments of simulated data, where species affinities for unestablished biomes were either entirely ambiguous, partially ambiguous, or entirely confirmed. The entirely ambiguous treatment reflects the worst case scenario described above, where no tip states for enabled and non-affinities are resolved. The partially ambiguous treatments reflect the more common scenario where a biologist has an accurate dataset for established affinities with only limited information to discern enabled from non-affinities for some unestablished biomes for some species. The ambiguous affinity treatments started with the worst case of total ambiguity for unestablished biomes, and then resolved unestablished affinities to species at different percentages for different treatments. We resolved either 33% or 66% of ambiguous states that were not the true state, equivalent to removing at most either two ambiguous states or one ambiguous state, respectively. We applied both affinity resolution strategies to 25%, or 75% of tips in the tree, for a total of five ambiguous treatments for the missing data experiment. The entirely confirmed treatment resolved all enabled and non-affinities for all unestablished biomes, with each species having 100% probability for only a single state.

For each treatment we simulated and evaluated 100 datasets using the same protocol as the previous HPD coverage simulation experiment, but only over trees with 150 tips using an exponential prior with a scale parameter of 1.0 for all rate parameters.

Third, we tested *RFBS* accuracy when estimating ancestral states over phylogenetic nodes before and immediately following cladogenesis. We were also interested in how current approaches that do not consider enabled affinities may estimate ancestral established affinities or non-affinities over the tree from datasets simulated under *RFBS*. To investigate this question, we then compared the *RFBS* fit of each simulated dataset to the fit of a *DEC* model. Because *DEC* requires presence-absence data as input, we recoded *RFBS* affinities. Each established biome affinity (2) is coded as presence of affinity and each unestablished biome (0 or 1) is coded as absence of affinity. We fit both *RFBS* and *DEC* models to simulated (and, for *DEC*, recoded) datasets using the same MCMC algorithm as in previous experiments, but with an additional ancestral state sampling step. For all simulations we estimated ancestral state patterns for the entire phylogeny every 100 steps during the MCMC, generating posterior frequencies of states at the nodes prior to cladogenesis, and at the start of branches immediately following cladogenesis. These posterior state frequencies were then transformed into posterior frequencies for affinities for each biome, where each node has a probability for each possible affinity-type (established affinity, enabled affinity, or non-affinity) that sums to one for each biome. For each simulated dataset, we compared the posterior support for true established, enabled, and non-affinities for each biome at each node of the phylogeny. We summed the posterior support for the true affinities for each biome at each node, and then divided by the number of each affinity type to get the model accuracy for a given affinity, which was then averaged over all simulation replicates. For example, if we recovered posterior frequencies of 100% for all true established biome affinities over the phylogeny in each of our simulation we would have 100% established biome affinity simulation-wide accuracy. We compared the distribution of the percent of accurately estimated affinities for *RFBS* and *DEC* models relative to the random chance of sampling the true ancestral affinity in MCMC. To calculate the null probability of this chance, we followed the approach of Braga et. al. (2020), calculating the mean posterior probability for each true established and non-biome affinity across the 50 replicate posterior distributions from the same simulation treatment. This step was important for comparing ancestral biome affinity accuracy between methods since it is easier to draw established biome affinities by chance under *DEC* (one out of two possible affinity states) than under *RFBS* (one out of three possible affinity states). To assess performance, we were most interested in the ancestral affinity accuracy relative to the baseline of random chance.

### Empirical Study: V iburnum Biome Shifts

To characterize model performance under a real evolutionary scenario, we applied *RFBS* to an empirical dataset to compare transition rate and ancestral affinity estimates under various model and data treatment assumptions. We selected *Viburnum* (Adoxaceae, Dipsacales), a flowering plant clade of approximately 165 living species that likely originated near the start of the Cenozoic (Spriggs et al. 2015; Landis et al. 2021b). *Viburnum* species are primarily distributed through Asia, Europe, and the Americas, and variously inhabit cold temperate, warm temperate, cloud, and tropical mesic forest biomes. Previous phylogenetic studies have inferred that *Viburnum* experienced a dynamic history of geographic movement and ecological shifts, with frequent instances of sister lineages occupying different biomes (Donoghue and Sanderson 2015; Lens et al. 2016; Edwards et al. 2017; Landis et al. 2021b,a). An early analysis using only extant taxa and simple models with just two biomes (tropical and temperate) inferred a tropical origin for *Viburnum* (Spriggs et al. 2015). Subsequent work by Lens et al. (2016), using three biomes, instead supports either a warm or a cold temperate origin, with a slight preference for colder forests. A third round of analyses that included extant and fossil taxa (Landis et al. 2021b), or that modeled spatiotemporal variation in biome availability and connectivity (Landis et al. 2021a), both favored warm temperate origins for *Viburnum*. All of these studies used phylogenetic trees and established biome affinity data to trace the ecological history of the clade, which suggests that the *RFBS* approach could provide another complementary perspective on *Viburnum* evolution and biogeography.

Our *RFBS* analysis used a maximum clade credibility (MCC) phylogeny of *Viburnum* that includes 163 extant species and 5 fossil pollen taxa, previously generated by Landis et al. (2021b). The main results of that study were based on a joint phylogenetic and biogeographic analysis of extant and fossil taxa. That study also generated a supplemental MCC tree under the same procedure, except biogeographic and biome affinity states were masked for its fossil taxa during inference. We used this “masked” MCC tree rather than the one from the main results so as not to double-count the influence of fossil biome states upon our *RFBS* results.

Next, we constructed a matrix of established biome affinities for *Viburnum* by modifying the Landis et al. (2021b) dataset. Methodological limitations in the original study only allowed each species to occupy a single biome at a time, so it scored species in multiple biomes as ambiguous across all occupied biomes. *RFBS* is not limited in this way, so each species was scored as established for each biome it occupied. Cloud forest affinities, shared by members of the neotropical *Oreinotinus* radiation (Donoghue et al. 2022), were recoded here as warm temperate. Species in either of these biomes can experience relatively similar temperature ranges, though on different timescales, but neither is subjected to the prolonged periods of freezing temperatures that characterize cold temperate forests.

As enabled biome affinities for many species are poorly understood, we fit *RFBS* to the empirical dataset while leaving tip states ambiguous for all unestablished affinities, per the worst-case scenario in simulation experiment 2. In this case a species with an established tropical affinity and enabled affinities for warm and cold temperate biomes (true state of 211), would be coded as ambiguous across the four states 211, 210, 201, and 200.

Combining what is known about *Viburnum* natural history, ecophysiology, development, and horticulture, we developed a biologically informed set of criteria to explicitly include and exclude enabled affinities for the *Viburnum* species for which we were able to obtain relevant data. We included enabled cold temperate affinities for 4 *Viburnum* species based on records of long-term survival under non-native continental conditions as defined by the Kóppen-Geiger Climate classification, where the average temperate of the coldest month must be below freezing. Evidence for long-term survival came from records of long-term persistence within gardens or arboreta from the Botanical Garden Conservation International Plant Search Database recorded in 2015 or earlier. We excluded enabled tropical affinities for 18 *Viburnum* species based on records of the failure to survive in arboreta or the failure of seed germination under constant, warm climatic conditions. With respect to seed germination, we note that some *Viburnum* species native to cold forests have evolved a germination strategy that requires a prolonged period of cold (freezing or near freezing) temperatures to complete the process, which then extends under natural conditions through one or more winter periods. We reason that species with this derived form of “deep simple epicotyl morpho-physiological dormancy” (Baskin et al., 2006) would be unable to establish in warm and tropical forests that never experience sufficiently prolonged cold periods. In some analyses, we scored only the 17 species known from germination experiments to have this germination requirement, while in others, we extended this scoring to their close relatives that also occupy cold forests. We also made use of data on leafing habit, where we have nearly complete information (Edwards et al., 2017). In this case, we reasoned, based in part on records from arboreta, that evergreen viburnums are unlikely to survive in climates with prolonged freezing. Supplement 2 provides an expanded explanation (with references) of each line of evidence used in any of our analyses to include or exclude biome affinities. Finally, we excluded some biome affinity set states based on climatic adjacency. Given that cold temperate, warm temperate, and tropical biomes are arrayed along a thermal gradient, we required species with cold temperate and tropical affinities to also minimally have an enabled affinity with warm temperate biomes.

We fit *RFBS* to *Viburnum* 18 times against all permutations of included affinity, excluded affinity, and climatic gradient-informed affinities, under bold and conservative constraints for inclusion and exclusion factors, allowing us to characterize how rate and ancestral state estimates differed with and without each source of enabled affinity data.

## Results

### Simulated results

Simulation experiments show that *RFBS* behaves as intended under various tree sizes and rate prior conditions. Bayesian coverage is appropriate in all treatments (Figure 3). Accuracy and precision decreased with slower rates and smaller tree size, with the smallest tree treatment (50 tips) with low (0.5) rates being the least accurate (Figure 3b). Ancestral state estimation accuracy was also high in most trials (*>*90% accuracy on average); however, *RFBS* inference of enabled affinities came at a small cost in accuracy when inferring ancestral non-affinities when compared to the fit of an enabled affinity-naive *DEC* -like model on the same data (Fig. 4, Fig. S1). When compared to a *DEC* -like equivalent, *RFBS* recovered affinities as accurately, if not better, under the slower rate regime (Figs. 4A and 4C). In the higher rate regime, non-affinity accuracy drops to a lower accuracy than the simpler *DEC* -like model (Figs. 4B and 4D, Figs. S1B and S1D). This accuracy increased with the larger tree size (500 tips), recovering more comparable performance (Figs. 4C and 4D, Figs. S1C and S1D). The lower accuracy of non-affinity estimation compared to the simpler *DEC* -like model is likely in part due to differences in the random probability of accurately recovering true non-affinities between the two models, which is much higher in the *DEC* -like model.

**Figure 3:**
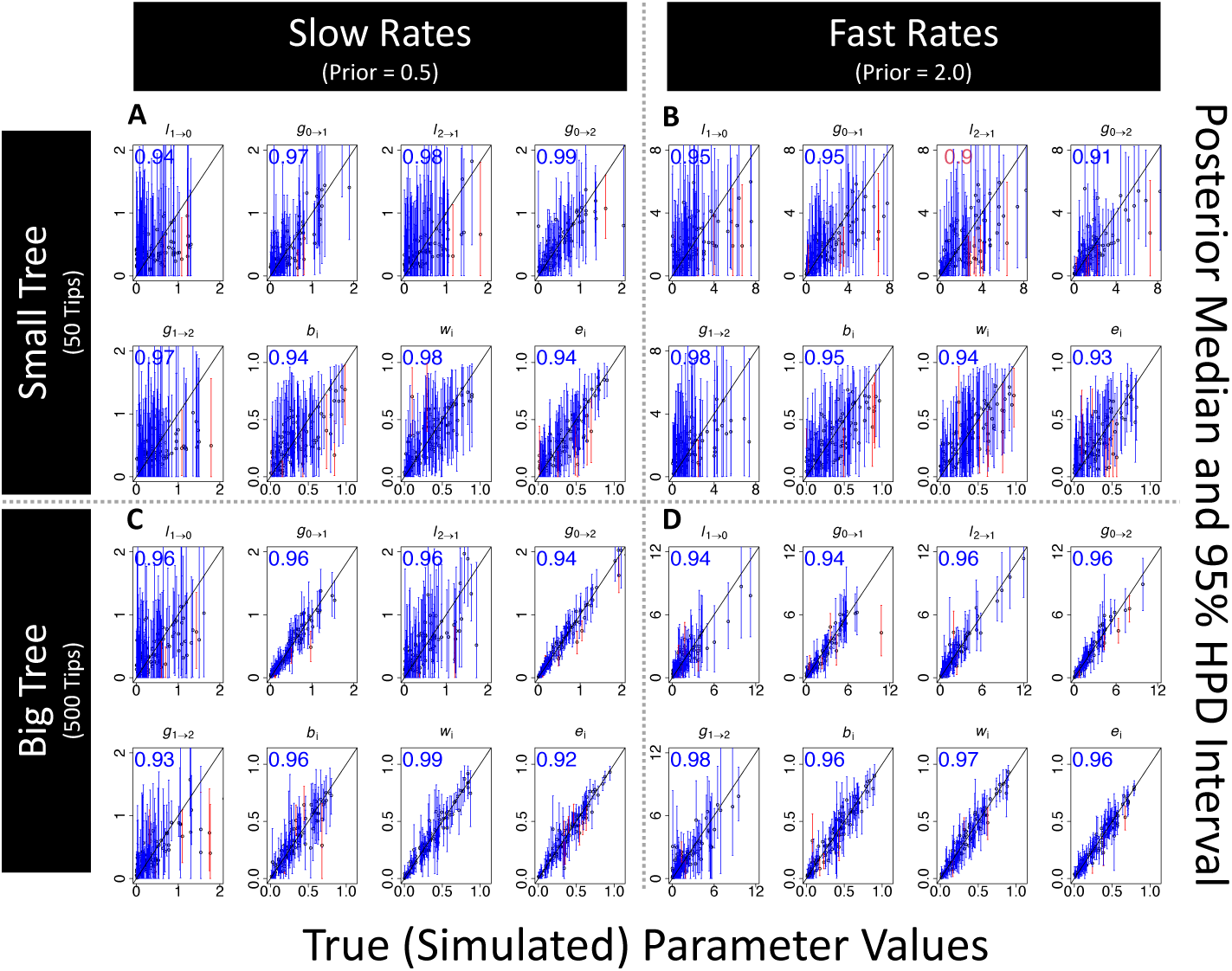
Rate parameter coverage simulation experiment assessing parameter estimation accuracy for a 8-parameter *RFBS* model with five free parameters for rates and constrained three parameters for the probability of cladogenetic splitting events. Four simulation treatments of 100 datasets are shown for two treatments of tree size (Small Tree= 50 tips, Large Tree=500 tips) and mean biome shift rates(Slow=0.5, Fast=2.0). Points plotted on each graph are the posterior medians on the y axis and true parameter value on the x axis, with the arrows from each point showing the 50 percent HPD. Red lines indicate estimates where the true data generating parameters were not in the 50 percent HPD and blue indicates estimates that were. The number on the upper left corner of each plot indicates the percentage of 100 simulation trials where the true parameters fell within the 50 percent HPD. Blue for this number indicate the percent is within two standard deviations of the expected mean given the number of samples and percent expected. A) Small tree and slow rate treatment, B) Small tree and fast rate treatment, C) Large tree and slow rate treatment, D) Large tree and fast rate treatment.

**Figure 4:**
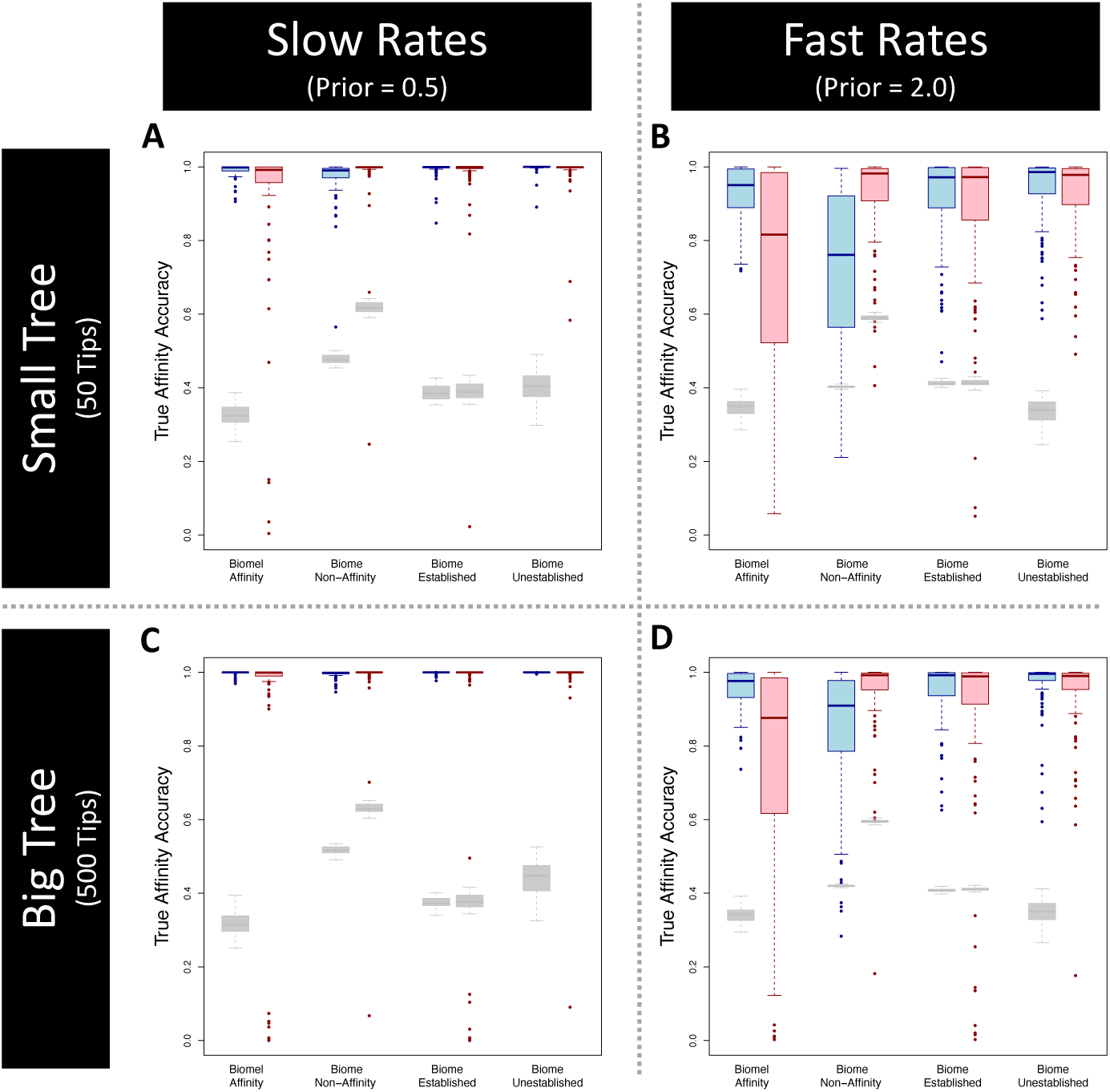
Ancestral affinity accuracy for *RFBS* (blue boxes) and *DEC* (red boxes) simulations. Box- plots represent the distribution of support for the true affinity for each biome at each node/corner of the ancestral state reconstructions. Gray boxplots represent the distribution of null probabilities of sampling the true ancestral affinity by chance, as done by Braga et. al. (2020). Columns of each plot include support for biome affinity (established/enabled) non-affinity, persistent biome occupancy (established affinity), biome non-occupancy (enabled/non-affinity). Four simulation treatments of 100 datasets are shown for two treatments of tree size (Small Tree= 50 tips, Large Tree=500 tips) and exponential prior scaling parameter(Slow=0.5, Fast=2.0). A) Small tree and slow rate treatment, B) Small tree and fast rate treatment, C) Large tree and slow rate treatment, D) Large tree and fast rate treatment.

Parameter estimation and ancestral biome affinity estimation accuracy decreased as ambiguity in tip states among unestablished biomes increased (Figs. S2 and S3). Higher amounts of unestablished biome ambiguity led to less precise and less accurate process rate estimates that tended towards underestimation, especially for rates of enabled affinity shifts (*g*_0_*_→_*_1_*, g*_0_*_→_*_2_*, l*_1_*_→_*_0_) (Fig. S2). In the worst case, where unestablished biomes at each tip were fully ambiguous among enabled and non-affinities, estimated rates of enabled affinity gain and loss nearly matched the prior, suggesting that at least some information resolving unestablished affinities is needed to infer those rates. Eliminating even just one state from the set of possible ambiguous states for 25% of species greatly improved rate inference. Estimation accuracy for ancestral biome affinities dropped with greater unestablished biome affinity uncertainty. Ancestral estimates for enabled and non-affinities markedly declined in accuracy and precision with more ambiguous unestablished affinities among sampled taxa, with the most significant decrease obtained when fewer than two ambiguous states were removed from the ambiguous state set for under 75% of tips. However, as expected, *RFBS* recovered biome affinities more accurately than the enabled-affinity-naive *DEC* -like model across all treatments. Even when the unestablished biome affinities were unresolved and entirely ambiguous, the ability of *RFBS* to infer ancestral patterns of biome occupancy vs. non-occupancy (established vs. unestablished affinity) remained very accurate across all treatments.

### Empirical results (Viburnum)

In our entirely ambiguous dataset, with no data that included or excluded ambiguous tip states, posterior rate estimates closely resembled the underlying priors (gray distributions in Fig. 5). When we added our biome climatic gradient rules (“A”), included enabled affinities (“I”), and excluded enabled affinities (“E”) for unestablished biomes, we found that the enabled biome affinity gain rates (*g*_0_*_→_*_1_; Fig. 5AC) decreased while enabled loss rates increased (*g*_1_*_→_*_0_; Fig. 5B). For all treatments informing unestablished affinities (colored distributions in Fig. 5), rates of enabled biome affinity gain (*g*_0_*_→_*_1_) were slower than rates of established biome affinity gain (*g*_1_*_→_*_2_) by at least an order of magnitude (Fig. 5A,C) while rates of established (*g*_2_*_→_*_1_) and enabled affinity loss (*g*_1_*_→_*_0_) remained comparable (Fig. 5B,D). With respect to different treatment strategies, gain and loss events entering the enabled state remained relatively constant (Fig. 5A,D), whereas gain and loss events leaving the enabled state fluctuated substantially (Fig. 5B,C). Rate estimates from runs that excluded unlikely enabled affinities (“E” treatment) formed a distinct cluster from the other treatments for most parameters, but most notably with respect to established single gain/loss rates and enabled affinity loss rates. The treatement that included unestablished enabled affinities (“I”) decreased both enabled affinity gain and loss rates. Estimates that both included and excluded enabled affinities as tip state information (“I” and “E”) tended to fall in between more extreme estimates produced when only included or excluded affinity information was used. Coding tip states to include particular enabled affinities (“I”) decreased the probability of subset inheritance of affinities during cladogenesis (*s*; Fig. 5G); otherwise, treatments to code enabled tip states had a weak impact on cladogenetic parameter estimates (*b, s, e*; Fig. 5F,G,H). Using bold (“Ib”) versus conservative (“Ic”) included affinities resulted in a substantial decrease in rates of gain and loss anagentic events leaving the enabled state (Fig. 5B,C) and decreased probabilities for subset inheritance cladogenetic events (Fig. 5G). Using bold (“Eb”) versus conservative (“Ec”) excluded affinities produced similar parameter estimates throughout.

**Figure 5:**
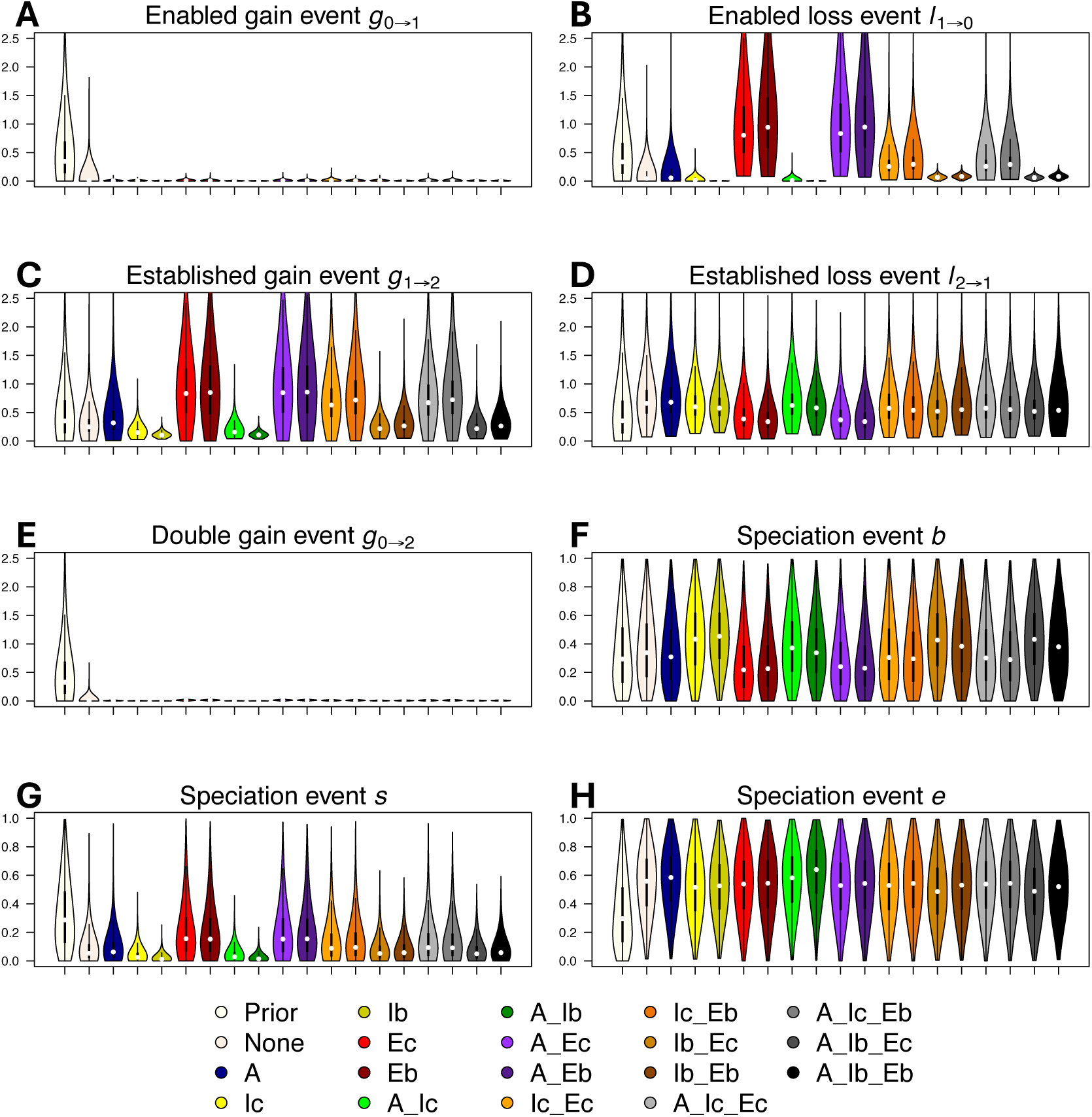
Posterior distributions of *RFBS* parameters for *Viburnum*. Rate parameters are *g* for affinity gain and *l* for affinity loss, which are subscripted by the transition type: 2=established affinity, 1=enabled affinity, an 0=non-affinity. Cladogenetic probability parameters are *e* for equal-affinity inheritance, *s* for subset-affinity inheritance, and *b* for between-affinity split inheritance. Colors indicate different treatments where different criteria informed unestablished biome affinities, including 1) a climatic adjacency rule (A) where species could not have a non-affinity (established or enabled) for warm temperate if there was an affinity for cold temperate and tropical; 2) included enabled affinities (I) from long term persistence in non-native biomes (via arboretums) under conservative (c) and bold (b) expert-based assessments; 3) excluded enabled affinities (E) based on evidence of non-affinity under conservative (c) and bold (b) expert-based assessments. Violin plots show parameter estimates, with circles indicating posterior medians, and lines indicating 25th and 75th percentile interquartile ranges. We show here the strongest exclusion criteria from our set of exclusion criteria (conservative/bold germination and bold leafing habit) which includes expert taxonomic interpolation to exclude the maximum possible set of enabled affinities.

Our ancestral biome affinity reconstructions are shown in Figures 6 and S5, informed by what we deem a conservative subset of our collected data on non-established biome affinities. This set of information includes freezing-dependent germination types, leafing habits, and USDA hardiness zone data, phylogenetic extrapolations from those data, and validated herbarium record data (Supplement 2; see Figs. S6, S7, S8, and S9 for reconstructions under using alternative sets of constraints). In agreement with Landis et al. (2021a, b), our *RFBS* analyses find that *Viburnum* likely originated in a warm temperate biome. Furthermore, we infer that the *Viburnum* ancestor likely had an enabled and established affinity for warm temperate biomes, and an enabled affinity for cold temperate biomes, with some support, albeit less likely, for an enabled affinity for tropical forests. This warm/cold temperate enabled affinity persisted as the most likely set of enabled affinities for most of the major clades, with lineages either retaining both affinities or losing an enabled cold or warm temperate affinity. We find no unequivocal instances of enabled cold temperate or warm temperate biome affinities being regained when lost. Tropical biome affinities were gained repeatedly, in several clades, all of which soon went on to occupy tropical biomes. Though it was not clear from the ancestral state reconstructions how frequently enabled and established affinities for tropical biomes were gained simultaneously or sequentially, we infer a far lower rate of “lock-step” events compared to “step-wise” gains of established affinities (Fig. 5).

**Figure 6:**
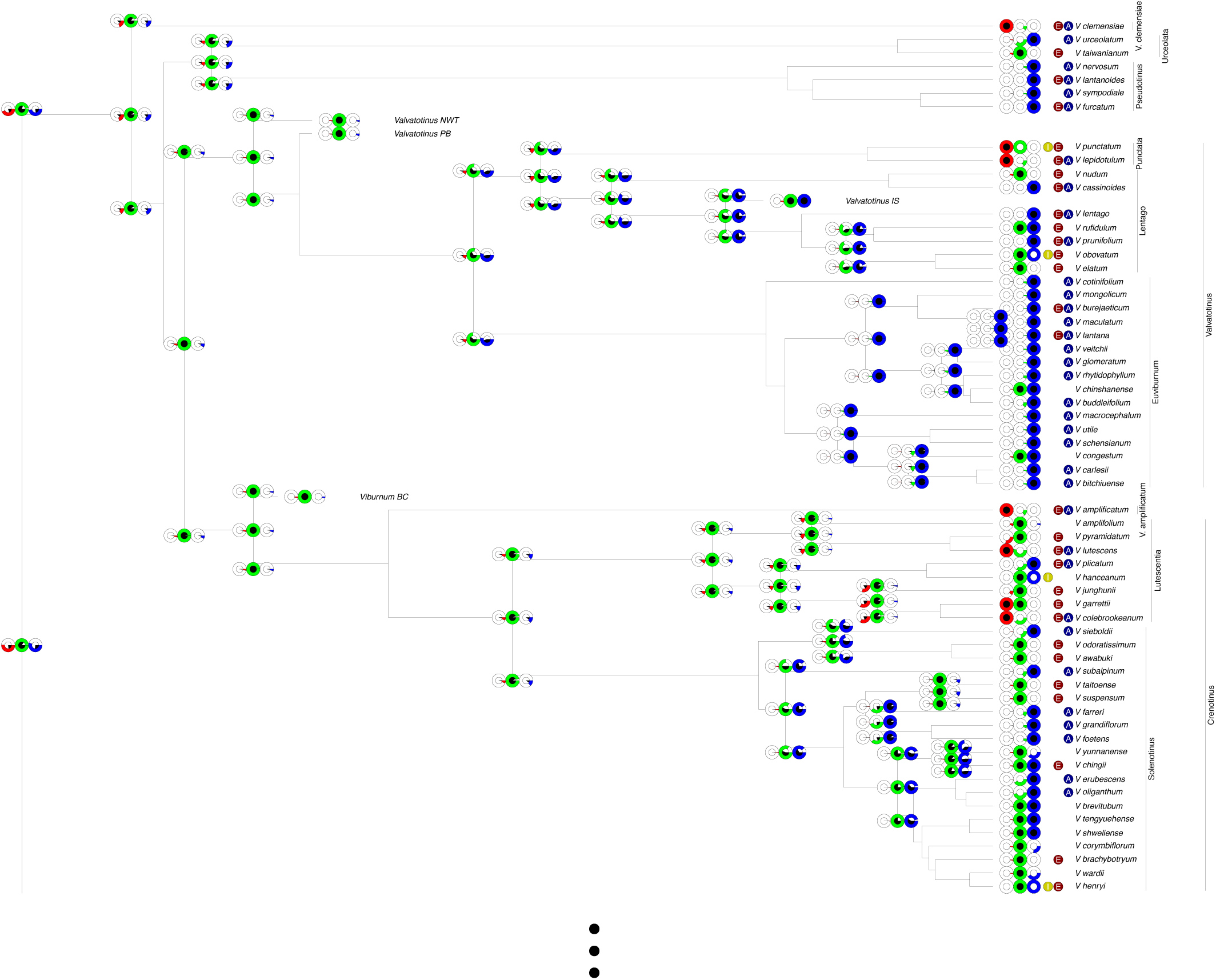
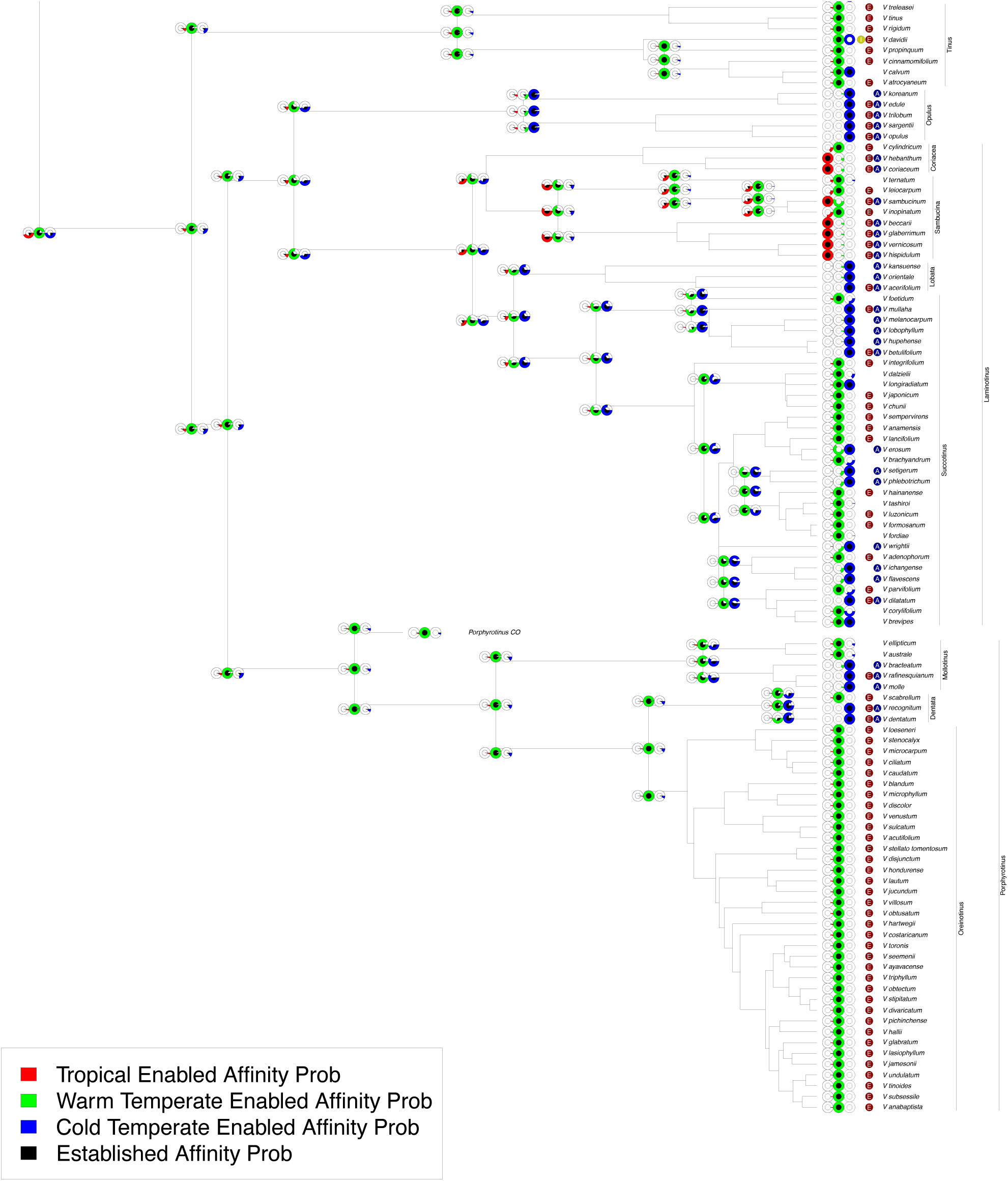
Ancestral state reconstruction of *Viburnum* biome affinities from *RFBS* using conservative included and excluded enabled affinity tip data. Note, this is a two-page figure. Node and corner pie chart triplets represent the probability for each biome established/enabled affinity given the probabilities for each state. Outer colored circles represent posterior probability support for enabled affinities (red: tropical, green: warm temperate, blue: cold temperate) while black inner circles represent posterior probability for established biome affinities. Colored circles between affinity probability pie charts and tip labels denote information used to reduce tip ambiguity. Dark red “E”: enabled biome affinity excluded; dark yellow “I”: established biome affinity included; dark blue “A”: climatic adjacency rule applied (i.e., a species cannot have an affinity for cold temperate and tropical biomes with no affinity for intermediary warm temperate biomes).

Ancestral state reconstructions did not change substantially after incorporating any of the enabled affinity data to our tip state probabilities (included affinities, excluded affinities, or climatic gradient) (Figs. 6 and S5-S7), with both supporting a warm temperate ancestor and predominantly warm temperate and cold temperate enabled affinity history. However, the inference of most model parameters responded to additional enabled affinity data treatments, with increasing rates of established affinity gain and loss when using more enabled affinity data with *RFBS*. The entirely ambiguous dataset analyses generally supported lower rates of established and enabled gain and loss, with bimodal posterior distributions for established affinity gain rates. Cladogenetic split probability also shifted from a higher rate of cladogenetic splitting to a probability closer to 0.5 with more enabled affinity data, particularly included enabled affinities

We introduce a new phylogenetic model, *RFBS*, to characterize how the enabled and established biome affinities of species shift through anagenetic and cladogenetic events (Fig. 1). Our approach to modeling biome shifts has multiple features absent from current approaches that only consider biome occupancy vs. non-occupancy. First, *RFBS* provides a means to distinguish between observations of native biome occupancy versus biome suitability, as recent biological invasions and ecological experiments suggest suitability is broader than occupancy for many species (Soberon and Peterson 2005; Hoogenboom and Connolly 2009; Gallagher et al. 2010; Jiménez et al. 2019; Bates and Bertelsmeier 2021). Under current models, biome affinities for each species are scored under a binary system as either having an affinity or not, with no distinction between unestablished but enabled affinities and complete non-affinities. Our results demonstrate that we can explicitly model this important distinction and that how biome affinity data is encoded impacts model performance. Second, the performance of *RFBS* is improved when incorporating information about enabled affinities from the wealth of observational, experimental, and natural history data that are available (Fig. 2). Traditional biome shift models do not readily digest this precious source of biological knowledge. Third, by distinguishing enabled versus biome shifts, we can test hypotheses that require the separation of evolutionary processes driving adaptation to biomes versus non-evolutionary ecological and biogeographic processes driving biome occupancy both during anagenesis and cladogenesis, such as how biome conservatism relates to these two factors (Wiens and Graham 2005; Cooper et al. 2010; Crisp and Cook 2012; Pyron et al. 2015), as we did with *Viburnum* biome affinities.

Our study assesses how *RFBS* performs under various realistic analysis conditions. One real-world limitation is that fundamental niche data is often sparse and difficult to collect. Though modeling niche evolution only using truncated realized niche data is a challenge for most models (Soberon and Peterson 2005; Saupe et al. 2018; Jiménez et al. 2019), *RFBS* might also misbehave when provided incomplete enabled biome affinity data. Our simulations show that *RFBS* can accurately infer model parameters and ancestral states, even when data on enabled affinities are sparse but not entirely missing (Figs. 3 and 4). *RFBS* also accurately infers biomes shift parameters in most datasets when percentages of ambiguous biome affinities for unestablished biomes (i.e., biomes where a species does not have an established affinity) is high. Parameter estimation accuracy worsens when all unestablished biomes for all species have ambiguous affinities (Fig. 4a). *RFBS* was reasonably accurate in inferring ancestral non-affinities for most datasets so long as some enabled affinity tip state data was provided. Overall, this suggests our model is most suitable for application to datasets where any knowledge of enabled affinities is known, including from observation and experimental sources, or structuring of biomes along a specific climatic gradient, such that some biome affinity sets for some species are safely eliminated. We suspect *RFBS* is most useful when applied to clades that possess evidence of affinities or non-affinities to unestablished biomes for at least 30% of terminal taxa. This modest fraction of data could be obtained for many clades using strategies similar to what we employed for *Viburnum*.

Our empirical analyses demonstrate *RFBS* can recover dynamic histories of enabled biome shifts when provided with a realistically attainable amount of unestablished biome affinity data (4 included affinities, 144 excluded affinities, and 63 species subject to the climatic adjacency rule). In *Viburnum*, *RFBS* recovers a phylogenetic pattern of deeply conserved enabled affinities in some clades and enabled affinity divergence in others (Figs. 6 and S5-S). Both warm and cold temperate enabled affinities are conserved throughout much of the tree, where clades with high proportions of tropical biome affinities are exceptions. Likewise, our estimates of enabled affinity gain and loss rates are two to three times slower than the rates of established affinity gain/loss, suggesting that established affinities are more ephemeral than enabled affinities in *Viburnum* (Fig. 5). We recover strong support for a common ancestor with warm temperate and cold temperate enabled affinity and weaker support for a tropical biome affinity. Our inference of an ancestral warm temperate established affinity aligns with (and provides further support for) the most recent studies of biome shifting in *Viburnum* using fossil data (Landis et al. 2021b) and the distribution of biomes in the past (Landis et al. 2021a). Our findings are at odds with Spriggs et al. (2105) who inferred that *Viburnum* originated in tropical forests, and also with Lens et al. (2016), who slightly favored an origin in cold temperate forests. We note, however, that our findings are broadly consistent with the hypothesis of Lens et al. (2016) that *Viburnum* retained scalariform vessel perforation plates as it transitioned into colder forests. Importantly, our *RFBS* analyses add a new dimension in suggesting that even from an early stage *Viburnum* species may have been enabled to occupy colder temperate forests and that such transitions may have therefore been relatively easy in these plants. Again, this aligns well with previous inferences of multiple (likely more than a dozen) such transitions as *Viburnum* lineages adapted to the global spread of colder climates from the Oligocene onward and evolved rounder leaves with toothed leaf margins (Schmerler et al. 2012; Spriggs et al. 2018) and the deciduous habit (Edwards et al. 2017). This high degree of consilience is gratifying, but we look forward to future studies that incorporate even more relevant historical and morphological data in a larger joint inference framework.

While our *RFBS* analyses generally align well with other recent analyses of biome shifts in *Viburnum*, we emphasize that they offer new insights and alternative interpretations in some critical places. As a concrete example, our analyses clear up ambiguity surrounding the origin of the neotropical cloud forest clade (*Oreinotinus*; Donoghue et al. (2022). In Landis et al. (2021a), this is uncertain, leaving open the possibility that cloud forests were first occupied by species from eastern North America that were already adapted to cold forests. Our analyses that incorporate information on seed germination rule this out, and instead indicate that cloud forests were occupied by a lineage that had long retained an affinity for warm forests. And, again in contrast to Landis et al. (2021a), the related *Dentata* and *Mollotinus* clades are here inferred to have had warm temperate and cold temperate ancestries, respectively. In turn, all of these findings have an important bearing on our interpretation of the direction of evolution in leaf traits.

With *Viburnum* and the mesic forest biomes that its species inhabit, we incorporated several specific lines of observational and experimental data (records from botanical gardens, data on seed germination requirements, etc.) to resolve non-established affinities as being either enabled or non-affinities. We expect that such strategies can be tailored to suit studies involving different clades and biomes. In *RFBS*, enabled biome affinities can be informed by relatively few traits measured across many species, or informed by many traits and rare observations that individually inform enabled affinities for just a few species. That said, such data are expected to sometimes be inconclusive or in conflict. To better address missing data in the future, *RFBS* can be extended to jointly infer the history of biome shifts with the species probability of suitability for each unobserved biome affinity, using correlative models where the affinity to each unestablished biome is a probability informed by a set of climatic, ecological, and morphological predictors (Groves and Castri 1991; Bartlett et al. 2012; Meloro et al. 2013; Hipsley and Müller 2017; Jiménez and Sobeŕon 2022).

While we developed *RFBS* with biome shifts in mind, the method is applicable for modeling any discrete realized-fundamental or enabled-established niche space with asymmetric cladogenesis. One interesting application of our approach would be including anthropogenic habitat types like urban environments (Martin et al. 2015; Winchell et al. 2020; Marsden et al. 2023) to predict which species have enabled affinities to urban settings and when such unestablished affinities are gained in the course of phylogeny. Other possible examples include dietary niche (Shipley et al. 2009; Brandl et al. 2015; Grundler and Rabosky 2020; Pansu et al. 2022), parasite host repertoires (Braga et al. 2020; Braga and Janz 2021), and zoonotic transmission (Baele et al. 2017; Fountain-Jones et al. 2018). In these cases, the discrete niche space is measured in term of interactions among particular species. For dietary niche, shifts in prey clade or trophic guild may be linked with adaptive phenotypic shifts or ecological speciation events (Felice et al. 2019; Román-Palacios et al. 2019). By contrast, many species are generalists with broad dietary niches yet may only consume species from a narrower realized range of species (Shipley et al. 2009; Pansu et al. 2022). Similarly, host repertoire shifts and parasite speciation are often linked, and are thought to be an important driver of diversification in many parasite clades (de Vienne et al. 2013; Hardy 2017). Current methods that consider fundamental versus realized host affinities are limited to scenarios of equal inheritance of host repertoires upon cladogenesis (Braga et al. 2020). Lastly, epidemiologists are concerned with the spread of pathogens, both in terms of its environmental niche and across species-level boundaries (particularly into livestock and/or humans) (Lloyd-Smith 2013; Guth et al. 2020; Latinne et al.

2020). Currently, phylodynamic approaches infer how niche variables impact host switching and transmission rates (Baele et al. 2017; Fountain-Jones et al. 2018), but typically only in terms of the observed environments of the pathogen and hosts rather than the range of conditions and hosts they may have an affinity to. In all of these cases, we suspect the modeling features of *RFBS* can be applied to better understand how fundamental and realized ecological relationships are inherited following cladogenesis.

That said, as noted in Braga et al. (2020), models such as *RFBS*, *DEC*, and the Braga host-repertoire model all have explosive state spaces, which increase rapidly with the number of analyzed biomes, regions, or host species, respectively. Data augmentation methods (Robinson et al. 2003; Lartillot and Poujol 2011; Landis et al. 2013; Quintero and Landis 2020; Braga et al. 2020) have proven useful for circumventing state space explosions for modeling anagenetic change. Integrating cladogenetic event inference into data augmented methods is an important future direction that will allow many features from *RFBS* and other methods to be useful for systems where the number of discrete niche-set states (either hosts repertoires, biome affinity sets, etc.) is large.

Phylogenetic signals of biome conservatism or divergence inferred solely from established affinity data (Crisp et al. 2009) cannot resolve how much those patterns are shaped by evolutionary versus ecological or geographical forces. To better understand to what extent biome affinities are shaped by these factors, *RFBS* could be modified to consider the geographical distribution of the biomes themselves (Landis et al. 2021b,a). Such an approach could would allow biome shift rates to be functions of geographic and/or climatic distances (Cardillo et al. 2017; Rinćon-Barrado et al. 2021; Quintero et al. 2023).

This addition would be important, because *RFBS* currently does not allow for species to gain established affinities without enabled affinities and defines established affinity as stable persistence within a biome, rather than any instance of individuals of a species occurring within a biome. Individuals of a species that enter a biome outside of its fundamental niche are expected to either rapidly go extinct or adapt so quickly that at a macroevolutionary scale, it is indiscernible from our double gain “lockstep” event (Edwards et al. 2017). However, being present in a biome without an enabled affinity for it necessarily entails an increased probability of local extinction until that enabled affinity is gained. Without explicitly accounting for this increased extinction probability, it is difficult for *RFBS* to restrict species from occupying a biome for millions of years without the requisite enabled affinity – an unrealistic scenario. When viewed through a historical lens, species can certainly occur in biomes to which they lack an enabled affinity, as paleobiome dynamics can cause rapid transitions in local environmental conditions and community composition. A species solely enabled to inhabit warm forests would face three fates if its local habitat was rapidly replaced with boreal forests: it would need to either adapt to the newly emergent cold conditions, move to track warmer habitats, or simply go extinct.

An important extension of *RFBS* would be to model established and enabled biome affinities in a state dependent speciation and extinction (SSE) context (Maddison et al. 2007; Goldberg et al. 2011; Caetano et al. 2018; Landis et al. 2022). Biome shifts are often hypothesized to be major drivers of diversification, where diversification rates for lineages increase as they access new biomes, and rates decrease for lineages that are restricted to biomes close to their niche optima (Brennan and Oliver 2017; Ashman et al. 2018; Quintero et al. 2023). Current analyses for biome-dependent diversification scenarios focus on established affinities, not on how enabled affinities may shape outcomes. This leaves a variety of interesting questions unanswerable using standard SSE models. For example, does a wide breadth of enabled affinities increase speciation rate and/or decrease extinction rate? Or, are species more likely to move, evolve, or go extinct when they have no enabled affinity with the biome they occupy? Such questions can be tested directly with state-dependent diversification processes that allow enabled and established biome affinities to evolve in distinct yet concerted ways.

## Conclusion

Distinguishing which biomes species *do* occupy from those biomes they *could* occupy is critical for learning how evolutionary, ecological, and geographical scenarios shape biodiversity patterns. The distinction has implications for how we expect past, present, and future species to contend with ecosystem-level changes, such as those instigated by climate change or community turnover. With *RFBS*, we introduce a phylogenetic approach to explicitly model shifts in both the established affinities with biomes that species do occupy and shifts in enabled affinities for biomes that species could occupy. By treating enabled and established affinities as states that change and are inherited in distinct ways, *RFBS* may improve how accurately we reconstruct historical biogeographic patterns concerning the movement and evolution of species among biomes. Our hope is that *RFBS* will allow biologists to onboard organismal knowledge (e.g. physiological, experimental, and horticultural data) to better understand phylogenetic biome shifts.

## Acknowledgements

Sarah Swiston, Fábio Mendes, Mariana Braga, Isaac Lichter Marck, Felipe Zapata, Adam Smith, and Kelly Zamudio provided valuable feedback that improved the design of the model and the clarity of the manuscript. We are grateful to Rebecca Sucher and Emily Warchefsky at Missouri Botanical Garden for help accessing the Botanical Garden Conservation International Plant Search Database. We also express thanks to Michael Dossman (Arnold Arboretum), Jess Goehler (Chicago Botanical Garden), and Pam Morris Olshefski (Morris Arboretum) for their assistance confirming living collection records for *Viburnum*.

*

