## Supplementary Materials for "A Phylogenetic Model of Established and Enabled Biome Shifts"

Version dated: August 29, 2024

### Supplement 1: Supplementary Model Details

The anagenetic rate matrix, $Q$, and cladogenetic probability matrix, $P$ (one matrix per starting state), for a system with three biomes and 19 biome affinity configurations are presented below.

$Q =$

| | $T$ | $W$ | $C$ | $Tw$ | $Tc$ | $Wt$ | $Wc$ | $Ct$ | $Cw$ | $TW$ | $TC$ | $WC$ | $Twc$ | $tWc$ | $twC$ | $TWc$ | $TwC$ | $tWC$ | $TWC$ |
|---|---|---|---|---|---|---|---|---|---|---|---|---|---|---|---|---|---|---|---|
| $T$ | – | 0 | 0 | $g_{0\to1}$ | $g_{0\to1}$ | 0 | 0 | 0 | 0 | $g_{0\to2}$ | $g_{0\to2}$ | 0 | 0 | 0 | 0 | 0 | 0 | 0 | 0 |
| $W$ | 0 | – | 0 | 0 | 0 | $g_{0\to1}$ | $g_{0\to1}$ | 0 | 0 | $g_{0\to2}$ | 0 | $g_{0\to2}$ | 0 | 0 | 0 | 0 | 0 | 0 | 0 |
| $C$ | 0 | 0 | – | 0 | 0 | 0 | 0 | $g_{0\to1}$ | $g_{0\to1}$ | 0 | $g_{0\to2}$ | $g_{0\to2}$ | 0 | 0 | 0 | 0 | 0 | 0 | 0 |
| $Tw$ | $l_{1\to0}$ | 0 | 0 | – | 0 | 0 | 0 | 0 | 0 | $g_{1\to2}$ | 0 | 0 | $g_{0\to1}$ | 0 | 0 | 0 | $g_{0\to2}$ | 0 | 0 |
| $Tc$ | $l_{1\to0}$ | 0 | 0 | 0 | – | 0 | 0 | 0 | 0 | 0 | $g_{1\to2}$ | 0 | $g_{0\to1}$ | 0 | 0 | $g_{0\to2}$ | 0 | 0 | 0 |
| $Wt$ | 0 | $l_{1\to0}$ | 0 | 0 | 0 | – | 0 | 0 | 0 | $g_{1\to2}$ | 0 | 0 | 0 | $g_{0\to1}$ | 0 | 0 | 0 | $g_{0\to2}$ | 0 |
| $Wc$ | 0 | $l_{1\to0}$ | 0 | 0 | 0 | 0 | – | 0 | 0 | 0 | 0 | $g_{1\to2}$ | 0 | $g_{0\to1}$ | 0 | $g_{0\to2}$ | 0 | 0 | 0 |
| $Ct$ | 0 | 0 | $l_{1\to0}$ | 0 | 0 | 0 | 0 | – | 0 | 0 | $g_{1\to2}$ | 0 | 0 | 0 | $g_{0\to1}$ | 0 | 0 | $g_{0\to2}$ | 0 |
| $Cw$ | 0 | 0 | $l_{1\to0}$ | 0 | 0 | 0 | 0 | 0 | – | 0 | 0 | $g_{1\to2}$ | 0 | 0 | $g_{0\to1}$ | 0 | $g_{0\to2}$ | 0 | 0 |
| $TW$ | 0 | 0 | 0 | $l_{2\to1}$ | 0 | $l_{2\to1}$ | 0 | 0 | 0 | – | 0 | 0 | 0 | 0 | 0 | $g_{0\to1}$ | 0 | 0 | $g_{0\to2}$ |
| $TC$ | 0 | 0 | 0 | 0 | $l_{2\to1}$ | 0 | 0 | $l_{2\to1}$ | 0 | 0 | – | 0 | 0 | 0 | 0 | 0 | $g_{0\to1}$ | 0 | $g_{0\to2}$ |
| $WC$ | 0 | 0 | 0 | 0 | 0 | 0 | $l_{2\to1}$ | 0 | $l_{2\to1}$ | 0 | 0 | – | 0 | 0 | 0 | 0 | 0 | $g_{0\to1}$ | $g_{0\to2}$ |
| $Twc$ | 0 | 0 | 0 | $l_{1\to0}$ | $l_{1\to0}$ | 0 | 0 | 0 | 0 | 0 | 0 | 0 | – | 0 | 0 | $g_{1\to2}$ | $g_{1\to2}$ | 0 | 0 |
| $tWc$ | 0 | 0 | 0 | 0 | 0 | $l_{1\to0}$ | $l_{1\to0}$ | 0 | 0 | 0 | 0 | 0 | 0 | – | 0 | $g_{1\to2}$ | 0 | $g_{1\to2}$ | 0 |
| $twC$ | 0 | 0 | 0 | 0 | 0 | 0 | 0 | $l_{1\to0}$ | $l_{1\to0}$ | 0 | 0 | 0 | 0 | 0 | – | 0 | $g_{1\to2}$ | $g_{1\to2}$ | 0 |
| $TWc$ | 0 | 0 | 0 | 0 | 0 | 0 | 0 | 0 | 0 | $l_{1\to0}$ | 0 | 0 | $l_{2\to1}$ | $l_{2\to1}$ | 0 | – | 0 | 0 | $g_{1\to2}$ |
| $TwC$ | 0 | 0 | 0 | 0 | 0 | 0 | 0 | 0 | 0 | 0 | $l_{1\to0}$ | 0 | $l_{2\to1}$ | 0 | $l_{2\to1}$ | 0 | – | 0 | $g_{1\to2}$ |
| $tWC$ | 0 | 0 | 0 | 0 | 0 | 0 | 0 | 0 | 0 | 0 | 0 | $l_{1\to0}$ | 0 | $l_{2\to1}$ | $l_{2\to1}$ | 0 | 0 | – | $g_{1\to2}$ |
| $TWC$ | 0 | 0 | 0 | 0 | 0 | 0 | 0 | 0 | 0 | 0 | 0 | 0 | 0 | 0 | 0 | $l_{2\to1}$ | $l_{2\to1}$ | $l_{2\to1}$ | – |

$P_T =$

| | $T$ | $W$ | $C$ | $Tw$ | $Tc$ | $Wt$ | $Wc$ | $Ct$ | $Cw$ | $TW$ | $TC$ | $WC$ | $Twc$ | $tWc$ | $twC$ | $TWc$ | $TwC$ | $tWC$ | $TWC$ |
|---|---|---|---|---|---|---|---|---|---|---|---|---|---|---|---|---|---|---|---|
| $T$ | 1 | 0 | 0 | 0 | 0 | 0 | 0 | 0 | 0 | 0 | 0 | 0 | 0 | 0 | 0 | 0 | 0 | 0 | 0 |
| $W$ | 0 | 0 | 0 | 0 | 0 | 0 | 0 | 0 | 0 | 0 | 0 | 0 | 0 | 0 | 0 | 0 | 0 | 0 | 0 |
| $C$ | 0 | 0 | 0 | 0 | 0 | 0 | 0 | 0 | 0 | 0 | 0 | 0 | 0 | 0 | 0 | 0 | 0 | 0 | 0 |
| $Tw$ | 0 | 0 | 0 | 0 | 0 | 0 | 0 | 0 | 0 | 0 | 0 | 0 | 0 | 0 | 0 | 0 | 0 | 0 | 0 |
| $Tc$ | 0 | 0 | 0 | 0 | 0 | 0 | 0 | 0 | 0 | 0 | 0 | 0 | 0 | 0 | 0 | 0 | 0 | 0 | 0 |
| $Wt$ | 0 | 0 | 0 | 0 | 0 | 0 | 0 | 0 | 0 | 0 | 0 | 0 | 0 | 0 | 0 | 0 | 0 | 0 | 0 |
| $Wc$ | 0 | 0 | 0 | 0 | 0 | 0 | 0 | 0 | 0 | 0 | 0 | 0 | 0 | 0 | 0 | 0 | 0 | 0 | 0 |
| $Ct$ | 0 | 0 | 0 | 0 | 0 | 0 | 0 | 0 | 0 | 0 | 0 | 0 | 0 | 0 | 0 | 0 | 0 | 0 | 0 |
| $Cw$ | 0 | 0 | 0 | 0 | 0 | 0 | 0 | 0 | 0 | 0 | 0 | 0 | 0 | 0 | 0 | 0 | 0 | 0 | 0 |
| $TW$ | 0 | 0 | 0 | 0 | 0 | 0 | 0 | 0 | 0 | 0 | 0 | 0 | 0 | 0 | 0 | 0 | 0 | 0 | 0 |
| $TC$ | 0 | 0 | 0 | 0 | 0 | 0 | 0 | 0 | 0 | 0 | 0 | 0 | 0 | 0 | 0 | 0 | 0 | 0 | 0 |
| $WC$ | 0 | 0 | 0 | 0 | 0 | 0 | 0 | 0 | 0 | 0 | 0 | 0 | 0 | 0 | 0 | 0 | 0 | 0 | 0 |
| $Twc$ | 0 | 0 | 0 | 0 | 0 | 0 | 0 | 0 | 0 | 0 | 0 | 0 | 0 | 0 | 0 | 0 | 0 | 0 | 0 |
| $tWc$ | 0 | 0 | 0 | 0 | 0 | 0 | 0 | 0 | 0 | 0 | 0 | 0 | 0 | 0 | 0 | 0 | 0 | 0 | 0 |
| $twC$ | 0 | 0 | 0 | 0 | 0 | 0 | 0 | 0 | 0 | 0 | 0 | 0 | 0 | 0 | 0 | 0 | 0 | 0 | 0 |
| $TWc$ | 0 | 0 | 0 | 0 | 0 | 0 | 0 | 0 | 0 | 0 | 0 | 0 | 0 | 0 | 0 | 0 | 0 | 0 | 0 |
| $TwC$ | 0 | 0 | 0 | 0 | 0 | 0 | 0 | 0 | 0 | 0 | 0 | 0 | 0 | 0 | 0 | 0 | 0 | 0 | 0 |
| $tWC$ | 0 | 0 | 0 | 0 | 0 | 0 | 0 | 0 | 0 | 0 | 0 | 0 | 0 | 0 | 0 | 0 | 0 | 0 | 0 |
| $TWC$ | 0 | 0 | 0 | 0 | 0 | 0 | 0 | 0 | 0 | 0 | 0 | 0 | 0 | 0 | 0 | 0 | 0 | 0 | 0 |

$$P_W = \left(\begin{array}{c|ccccccccccccccccccc}
 & T & W & C & Tw & Tc & Wt & Wc & Ct & Cw & TW & TC & WC & Twc & tWc & twC & TWc & TwC & tWC & TWC \\
\hline
T & 0 & 0 & 0 & 0 & 0 & 0 & 0 & 0 & 0 & 0 & 0 & 0 & 0 & 0 & 0 & 0 & 0 & 0 & 0 \\
W & 0 & 1 & 0 & 0 & 0 & 0 & 0 & 0 & 0 & 0 & 0 & 0 & 0 & 0 & 0 & 0 & 0 & 0 & 0 \\
C & 0 & 0 & 0 & 0 & 0 & 0 & 0 & 0 & 0 & 0 & 0 & 0 & 0 & 0 & 0 & 0 & 0 & 0 & 0 \\
Tw & 0 & 0 & 0 & 0 & 0 & 0 & 0 & 0 & 0 & 0 & 0 & 0 & 0 & 0 & 0 & 0 & 0 & 0 & 0 \\
Tc & 0 & 0 & 0 & 0 & 0 & 0 & 0 & 0 & 0 & 0 & 0 & 0 & 0 & 0 & 0 & 0 & 0 & 0 & 0 \\
Wt & 0 & 0 & 0 & 0 & 0 & 0 & 0 & 0 & 0 & 0 & 0 & 0 & 0 & 0 & 0 & 0 & 0 & 0 & 0 \\
Wc & 0 & 0 & 0 & 0 & 0 & 0 & 0 & 0 & 0 & 0 & 0 & 0 & 0 & 0 & 0 & 0 & 0 & 0 & 0 \\
Ct & 0 & 0 & 0 & 0 & 0 & 0 & 0 & 0 & 0 & 0 & 0 & 0 & 0 & 0 & 0 & 0 & 0 & 0 & 0 \\
Cw & 0 & 0 & 0 & 0 & 0 & 0 & 0 & 0 & 0 & 0 & 0 & 0 & 0 & 0 & 0 & 0 & 0 & 0 & 0 \\
TW & 0 & 0 & 0 & 0 & 0 & 0 & 0 & 0 & 0 & 0 & 0 & 0 & 0 & 0 & 0 & 0 & 0 & 0 & 0 \\
TC & 0 & 0 & 0 & 0 & 0 & 0 & 0 & 0 & 0 & 0 & 0 & 0 & 0 & 0 & 0 & 0 & 0 & 0 & 0 \\
WC & 0 & 0 & 0 & 0 & 0 & 0 & 0 & 0 & 0 & 0 & 0 & 0 & 0 & 0 & 0 & 0 & 0 & 0 & 0 \\
Twc & 0 & 0 & 0 & 0 & 0 & 0 & 0 & 0 & 0 & 0 & 0 & 0 & 0 & 0 & 0 & 0 & 0 & 0 & 0 \\
tWc & 0 & 0 & 0 & 0 & 0 & 0 & 0 & 0 & 0 & 0 & 0 & 0 & 0 & 0 & 0 & 0 & 0 & 0 & 0 \\
twC & 0 & 0 & 0 & 0 & 0 & 0 & 0 & 0 & 0 & 0 & 0 & 0 & 0 & 0 & 0 & 0 & 0 & 0 & 0 \\
TWc & 0 & 0 & 0 & 0 & 0 & 0 & 0 & 0 & 0 & 0 & 0 & 0 & 0 & 0 & 0 & 0 & 0 & 0 & 0 \\
TwC & 0 & 0 & 0 & 0 & 0 & 0 & 0 & 0 & 0 & 0 & 0 & 0 & 0 & 0 & 0 & 0 & 0 & 0 & 0 \\
tWC & 0 & 0 & 0 & 0 & 0 & 0 & 0 & 0 & 0 & 0 & 0 & 0 & 0 & 0 & 0 & 0 & 0 & 0 & 0 \\
TWC & 0 & 0 & 0 & 0 & 0 & 0 & 0 & 0 & 0 & 0 & 0 & 0 & 0 & 0 & 0 & 0 & 0 & 0 & 0
\end{array}\right)$$

$$P_C = \left(\begin{array}{c|ccccccccccccccccccc}
 & T & W & C & Tw & Tc & Wt & Wc & Ct & Cw & TW & TC & WC & Twc & tWc & twC & TWc & TwC & tWC & TWC \\
\hline
T & 0 & 0 & 0 & 0 & 0 & 0 & 0 & 0 & 0 & 0 & 0 & 0 & 0 & 0 & 0 & 0 & 0 & 0 & 0 \\
W & 0 & 0 & 0 & 0 & 0 & 0 & 0 & 0 & 0 & 0 & 0 & 0 & 0 & 0 & 0 & 0 & 0 & 0 & 0 \\
C & 0 & 0 & 1 & 0 & 0 & 0 & 0 & 0 & 0 & 0 & 0 & 0 & 0 & 0 & 0 & 0 & 0 & 0 & 0 \\
Tw & 0 & 0 & 0 & 0 & 0 & 0 & 0 & 0 & 0 & 0 & 0 & 0 & 0 & 0 & 0 & 0 & 0 & 0 & 0 \\
Tc & 0 & 0 & 0 & 0 & 0 & 0 & 0 & 0 & 0 & 0 & 0 & 0 & 0 & 0 & 0 & 0 & 0 & 0 & 0 \\
Wt & 0 & 0 & 0 & 0 & 0 & 0 & 0 & 0 & 0 & 0 & 0 & 0 & 0 & 0 & 0 & 0 & 0 & 0 & 0 \\
Wc & 0 & 0 & 0 & 0 & 0 & 0 & 0 & 0 & 0 & 0 & 0 & 0 & 0 & 0 & 0 & 0 & 0 & 0 & 0 \\
Ct & 0 & 0 & 0 & 0 & 0 & 0 & 0 & 0 & 0 & 0 & 0 & 0 & 0 & 0 & 0 & 0 & 0 & 0 & 0 \\
Cw & 0 & 0 & 0 & 0 & 0 & 0 & 0 & 0 & 0 & 0 & 0 & 0 & 0 & 0 & 0 & 0 & 0 & 0 & 0 \\
TW & 0 & 0 & 0 & 0 & 0 & 0 & 0 & 0 & 0 & 0 & 0 & 0 & 0 & 0 & 0 & 0 & 0 & 0 & 0 \\
TC & 0 & 0 & 0 & 0 & 0 & 0 & 0 & 0 & 0 & 0 & 0 & 0 & 0 & 0 & 0 & 0 & 0 & 0 & 0 \\
WC & 0 & 0 & 0 & 0 & 0 & 0 & 0 & 0 & 0 & 0 & 0 & 0 & 0 & 0 & 0 & 0 & 0 & 0 & 0 \\
Twc & 0 & 0 & 0 & 0 & 0 & 0 & 0 & 0 & 0 & 0 & 0 & 0 & 0 & 0 & 0 & 0 & 0 & 0 & 0 \\
tWc & 0 & 0 & 0 & 0 & 0 & 0 & 0 & 0 & 0 & 0 & 0 & 0 & 0 & 0 & 0 & 0 & 0 & 0 & 0 \\
twC & 0 & 0 & 0 & 0 & 0 & 0 & 0 & 0 & 0 & 0 & 0 & 0 & 0 & 0 & 0 & 0 & 0 & 0 & 0 \\
TWc & 0 & 0 & 0 & 0 & 0 & 0 & 0 & 0 & 0 & 0 & 0 & 0 & 0 & 0 & 0 & 0 & 0 & 0 & 0 \\
TwC & 0 & 0 & 0 & 0 & 0 & 0 & 0 & 0 & 0 & 0 & 0 & 0 & 0 & 0 & 0 & 0 & 0 & 0 & 0 \\
tWC & 0 & 0 & 0 & 0 & 0 & 0 & 0 & 0 & 0 & 0 & 0 & 0 & 0 & 0 & 0 & 0 & 0 & 0 & 0 \\
TWC & 0 & 0 & 0 & 0 & 0 & 0 & 0 & 0 & 0 & 0 & 0 & 0 & 0 & 0 & 0 & 0 & 0 & 0 & 0
\end{array}\right)$$

$P_{Tw} =$

| | $T$ | $W$ | $C$ | $Tw$ | $Tc$ | $Wt$ | $Wc$ | $Ct$ | $Cw$ | $TW$ | $TC$ | $WC$ | $Twc$ | $tWc$ | $twC$ | $TWc$ | $TwC$ | $tWC$ | $TWC$ |
|---|---|---|---|---|---|---|---|---|---|---|---|---|---|---|---|---|---|---|---|
| $T$ | 0 | 0 | 0 | 0 | 0 | 0 | 0 | 0 | 0 | 0 | 0 | 0 | 0 | 0 | 0 | 0 | 0 | 0 | 0 |
| $W$ | 0 | 0 | 0 | 0 | 0 | 0 | 0 | 0 | 0 | 0 | 0 | 0 | 0 | 0 | 0 | 0 | 0 | 0 | 0 |
| $C$ | 0 | 0 | 0 | 0 | 0 | 0 | 0 | 0 | 0 | 0 | 0 | 0 | 0 | 0 | 0 | 0 | 0 | 0 | 0 |
| $Tw$ | 0 | 0 | 0 | 1 | 0 | 0 | 0 | 0 | 0 | 0 | 0 | 0 | 0 | 0 | 0 | 0 | 0 | 0 | 0 |
| $Tc$ | 0 | 0 | 0 | 0 | 0 | 0 | 0 | 0 | 0 | 0 | 0 | 0 | 0 | 0 | 0 | 0 | 0 | 0 | 0 |
| $Wt$ | 0 | 0 | 0 | 0 | 0 | 0 | 0 | 0 | 0 | 0 | 0 | 0 | 0 | 0 | 0 | 0 | 0 | 0 | 0 |
| $Wc$ | 0 | 0 | 0 | 0 | 0 | 0 | 0 | 0 | 0 | 0 | 0 | 0 | 0 | 0 | 0 | 0 | 0 | 0 | 0 |
| $Ct$ | 0 | 0 | 0 | 0 | 0 | 0 | 0 | 0 | 0 | 0 | 0 | 0 | 0 | 0 | 0 | 0 | 0 | 0 | 0 |
| $Cw$ | 0 | 0 | 0 | 0 | 0 | 0 | 0 | 0 | 0 | 0 | 0 | 0 | 0 | 0 | 0 | 0 | 0 | 0 | 0 |
| $TW$ | 0 | 0 | 0 | 0 | 0 | 0 | 0 | 0 | 0 | 0 | 0 | 0 | 0 | 0 | 0 | 0 | 0 | 0 | 0 |
| $TC$ | 0 | 0 | 0 | 0 | 0 | 0 | 0 | 0 | 0 | 0 | 0 | 0 | 0 | 0 | 0 | 0 | 0 | 0 | 0 |
| $WC$ | 0 | 0 | 0 | 0 | 0 | 0 | 0 | 0 | 0 | 0 | 0 | 0 | 0 | 0 | 0 | 0 | 0 | 0 | 0 |
| $Twc$ | 0 | 0 | 0 | 0 | 0 | 0 | 0 | 0 | 0 | 0 | 0 | 0 | 0 | 0 | 0 | 0 | 0 | 0 | 0 |
| $tWc$ | 0 | 0 | 0 | 0 | 0 | 0 | 0 | 0 | 0 | 0 | 0 | 0 | 0 | 0 | 0 | 0 | 0 | 0 | 0 |
| $twC$ | 0 | 0 | 0 | 0 | 0 | 0 | 0 | 0 | 0 | 0 | 0 | 0 | 0 | 0 | 0 | 0 | 0 | 0 | 0 |
| $TWc$ | 0 | 0 | 0 | 0 | 0 | 0 | 0 | 0 | 0 | 0 | 0 | 0 | 0 | 0 | 0 | 0 | 0 | 0 | 0 |
| $TwC$ | 0 | 0 | 0 | 0 | 0 | 0 | 0 | 0 | 0 | 0 | 0 | 0 | 0 | 0 | 0 | 0 | 0 | 0 | 0 |
| $tWC$ | 0 | 0 | 0 | 0 | 0 | 0 | 0 | 0 | 0 | 0 | 0 | 0 | 0 | 0 | 0 | 0 | 0 | 0 | 0 |
| $TWC$ | 0 | 0 | 0 | 0 | 0 | 0 | 0 | 0 | 0 | 0 | 0 | 0 | 0 | 0 | 0 | 0 | 0 | 0 | 0 |

$P_{Tc} =$

| | $T$ | $W$ | $C$ | $Tw$ | $Tc$ | $Wt$ | $Wc$ | $Ct$ | $Cw$ | $TW$ | $TC$ | $WC$ | $Twc$ | $tWc$ | $twC$ | $TWc$ | $TwC$ | $tWC$ | $TWC$ |
|---|---|---|---|---|---|---|---|---|---|---|---|---|---|---|---|---|---|---|---|
| $T$ | 0 | 0 | 0 | 0 | 0 | 0 | 0 | 0 | 0 | 0 | 0 | 0 | 0 | 0 | 0 | 0 | 0 | 0 | 0 |
| $W$ | 0 | 0 | 0 | 0 | 0 | 0 | 0 | 0 | 0 | 0 | 0 | 0 | 0 | 0 | 0 | 0 | 0 | 0 | 0 |
| $C$ | 0 | 0 | 0 | 0 | 0 | 0 | 0 | 0 | 0 | 0 | 0 | 0 | 0 | 0 | 0 | 0 | 0 | 0 | 0 |
| $Tw$ | 0 | 0 | 0 | 0 | 0 | 0 | 0 | 0 | 0 | 0 | 0 | 0 | 0 | 0 | 0 | 0 | 0 | 0 | 0 |
| $Tc$ | 0 | 0 | 0 | 0 | 1 | 0 | 0 | 0 | 0 | 0 | 0 | 0 | 0 | 0 | 0 | 0 | 0 | 0 | 0 |
| $Wt$ | 0 | 0 | 0 | 0 | 0 | 0 | 0 | 0 | 0 | 0 | 0 | 0 | 0 | 0 | 0 | 0 | 0 | 0 | 0 |
| $Wc$ | 0 | 0 | 0 | 0 | 0 | 0 | 0 | 0 | 0 | 0 | 0 | 0 | 0 | 0 | 0 | 0 | 0 | 0 | 0 |
| $Ct$ | 0 | 0 | 0 | 0 | 0 | 0 | 0 | 0 | 0 | 0 | 0 | 0 | 0 | 0 | 0 | 0 | 0 | 0 | 0 |
| $Cw$ | 0 | 0 | 0 | 0 | 0 | 0 | 0 | 0 | 0 | 0 | 0 | 0 | 0 | 0 | 0 | 0 | 0 | 0 | 0 |
| $TW$ | 0 | 0 | 0 | 0 | 0 | 0 | 0 | 0 | 0 | 0 | 0 | 0 | 0 | 0 | 0 | 0 | 0 | 0 | 0 |
| $TC$ | 0 | 0 | 0 | 0 | 0 | 0 | 0 | 0 | 0 | 0 | 0 | 0 | 0 | 0 | 0 | 0 | 0 | 0 | 0 |
| $WC$ | 0 | 0 | 0 | 0 | 0 | 0 | 0 | 0 | 0 | 0 | 0 | 0 | 0 | 0 | 0 | 0 | 0 | 0 | 0 |
| $Twc$ | 0 | 0 | 0 | 0 | 0 | 0 | 0 | 0 | 0 | 0 | 0 | 0 | 0 | 0 | 0 | 0 | 0 | 0 | 0 |
| $tWc$ | 0 | 0 | 0 | 0 | 0 | 0 | 0 | 0 | 0 | 0 | 0 | 0 | 0 | 0 | 0 | 0 | 0 | 0 | 0 |
| $twC$ | 0 | 0 | 0 | 0 | 0 | 0 | 0 | 0 | 0 | 0 | 0 | 0 | 0 | 0 | 0 | 0 | 0 | 0 | 0 |
| $TWc$ | 0 | 0 | 0 | 0 | 0 | 0 | 0 | 0 | 0 | 0 | 0 | 0 | 0 | 0 | 0 | 0 | 0 | 0 | 0 |
| $TwC$ | 0 | 0 | 0 | 0 | 0 | 0 | 0 | 0 | 0 | 0 | 0 | 0 | 0 | 0 | 0 | 0 | 0 | 0 | 0 |
| $tWC$ | 0 | 0 | 0 | 0 | 0 | 0 | 0 | 0 | 0 | 0 | 0 | 0 | 0 | 0 | 0 | 0 | 0 | 0 | 0 |
| $TWC$ | 0 | 0 | 0 | 0 | 0 | 0 | 0 | 0 | 0 | 0 | 0 | 0 | 0 | 0 | 0 | 0 | 0 | 0 | 0 |

$$P_{Wt} = \left(\begin{array}{c|ccccccccccccccccccc}
 & T & W & C & Tw & Tc & Wt & Wc & Ct & Cw & TW & TC & WC & Twc & tWc & twC & TWc & TwC & tWC & TWC \\
\hline
T & 0 & 0 & 0 & 0 & 0 & 0 & 0 & 0 & 0 & 0 & 0 & 0 & 0 & 0 & 0 & 0 & 0 & 0 & 0 \\
W & 0 & 0 & 0 & 0 & 0 & 0 & 0 & 0 & 0 & 0 & 0 & 0 & 0 & 0 & 0 & 0 & 0 & 0 & 0 \\
C & 0 & 0 & 0 & 0 & 0 & 0 & 0 & 0 & 0 & 0 & 0 & 0 & 0 & 0 & 0 & 0 & 0 & 0 & 0 \\
Tw & 0 & 0 & 0 & 0 & 0 & 0 & 0 & 0 & 0 & 0 & 0 & 0 & 0 & 0 & 0 & 0 & 0 & 0 & 0 \\
Tc & 0 & 0 & 0 & 0 & 0 & 0 & 0 & 0 & 0 & 0 & 0 & 0 & 0 & 0 & 0 & 0 & 0 & 0 & 0 \\
Wt & 0 & 0 & 0 & 0 & 0 & 1 & 0 & 0 & 0 & 0 & 0 & 0 & 0 & 0 & 0 & 0 & 0 & 0 & 0 \\
Wc & 0 & 0 & 0 & 0 & 0 & 0 & 0 & 0 & 0 & 0 & 0 & 0 & 0 & 0 & 0 & 0 & 0 & 0 & 0 \\
Ct & 0 & 0 & 0 & 0 & 0 & 0 & 0 & 0 & 0 & 0 & 0 & 0 & 0 & 0 & 0 & 0 & 0 & 0 & 0 \\
Cw & 0 & 0 & 0 & 0 & 0 & 0 & 0 & 0 & 0 & 0 & 0 & 0 & 0 & 0 & 0 & 0 & 0 & 0 & 0 \\
TW & 0 & 0 & 0 & 0 & 0 & 0 & 0 & 0 & 0 & 0 & 0 & 0 & 0 & 0 & 0 & 0 & 0 & 0 & 0 \\
TC & 0 & 0 & 0 & 0 & 0 & 0 & 0 & 0 & 0 & 0 & 0 & 0 & 0 & 0 & 0 & 0 & 0 & 0 & 0 \\
WC & 0 & 0 & 0 & 0 & 0 & 0 & 0 & 0 & 0 & 0 & 0 & 0 & 0 & 0 & 0 & 0 & 0 & 0 & 0 \\
Twc & 0 & 0 & 0 & 0 & 0 & 0 & 0 & 0 & 0 & 0 & 0 & 0 & 0 & 0 & 0 & 0 & 0 & 0 & 0 \\
tWc & 0 & 0 & 0 & 0 & 0 & 0 & 0 & 0 & 0 & 0 & 0 & 0 & 0 & 0 & 0 & 0 & 0 & 0 & 0 \\
twC & 0 & 0 & 0 & 0 & 0 & 0 & 0 & 0 & 0 & 0 & 0 & 0 & 0 & 0 & 0 & 0 & 0 & 0 & 0 \\
TWc & 0 & 0 & 0 & 0 & 0 & 0 & 0 & 0 & 0 & 0 & 0 & 0 & 0 & 0 & 0 & 0 & 0 & 0 & 0 \\
TwC & 0 & 0 & 0 & 0 & 0 & 0 & 0 & 0 & 0 & 0 & 0 & 0 & 0 & 0 & 0 & 0 & 0 & 0 & 0 \\
tWC & 0 & 0 & 0 & 0 & 0 & 0 & 0 & 0 & 0 & 0 & 0 & 0 & 0 & 0 & 0 & 0 & 0 & 0 & 0 \\
TWC & 0 & 0 & 0 & 0 & 0 & 0 & 0 & 0 & 0 & 0 & 0 & 0 & 0 & 0 & 0 & 0 & 0 & 0 & 0
\end{array}\right)$$

$$P_{Wc} = \left(\begin{array}{c|ccccccccccccccccccc}
 & T & W & C & Tw & Tc & Wt & Wc & Ct & Cw & TW & TC & WC & Twc & tWc & twC & TWc & TwC & tWC & TWC \\
\hline
T & 0 & 0 & 0 & 0 & 0 & 0 & 0 & 0 & 0 & 0 & 0 & 0 & 0 & 0 & 0 & 0 & 0 & 0 & 0 \\
W & 0 & 0 & 0 & 0 & 0 & 0 & 0 & 0 & 0 & 0 & 0 & 0 & 0 & 0 & 0 & 0 & 0 & 0 & 0 \\
C & 0 & 0 & 0 & 0 & 0 & 0 & 0 & 0 & 0 & 0 & 0 & 0 & 0 & 0 & 0 & 0 & 0 & 0 & 0 \\
Tw & 0 & 0 & 0 & 0 & 0 & 0 & 0 & 0 & 0 & 0 & 0 & 0 & 0 & 0 & 0 & 0 & 0 & 0 & 0 \\
Tc & 0 & 0 & 0 & 0 & 0 & 0 & 0 & 0 & 0 & 0 & 0 & 0 & 0 & 0 & 0 & 0 & 0 & 0 & 0 \\
Wt & 0 & 0 & 0 & 0 & 0 & 0 & 0 & 0 & 0 & 0 & 0 & 0 & 0 & 0 & 0 & 0 & 0 & 0 & 0 \\
Wc & 0 & 0 & 0 & 0 & 0 & 0 & 1 & 0 & 0 & 0 & 0 & 0 & 0 & 0 & 0 & 0 & 0 & 0 & 0 \\
Ct & 0 & 0 & 0 & 0 & 0 & 0 & 0 & 0 & 0 & 0 & 0 & 0 & 0 & 0 & 0 & 0 & 0 & 0 & 0 \\
Cw & 0 & 0 & 0 & 0 & 0 & 0 & 0 & 0 & 0 & 0 & 0 & 0 & 0 & 0 & 0 & 0 & 0 & 0 & 0 \\
TW & 0 & 0 & 0 & 0 & 0 & 0 & 0 & 0 & 0 & 0 & 0 & 0 & 0 & 0 & 0 & 0 & 0 & 0 & 0 \\
TC & 0 & 0 & 0 & 0 & 0 & 0 & 0 & 0 & 0 & 0 & 0 & 0 & 0 & 0 & 0 & 0 & 0 & 0 & 0 \\
WC & 0 & 0 & 0 & 0 & 0 & 0 & 0 & 0 & 0 & 0 & 0 & 0 & 0 & 0 & 0 & 0 & 0 & 0 & 0 \\
Twc & 0 & 0 & 0 & 0 & 0 & 0 & 0 & 0 & 0 & 0 & 0 & 0 & 0 & 0 & 0 & 0 & 0 & 0 & 0 \\
tWc & 0 & 0 & 0 & 0 & 0 & 0 & 0 & 0 & 0 & 0 & 0 & 0 & 0 & 0 & 0 & 0 & 0 & 0 & 0 \\
twC & 0 & 0 & 0 & 0 & 0 & 0 & 0 & 0 & 0 & 0 & 0 & 0 & 0 & 0 & 0 & 0 & 0 & 0 & 0 \\
TWc & 0 & 0 & 0 & 0 & 0 & 0 & 0 & 0 & 0 & 0 & 0 & 0 & 0 & 0 & 0 & 0 & 0 & 0 & 0 \\
TwC & 0 & 0 & 0 & 0 & 0 & 0 & 0 & 0 & 0 & 0 & 0 & 0 & 0 & 0 & 0 & 0 & 0 & 0 & 0 \\
tWC & 0 & 0 & 0 & 0 & 0 & 0 & 0 & 0 & 0 & 0 & 0 & 0 & 0 & 0 & 0 & 0 & 0 & 0 & 0 \\
TWC & 0 & 0 & 0 & 0 & 0 & 0 & 0 & 0 & 0 & 0 & 0 & 0 & 0 & 0 & 0 & 0 & 0 & 0 & 0
\end{array}\right)$$

$P_{Ct} =$

| | $T$ | $W$ | $C$ | $Tw$ | $Tc$ | $Wt$ | $Wc$ | $Ct$ | $Cw$ | $TW$ | $TC$ | $WC$ | $Twc$ | $tWc$ | $twC$ | $TWc$ | $TwC$ | $tWC$ | $TWC$ |
|---|---|---|---|---|---|---|---|---|---|---|---|---|---|---|---|---|---|---|---|
| $T$ | 0 | 0 | 0 | 0 | 0 | 0 | 0 | 0 | 0 | 0 | 0 | 0 | 0 | 0 | 0 | 0 | 0 | 0 | 0 |
| $W$ | 0 | 0 | 0 | 0 | 0 | 0 | 0 | 0 | 0 | 0 | 0 | 0 | 0 | 0 | 0 | 0 | 0 | 0 | 0 |
| $C$ | 0 | 0 | 0 | 0 | 0 | 0 | 0 | 0 | 0 | 0 | 0 | 0 | 0 | 0 | 0 | 0 | 0 | 0 | 0 |
| $Tw$ | 0 | 0 | 0 | 0 | 0 | 0 | 0 | 0 | 0 | 0 | 0 | 0 | 0 | 0 | 0 | 0 | 0 | 0 | 0 |
| $Tc$ | 0 | 0 | 0 | 0 | 0 | 0 | 0 | 0 | 0 | 0 | 0 | 0 | 0 | 0 | 0 | 0 | 0 | 0 | 0 |
| $Wt$ | 0 | 0 | 0 | 0 | 0 | 0 | 0 | 0 | 0 | 0 | 0 | 0 | 0 | 0 | 0 | 0 | 0 | 0 | 0 |
| $Wc$ | 0 | 0 | 0 | 0 | 0 | 0 | 0 | 0 | 0 | 0 | 0 | 0 | 0 | 0 | 0 | 0 | 0 | 0 | 0 |
| $Ct$ | 0 | 0 | 0 | 0 | 0 | 0 | 0 | 1 | 0 | 0 | 0 | 0 | 0 | 0 | 0 | 0 | 0 | 0 | 0 |
| $Cw$ | 0 | 0 | 0 | 0 | 0 | 0 | 0 | 0 | 0 | 0 | 0 | 0 | 0 | 0 | 0 | 0 | 0 | 0 | 0 |
| $TW$ | 0 | 0 | 0 | 0 | 0 | 0 | 0 | 0 | 0 | 0 | 0 | 0 | 0 | 0 | 0 | 0 | 0 | 0 | 0 |
| $TC$ | 0 | 0 | 0 | 0 | 0 | 0 | 0 | 0 | 0 | 0 | 0 | 0 | 0 | 0 | 0 | 0 | 0 | 0 | 0 |
| $WC$ | 0 | 0 | 0 | 0 | 0 | 0 | 0 | 0 | 0 | 0 | 0 | 0 | 0 | 0 | 0 | 0 | 0 | 0 | 0 |
| $Twc$ | 0 | 0 | 0 | 0 | 0 | 0 | 0 | 0 | 0 | 0 | 0 | 0 | 0 | 0 | 0 | 0 | 0 | 0 | 0 |
| $tWc$ | 0 | 0 | 0 | 0 | 0 | 0 | 0 | 0 | 0 | 0 | 0 | 0 | 0 | 0 | 0 | 0 | 0 | 0 | 0 |
| $twC$ | 0 | 0 | 0 | 0 | 0 | 0 | 0 | 0 | 0 | 0 | 0 | 0 | 0 | 0 | 0 | 0 | 0 | 0 | 0 |
| $TWc$ | 0 | 0 | 0 | 0 | 0 | 0 | 0 | 0 | 0 | 0 | 0 | 0 | 0 | 0 | 0 | 0 | 0 | 0 | 0 |
| $TwC$ | 0 | 0 | 0 | 0 | 0 | 0 | 0 | 0 | 0 | 0 | 0 | 0 | 0 | 0 | 0 | 0 | 0 | 0 | 0 |
| $tWC$ | 0 | 0 | 0 | 0 | 0 | 0 | 0 | 0 | 0 | 0 | 0 | 0 | 0 | 0 | 0 | 0 | 0 | 0 | 0 |
| $TWC$ | 0 | 0 | 0 | 0 | 0 | 0 | 0 | 0 | 0 | 0 | 0 | 0 | 0 | 0 | 0 | 0 | 0 | 0 | 0 |

$P_{Cw} =$

| | $T$ | $W$ | $C$ | $Tw$ | $Tc$ | $Wt$ | $Wc$ | $Ct$ | $Cw$ | $TW$ | $TC$ | $WC$ | $Twc$ | $tWc$ | $twC$ | $TWc$ | $TwC$ | $tWC$ | $TWC$ |
|---|---|---|---|---|---|---|---|---|---|---|---|---|---|---|---|---|---|---|---|
| $T$ | 0 | 0 | 0 | 0 | 0 | 0 | 0 | 0 | 0 | 0 | 0 | 0 | 0 | 0 | 0 | 0 | 0 | 0 | 0 |
| $W$ | 0 | 0 | 0 | 0 | 0 | 0 | 0 | 0 | 0 | 0 | 0 | 0 | 0 | 0 | 0 | 0 | 0 | 0 | 0 |
| $C$ | 0 | 0 | 0 | 0 | 0 | 0 | 0 | 0 | 0 | 0 | 0 | 0 | 0 | 0 | 0 | 0 | 0 | 0 | 0 |
| $Tw$ | 0 | 0 | 0 | 0 | 0 | 0 | 0 | 0 | 0 | 0 | 0 | 0 | 0 | 0 | 0 | 0 | 0 | 0 | 0 |
| $Tc$ | 0 | 0 | 0 | 0 | 0 | 0 | 0 | 0 | 0 | 0 | 0 | 0 | 0 | 0 | 0 | 0 | 0 | 0 | 0 |
| $Wt$ | 0 | 0 | 0 | 0 | 0 | 0 | 0 | 0 | 0 | 0 | 0 | 0 | 0 | 0 | 0 | 0 | 0 | 0 | 0 |
| $Wc$ | 0 | 0 | 0 | 0 | 0 | 0 | 0 | 0 | 0 | 0 | 0 | 0 | 0 | 0 | 0 | 0 | 0 | 0 | 0 |
| $Ct$ | 0 | 0 | 0 | 0 | 0 | 0 | 0 | 0 | 0 | 0 | 0 | 0 | 0 | 0 | 0 | 0 | 0 | 0 | 0 |
| $Cw$ | 0 | 0 | 0 | 0 | 0 | 0 | 0 | 0 | 1 | 0 | 0 | 0 | 0 | 0 | 0 | 0 | 0 | 0 | 0 |
| $TW$ | 0 | 0 | 0 | 0 | 0 | 0 | 0 | 0 | 0 | 0 | 0 | 0 | 0 | 0 | 0 | 0 | 0 | 0 | 0 |
| $TC$ | 0 | 0 | 0 | 0 | 0 | 0 | 0 | 0 | 0 | 0 | 0 | 0 | 0 | 0 | 0 | 0 | 0 | 0 | 0 |
| $WC$ | 0 | 0 | 0 | 0 | 0 | 0 | 0 | 0 | 0 | 0 | 0 | 0 | 0 | 0 | 0 | 0 | 0 | 0 | 0 |
| $Twc$ | 0 | 0 | 0 | 0 | 0 | 0 | 0 | 0 | 0 | 0 | 0 | 0 | 0 | 0 | 0 | 0 | 0 | 0 | 0 |
| $tWc$ | 0 | 0 | 0 | 0 | 0 | 0 | 0 | 0 | 0 | 0 | 0 | 0 | 0 | 0 | 0 | 0 | 0 | 0 | 0 |
| $twC$ | 0 | 0 | 0 | 0 | 0 | 0 | 0 | 0 | 0 | 0 | 0 | 0 | 0 | 0 | 0 | 0 | 0 | 0 | 0 |
| $TWc$ | 0 | 0 | 0 | 0 | 0 | 0 | 0 | 0 | 0 | 0 | 0 | 0 | 0 | 0 | 0 | 0 | 0 | 0 | 0 |
| $TwC$ | 0 | 0 | 0 | 0 | 0 | 0 | 0 | 0 | 0 | 0 | 0 | 0 | 0 | 0 | 0 | 0 | 0 | 0 | 0 |
| $tWC$ | 0 | 0 | 0 | 0 | 0 | 0 | 0 | 0 | 0 | 0 | 0 | 0 | 0 | 0 | 0 | 0 | 0 | 0 | 0 |
| $TWC$ | 0 | 0 | 0 | 0 | 0 | 0 | 0 | 0 | 0 | 0 | 0 | 0 | 0 | 0 | 0 | 0 | 0 | 0 | 0 |

$$
P_{TW} = \left(\begin{array}{c|ccccccccccccccccccc}
 & T & W & C & Tw & Tc & Wt & Wc & Ct & Cw & TW & TC & WC & Twc & tWc & twC & TWc & TwC & tWC & TWC \\
\hline
T & 0 & \frac{p_b}{N_{s,TW}} & 0 & 0 & 0 & 0 & 0 & 0 & 0 & \frac{p_s}{N_{s,TW}} & 0 & 0 & 0 & 0 & 0 & 0 & 0 & 0 & 0 \\
W & \frac{p_b}{N_{s,TW}} & 0 & 0 & 0 & 0 & 0 & 0 & 0 & 0 & \frac{p_s}{N_{s,TW}} & 0 & 0 & 0 & 0 & 0 & 0 & 0 & 0 & 0 \\
C & 0 & 0 & 0 & 0 & 0 & 0 & 0 & 0 & 0 & 0 & 0 & 0 & 0 & 0 & 0 & 0 & 0 & 0 & 0 \\
Tw & 0 & 0 & 0 & 0 & 0 & \frac{p_b}{N_{b,TW}} & 0 & 0 & 0 & \frac{p_s}{N_{s,TW}} & 0 & 0 & 0 & 0 & 0 & 0 & 0 & 0 & 0 \\
Tc & 0 & 0 & 0 & 0 & 0 & 0 & 0 & 0 & 0 & 0 & 0 & 0 & 0 & 0 & 0 & 0 & 0 & 0 & 0 \\
Wt & 0 & 0 & 0 & \frac{p_b}{N_{b,TW}} & 0 & 0 & 0 & 0 & 0 & \frac{p_s}{N_{s,TW}} & 0 & 0 & 0 & 0 & 0 & 0 & 0 & 0 & 0 \\
Wc & 0 & 0 & 0 & 0 & 0 & 0 & 0 & 0 & 0 & 0 & 0 & 0 & 0 & 0 & 0 & 0 & 0 & 0 & 0 \\
Ct & 0 & 0 & 0 & 0 & 0 & 0 & 0 & 0 & 0 & 0 & 0 & 0 & 0 & 0 & 0 & 0 & 0 & 0 & 0 \\
Cw & 0 & 0 & 0 & 0 & 0 & 0 & 0 & 0 & 0 & 0 & 0 & 0 & 0 & 0 & 0 & 0 & 0 & 0 & 0 \\
TW & \frac{p_s}{N_{s,TW}} & \frac{p_s}{N_{s,TW}} & 0 & \frac{p_s}{N_{s,TW}} & 0 & \frac{p_s}{N_{s,TW}} & 0 & 0 & 0 & \frac{p_e}{N_{e,TW}} & 0 & 0 & 0 & 0 & 0 & 0 & 0 & 0 & 0 \\
TC & 0 & 0 & 0 & 0 & 0 & 0 & 0 & 0 & 0 & 0 & 0 & 0 & 0 & 0 & 0 & 0 & 0 & 0 & 0 \\
WC & 0 & 0 & 0 & 0 & 0 & 0 & 0 & 0 & 0 & 0 & 0 & 0 & 0 & 0 & 0 & 0 & 0 & 0 & 0 \\
Twc & 0 & 0 & 0 & 0 & 0 & 0 & 0 & 0 & 0 & 0 & 0 & 0 & 0 & 0 & 0 & 0 & 0 & 0 & 0 \\
tWc & 0 & 0 & 0 & 0 & 0 & 0 & 0 & 0 & 0 & 0 & 0 & 0 & 0 & 0 & 0 & 0 & 0 & 0 & 0 \\
twC & 0 & 0 & 0 & 0 & 0 & 0 & 0 & 0 & 0 & 0 & 0 & 0 & 0 & 0 & 0 & 0 & 0 & 0 & 0 \\
TWc & 0 & 0 & 0 & 0 & 0 & 0 & 0 & 0 & 0 & 0 & 0 & 0 & 0 & 0 & 0 & 0 & 0 & 0 & 0 \\
TwC & 0 & 0 & 0 & 0 & 0 & 0 & 0 & 0 & 0 & 0 & 0 & 0 & 0 & 0 & 0 & 0 & 0 & 0 & 0 \\
tWC & 0 & 0 & 0 & 0 & 0 & 0 & 0 & 0 & 0 & 0 & 0 & 0 & 0 & 0 & 0 & 0 & 0 & 0 & 0 \\
TWC & 0 & 0 & 0 & 0 & 0 & 0 & 0 & 0 & 0 & 0 & 0 & 0 & 0 & 0 & 0 & 0 & 0 & 0 & 0
\end{array}\right)
$$

$$P_{TC} =$$

| | $T$ | $W$ | $C$ | $Tw$ | $Tc$ | $Wt$ | $Wc$ | $Ct$ | $Cw$ | $TW$ | $TC$ | $WC$ | $Twc$ | $tWc$ | $twC$ | $TWc$ | $TwC$ | $tWC$ | $TWC$ |
|---|---|---|---|---|---|---|---|---|---|---|---|---|---|---|---|---|---|---|---|
| $T$ | 0 | 0 | $\frac{p_b}{N_{b,TC}}$ | 0 | 0 | 0 | 0 | 0 | 0 | 0 | $\frac{p_s}{N_{s,TC}}$ | 0 | 0 | 0 | 0 | 0 | 0 | 0 | 0 |
| $W$ | 0 | 0 | 0 | 0 | 0 | 0 | 0 | 0 | 0 | 0 | 0 | 0 | 0 | 0 | 0 | 0 | 0 | 0 | 0 |
| $C$ | $\frac{p_b}{N_{b,TC}}$ | 0 | 0 | 0 | 0 | 0 | 0 | 0 | 0 | 0 | $\frac{p_s}{N_{s,TC}}$ | 0 | 0 | 0 | 0 | 0 | 0 | 0 | 0 |
| $Tw$ | 0 | 0 | 0 | 0 | 0 | 0 | 0 | 0 | 0 | 0 | 0 | 0 | 0 | 0 | 0 | 0 | 0 | 0 | 0 |
| $Tc$ | 0 | 0 | 0 | 0 | 0 | 0 | 0 | $\frac{p_b}{N_{b,TC}}$ | 0 | 0 | $\frac{p_s}{N_{s,TC}}$ | 0 | 0 | 0 | 0 | 0 | 0 | 0 | 0 |
| $Wt$ | 0 | 0 | 0 | 0 | 0 | 0 | 0 | 0 | 0 | 0 | 0 | 0 | 0 | 0 | 0 | 0 | 0 | 0 | 0 |
| $Wc$ | 0 | 0 | 0 | 0 | 0 | 0 | 0 | 0 | 0 | 0 | 0 | 0 | 0 | 0 | 0 | 0 | 0 | 0 | 0 |
| $Ct$ | 0 | 0 | 0 | 0 | $\frac{p_b}{N_{b,TC}}$ | 0 | 0 | 0 | 0 | 0 | $\frac{p_s}{N_{s,TC}}$ | 0 | 0 | 0 | 0 | 0 | 0 | 0 | 0 |
| $Cw$ | 0 | 0 | 0 | 0 | 0 | 0 | 0 | 0 | 0 | 0 | 0 | 0 | 0 | 0 | 0 | 0 | 0 | 0 | 0 |
| $TW$ | 0 | 0 | 0 | 0 | 0 | 0 | 0 | 0 | 0 | 0 | 0 | 0 | 0 | 0 | 0 | 0 | 0 | 0 | 0 |
| $TC$ | $\frac{p_s}{N_{s,TC}}$ | 0 | $\frac{p_s}{N_{s,TC}}$ | 0 | $\frac{p_s}{N_{s,TC}}$ | 0 | 0 | $\frac{p_s}{N_{s,TC}}$ | 0 | 0 | $\frac{p_e}{N_{e,TC}}$ | 0 | 0 | 0 | 0 | 0 | 0 | 0 | 0 |
| $WC$ | 0 | 0 | 0 | 0 | 0 | 0 | 0 | 0 | 0 | 0 | 0 | 0 | 0 | 0 | 0 | 0 | 0 | 0 | 0 |
| $Twc$ | 0 | 0 | 0 | 0 | 0 | 0 | 0 | 0 | 0 | 0 | 0 | 0 | 0 | 0 | 0 | 0 | 0 | 0 | 0 |
| $tWc$ | 0 | 0 | 0 | 0 | 0 | 0 | 0 | 0 | 0 | 0 | 0 | 0 | 0 | 0 | 0 | 0 | 0 | 0 | 0 |
| $twC$ | 0 | 0 | 0 | 0 | 0 | 0 | 0 | 0 | 0 | 0 | 0 | 0 | 0 | 0 | 0 | 0 | 0 | 0 | 0 |
| $TWc$ | 0 | 0 | 0 | 0 | 0 | 0 | 0 | 0 | 0 | 0 | 0 | 0 | 0 | 0 | 0 | 0 | 0 | 0 | 0 |
| $TwC$ | 0 | 0 | 0 | 0 | 0 | 0 | 0 | 0 | 0 | 0 | 0 | 0 | 0 | 0 | 0 | 0 | 0 | 0 | 0 |
| $tWC$ | 0 | 0 | 0 | 0 | 0 | 0 | 0 | 0 | 0 | 0 | 0 | 0 | 0 | 0 | 0 | 0 | 0 | 0 | 0 |
| $TWC$ | 0 | 0 | 0 | 0 | 0 | 0 | 0 | 0 | 0 | 0 | 0 | 0 | 0 | 0 | 0 | 0 | 0 | 0 | 0 |

$$P_{WC} = \left(\begin{array}{c|ccccccccccccccccccc}
 & T & W & C & Tw & Tc & Wt & Wc & Ct & Cw & TW & TC & WC & Twc & tWc & twC & TWc & TwC & tWC & TWC \\
\hline
T & 0 & 0 & 0 & 0 & 0 & 0 & 0 & 0 & 0 & 0 & 0 & 0 & 0 & 0 & 0 & 0 & 0 & 0 & 0 \\
W & 0 & 0 & \frac{p_b}{N_{b,WC}} & 0 & 0 & 0 & 0 & 0 & 0 & 0 & 0 & \frac{p_s}{N_{s,WC}} & 0 & 0 & 0 & 0 & 0 & 0 & 0 \\
C & 0 & \frac{p_b}{N_{b,WC}} & 0 & 0 & 0 & 0 & 0 & 0 & 0 & 0 & 0 & \frac{p_s}{N_{s,WC}} & 0 & 0 & 0 & 0 & 0 & 0 & 0 \\
Tw & 0 & 0 & 0 & 0 & 0 & 0 & 0 & 0 & 0 & 0 & 0 & 0 & 0 & 0 & 0 & 0 & 0 & 0 & 0 \\
Tc & 0 & 0 & 0 & 0 & 0 & 0 & 0 & 0 & 0 & 0 & 0 & 0 & 0 & 0 & 0 & 0 & 0 & 0 & 0 \\
Wt & 0 & 0 & 0 & 0 & 0 & 0 & 0 & 0 & 0 & 0 & 0 & 0 & 0 & 0 & 0 & 0 & 0 & 0 & 0 \\
Wc & 0 & 0 & 0 & 0 & 0 & 0 & 0 & 0 & \frac{p_b}{N_{b,WC}} & 0 & 0 & \frac{p_s}{N_{s,WC}} & 0 & 0 & 0 & 0 & 0 & 0 & 0 \\
Ct & 0 & 0 & 0 & 0 & 0 & 0 & 0 & 0 & 0 & 0 & 0 & 0 & 0 & 0 & 0 & 0 & 0 & 0 & 0 \\
Cw & 0 & 0 & 0 & 0 & 0 & 0 & \frac{p_b}{N_{b,WC}} & 0 & 0 & 0 & 0 & \frac{p_s}{N_{s,WC}} & 0 & 0 & 0 & 0 & 0 & 0 & 0 \\
TW & 0 & 0 & 0 & 0 & 0 & 0 & 0 & 0 & 0 & 0 & 0 & 0 & 0 & 0 & 0 & 0 & 0 & 0 & 0 \\
TC & 0 & 0 & 0 & 0 & 0 & 0 & 0 & 0 & 0 & 0 & 0 & 0 & 0 & 0 & 0 & 0 & 0 & 0 & 0 \\
WC & 0 & \frac{p_s}{N_{s,WC}} & \frac{p_s}{N_{s,WC}} & 0 & 0 & 0 & \frac{p_s}{N_{s,WC}} & 0 & \frac{p_s}{N_{s,WC}} & 0 & 0 & \frac{p_e}{N_{e,WC}} & 0 & 0 & 0 & 0 & 0 & 0 & 0 \\
Twc & 0 & 0 & 0 & 0 & 0 & 0 & 0 & 0 & 0 & 0 & 0 & 0 & 0 & 0 & 0 & 0 & 0 & 0 & 0 \\
tWc & 0 & 0 & 0 & 0 & 0 & 0 & 0 & 0 & 0 & 0 & 0 & 0 & 0 & 0 & 0 & 0 & 0 & 0 & 0 \\
twC & 0 & 0 & 0 & 0 & 0 & 0 & 0 & 0 & 0 & 0 & 0 & 0 & 0 & 0 & 0 & 0 & 0 & 0 & 0 \\
TWc & 0 & 0 & 0 & 0 & 0 & 0 & 0 & 0 & 0 & 0 & 0 & 0 & 0 & 0 & 0 & 0 & 0 & 0 & 0 \\
TwC & 0 & 0 & 0 & 0 & 0 & 0 & 0 & 0 & 0 & 0 & 0 & 0 & 0 & 0 & 0 & 0 & 0 & 0 & 0 \\
tWC & 0 & 0 & 0 & 0 & 0 & 0 & 0 & 0 & 0 & 0 & 0 & 0 & 0 & 0 & 0 & 0 & 0 & 0 & 0 \\
TWC & 0 & 0 & 0 & 0 & 0 & 0 & 0 & 0 & 0 & 0 & 0 & 0 & 0 & 0 & 0 & 0 & 0 & 0 & 0
\end{array}\right)$$

$$P_{Twc} = \left(\begin{array}{c|ccccccccccccccccccc}
 & T & W & C & Tw & Tc & Wt & Wc & Ct & Cw & TW & TC & WC & Twc & tWc & twC & TWc & TwC & tWC & TWC \\
\hline
T & 0 & 0 & 0 & 0 & 0 & 0 & 0 & 0 & 0 & 0 & 0 & 0 & 0 & 0 & 0 & 0 & 0 & 0 & 0 \\
W & 0 & 0 & 0 & 0 & 0 & 0 & 0 & 0 & 0 & 0 & 0 & 0 & 0 & 0 & 0 & 0 & 0 & 0 & 0 \\
C & 0 & 0 & 0 & 0 & 0 & 0 & 0 & 0 & 0 & 0 & 0 & 0 & 0 & 0 & 0 & 0 & 0 & 0 & 0 \\
Tw & 0 & 0 & 0 & 0 & 0 & 0 & 0 & 0 & 0 & 0 & 0 & 0 & 0 & 0 & 0 & 0 & 0 & 0 & 0 \\
Tc & 0 & 0 & 0 & 0 & 0 & 0 & 0 & 0 & 0 & 0 & 0 & 0 & 0 & 0 & 0 & 0 & 0 & 0 & 0 \\
Wt & 0 & 0 & 0 & 0 & 0 & 0 & 0 & 0 & 0 & 0 & 0 & 0 & 0 & 0 & 0 & 0 & 0 & 0 & 0 \\
Wc & 0 & 0 & 0 & 0 & 0 & 0 & 0 & 0 & 0 & 0 & 0 & 0 & 0 & 0 & 0 & 0 & 0 & 0 & 0 \\
Ct & 0 & 0 & 0 & 0 & 0 & 0 & 0 & 0 & 0 & 0 & 0 & 0 & 0 & 0 & 0 & 0 & 0 & 0 & 0 \\
Cw & 0 & 0 & 0 & 0 & 0 & 0 & 0 & 0 & 0 & 0 & 0 & 0 & 0 & 0 & 0 & 0 & 0 & 0 & 0 \\
TW & 0 & 0 & 0 & 0 & 0 & 0 & 0 & 0 & 0 & 0 & 0 & 0 & 0 & 0 & 0 & 0 & 0 & 0 & 0 \\
TC & 0 & 0 & 0 & 0 & 0 & 0 & 0 & 0 & 0 & 0 & 0 & 0 & 0 & 0 & 0 & 0 & 0 & 0 & 0 \\
WC & 0 & 0 & 0 & 0 & 0 & 0 & 0 & 0 & 0 & 0 & 0 & 0 & 0 & 0 & 0 & 0 & 0 & 0 & 0 \\
Twc & 0 & 0 & 0 & 0 & 0 & 0 & 0 & 0 & 0 & 0 & 0 & 0 & 1 & 0 & 0 & 0 & 0 & 0 & 0 \\
tWc & 0 & 0 & 0 & 0 & 0 & 0 & 0 & 0 & 0 & 0 & 0 & 0 & 0 & 0 & 0 & 0 & 0 & 0 & 0 \\
twC & 0 & 0 & 0 & 0 & 0 & 0 & 0 & 0 & 0 & 0 & 0 & 0 & 0 & 0 & 0 & 0 & 0 & 0 & 0 \\
TWc & 0 & 0 & 0 & 0 & 0 & 0 & 0 & 0 & 0 & 0 & 0 & 0 & 0 & 0 & 0 & 0 & 0 & 0 & 0 \\
TwC & 0 & 0 & 0 & 0 & 0 & 0 & 0 & 0 & 0 & 0 & 0 & 0 & 0 & 0 & 0 & 0 & 0 & 0 & 0 \\
tWC & 0 & 0 & 0 & 0 & 0 & 0 & 0 & 0 & 0 & 0 & 0 & 0 & 0 & 0 & 0 & 0 & 0 & 0 & 0 \\
TWC & 0 & 0 & 0 & 0 & 0 & 0 & 0 & 0 & 0 & 0 & 0 & 0 & 0 & 0 & 0 & 0 & 0 & 0 & 0
\end{array}\right)$$

$$P_{tWc} =$$

| | $T$ | $W$ | $C$ | $Tw$ | $Tc$ | $Wt$ | $Wc$ | $Ct$ | $Cw$ | $TW$ | $TC$ | $WC$ | $Twc$ | $tWc$ | $twC$ | $TWc$ | $TwC$ | $tWC$ | $TWC$ |
|---|---|---|---|---|---|---|---|---|---|---|---|---|---|---|---|---|---|---|---|
| $T$ | 0 | 0 | 0 | 0 | 0 | 0 | 0 | 0 | 0 | 0 | 0 | 0 | 0 | 0 | 0 | 0 | 0 | 0 | 0 |
| $W$ | 0 | 0 | 0 | 0 | 0 | 0 | 0 | 0 | 0 | 0 | 0 | 0 | 0 | 0 | 0 | 0 | 0 | 0 | 0 |
| $C$ | 0 | 0 | 0 | 0 | 0 | 0 | 0 | 0 | 0 | 0 | 0 | 0 | 0 | 0 | 0 | 0 | 0 | 0 | 0 |
| $Tw$ | 0 | 0 | 0 | 0 | 0 | 0 | 0 | 0 | 0 | 0 | 0 | 0 | 0 | 0 | 0 | 0 | 0 | 0 | 0 |
| $Tc$ | 0 | 0 | 0 | 0 | 0 | 0 | 0 | 0 | 0 | 0 | 0 | 0 | 0 | 0 | 0 | 0 | 0 | 0 | 0 |
| $Wt$ | 0 | 0 | 0 | 0 | 0 | 0 | 0 | 0 | 0 | 0 | 0 | 0 | 0 | 0 | 0 | 0 | 0 | 0 | 0 |
| $Wc$ | 0 | 0 | 0 | 0 | 0 | 0 | 0 | 0 | 0 | 0 | 0 | 0 | 0 | 0 | 0 | 0 | 0 | 0 | 0 |
| $Ct$ | 0 | 0 | 0 | 0 | 0 | 0 | 0 | 0 | 0 | 0 | 0 | 0 | 0 | 0 | 0 | 0 | 0 | 0 | 0 |
| $Cw$ | 0 | 0 | 0 | 0 | 0 | 0 | 0 | 0 | 0 | 0 | 0 | 0 | 0 | 0 | 0 | 0 | 0 | 0 | 0 |
| $TW$ | 0 | 0 | 0 | 0 | 0 | 0 | 0 | 0 | 0 | 0 | 0 | 0 | 0 | 0 | 0 | 0 | 0 | 0 | 0 |
| $TC$ | 0 | 0 | 0 | 0 | 0 | 0 | 0 | 0 | 0 | 0 | 0 | 0 | 0 | 0 | 0 | 0 | 0 | 0 | 0 |
| $WC$ | 0 | 0 | 0 | 0 | 0 | 0 | 0 | 0 | 0 | 0 | 0 | 0 | 0 | 0 | 0 | 0 | 0 | 0 | 0 |
| $Twc$ | 0 | 0 | 0 | 0 | 0 | 0 | 0 | 0 | 0 | 0 | 0 | 0 | 0 | 0 | 0 | 0 | 0 | 0 | 0 |
| $tWc$ | 0 | 0 | 0 | 0 | 0 | 0 | 0 | 0 | 0 | 0 | 0 | 0 | 0 | 1 | 0 | 0 | 0 | 0 | 0 |
| $twC$ | 0 | 0 | 0 | 0 | 0 | 0 | 0 | 0 | 0 | 0 | 0 | 0 | 0 | 0 | 0 | 0 | 0 | 0 | 0 |
| $TWc$ | 0 | 0 | 0 | 0 | 0 | 0 | 0 | 0 | 0 | 0 | 0 | 0 | 0 | 0 | 0 | 0 | 0 | 0 | 0 |
| $TwC$ | 0 | 0 | 0 | 0 | 0 | 0 | 0 | 0 | 0 | 0 | 0 | 0 | 0 | 0 | 0 | 0 | 0 | 0 | 0 |
| $tWC$ | 0 | 0 | 0 | 0 | 0 | 0 | 0 | 0 | 0 | 0 | 0 | 0 | 0 | 0 | 0 | 0 | 0 | 0 | 0 |
| $TWC$ | 0 | 0 | 0 | 0 | 0 | 0 | 0 | 0 | 0 | 0 | 0 | 0 | 0 | 0 | 0 | 0 | 0 | 0 | 0 |

$$P_{twC} =$$

| | $T$ | $W$ | $C$ | $Tw$ | $Tc$ | $Wt$ | $Wc$ | $Ct$ | $Cw$ | $TW$ | $TC$ | $WC$ | $Twc$ | $tWc$ | $twC$ | $TWc$ | $TwC$ | $tWC$ | $TWC$ |
|---|---|---|---|---|---|---|---|---|---|---|---|---|---|---|---|---|---|---|---|
| $T$ | 0 | 0 | 0 | 0 | 0 | 0 | 0 | 0 | 0 | 0 | 0 | 0 | 0 | 0 | 0 | 0 | 0 | 0 | 0 |
| $W$ | 0 | 0 | 0 | 0 | 0 | 0 | 0 | 0 | 0 | 0 | 0 | 0 | 0 | 0 | 0 | 0 | 0 | 0 | 0 |
| $C$ | 0 | 0 | 0 | 0 | 0 | 0 | 0 | 0 | 0 | 0 | 0 | 0 | 0 | 0 | 0 | 0 | 0 | 0 | 0 |
| $Tw$ | 0 | 0 | 0 | 0 | 0 | 0 | 0 | 0 | 0 | 0 | 0 | 0 | 0 | 0 | 0 | 0 | 0 | 0 | 0 |
| $Tc$ | 0 | 0 | 0 | 0 | 0 | 0 | 0 | 0 | 0 | 0 | 0 | 0 | 0 | 0 | 0 | 0 | 0 | 0 | 0 |
| $Wt$ | 0 | 0 | 0 | 0 | 0 | 0 | 0 | 0 | 0 | 0 | 0 | 0 | 0 | 0 | 0 | 0 | 0 | 0 | 0 |
| $Wc$ | 0 | 0 | 0 | 0 | 0 | 0 | 0 | 0 | 0 | 0 | 0 | 0 | 0 | 0 | 0 | 0 | 0 | 0 | 0 |
| $Ct$ | 0 | 0 | 0 | 0 | 0 | 0 | 0 | 0 | 0 | 0 | 0 | 0 | 0 | 0 | 0 | 0 | 0 | 0 | 0 |
| $Cw$ | 0 | 0 | 0 | 0 | 0 | 0 | 0 | 0 | 0 | 0 | 0 | 0 | 0 | 0 | 0 | 0 | 0 | 0 | 0 |
| $TW$ | 0 | 0 | 0 | 0 | 0 | 0 | 0 | 0 | 0 | 0 | 0 | 0 | 0 | 0 | 0 | 0 | 0 | 0 | 0 |
| $TC$ | 0 | 0 | 0 | 0 | 0 | 0 | 0 | 0 | 0 | 0 | 0 | 0 | 0 | 0 | 0 | 0 | 0 | 0 | 0 |
| $WC$ | 0 | 0 | 0 | 0 | 0 | 0 | 0 | 0 | 0 | 0 | 0 | 0 | 0 | 0 | 0 | 0 | 0 | 0 | 0 |
| $Twc$ | 0 | 0 | 0 | 0 | 0 | 0 | 0 | 0 | 0 | 0 | 0 | 0 | 0 | 0 | 0 | 0 | 0 | 0 | 0 |
| $tWc$ | 0 | 0 | 0 | 0 | 0 | 0 | 0 | 0 | 0 | 0 | 0 | 0 | 0 | 0 | 0 | 0 | 0 | 0 | 0 |
| $twC$ | 0 | 0 | 0 | 0 | 0 | 0 | 0 | 0 | 0 | 0 | 0 | 0 | 0 | 0 | 1 | 0 | 0 | 0 | 0 |
| $TWc$ | 0 | 0 | 0 | 0 | 0 | 0 | 0 | 0 | 0 | 0 | 0 | 0 | 0 | 0 | 0 | 0 | 0 | 0 | 0 |
| $TwC$ | 0 | 0 | 0 | 0 | 0 | 0 | 0 | 0 | 0 | 0 | 0 | 0 | 0 | 0 | 0 | 0 | 0 | 0 | 0 |
| $tWC$ | 0 | 0 | 0 | 0 | 0 | 0 | 0 | 0 | 0 | 0 | 0 | 0 | 0 | 0 | 0 | 0 | 0 | 0 | 0 |
| $TWC$ | 0 | 0 | 0 | 0 | 0 | 0 | 0 | 0 | 0 | 0 | 0 | 0 | 0 | 0 | 0 | 0 | 0 | 0 | 0 |

$$
P_{TWc} = \left(\begin{array}{c|ccccccccccccccccccc}
 & T & W & C & Tw & Tc & Wt & Wc & Ct & Cw & TW & TC & WC & Twc & tWc & twC & TWc & TwC & tWC & TWC \\
\hline
T & 0 & 0 & 0 & 0 & 0 & 0 & 0 & 0 & 0 & 0 & 0 & 0 & 0 & 0 & 0 & 0 & 0 & 0 & 0 \\
W & 0 & 0 & 0 & 0 & 0 & 0 & 0 & 0 & 0 & 0 & 0 & 0 & 0 & 0 & 0 & 0 & 0 & 0 & 0 \\
C & 0 & 0 & 0 & 0 & 0 & 0 & 0 & 0 & 0 & 0 & 0 & 0 & 0 & 0 & 0 & 0 & 0 & 0 & 0 \\
Tw & 0 & 0 & 0 & 0 & 0 & 0 & 0 & 0 & 0 & 0 & 0 & 0 & 0 & 0 & 0 & 0 & 0 & 0 & 0 \\
Tc & 0 & 0 & 0 & 0 & 0 & 0 & \frac{p_b}{N_{b,TWc}} & 0 & 0 & 0 & 0 & 0 & 0 & 0 & 0 & \frac{p_s}{N_{s,TWc}} & 0 & 0 & 0 \\
Wt & 0 & 0 & 0 & 0 & 0 & 0 & 0 & 0 & 0 & 0 & 0 & 0 & 0 & 0 & 0 & 0 & 0 & 0 & 0 \\
Wc & 0 & 0 & 0 & 0 & \frac{p_b}{N_{b,TWc}} & 0 & 0 & 0 & 0 & 0 & 0 & 0 & 0 & 0 & 0 & \frac{p_s}{N_{s,TWc}} & 0 & 0 & 0 \\
Ct & 0 & 0 & 0 & 0 & 0 & 0 & 0 & 0 & 0 & 0 & 0 & 0 & 0 & 0 & 0 & 0 & 0 & 0 & 0 \\
Cw & 0 & 0 & 0 & 0 & 0 & 0 & 0 & 0 & 0 & 0 & 0 & 0 & 0 & 0 & 0 & 0 & 0 & 0 & 0 \\
TW & 0 & 0 & 0 & 0 & 0 & 0 & 0 & 0 & 0 & 0 & 0 & 0 & 0 & 0 & 0 & 0 & 0 & 0 & 0 \\
TC & 0 & 0 & 0 & 0 & 0 & 0 & 0 & 0 & 0 & 0 & 0 & 0 & 0 & 0 & 0 & 0 & 0 & 0 & 0 \\
WC & 0 & 0 & 0 & 0 & 0 & 0 & 0 & 0 & 0 & 0 & 0 & 0 & 0 & 0 & 0 & 0 & 0 & 0 & 0 \\
Twc & 0 & 0 & 0 & 0 & 0 & 0 & 0 & 0 & 0 & 0 & 0 & 0 & 0 & \frac{p_b}{N_{b,TWc}} & 0 & \frac{p_s}{N_{s,TWc}} & 0 & 0 & 0 \\
tWc & 0 & 0 & 0 & 0 & 0 & 0 & 0 & 0 & 0 & 0 & 0 & 0 & \frac{p_b}{N_{b,TWc}} & 0 & 0 & \frac{p_s}{N_{s,TWc}} & 0 & 0 & 0 \\
twC & 0 & 0 & 0 & 0 & 0 & 0 & 0 & 0 & 0 & 0 & 0 & 0 & 0 & 0 & 0 & 0 & 0 & 0 & 0 \\
TWc & 0 & 0 & 0 & 0 & \frac{p_s}{N_{s,TWc}} & 0 & \frac{p_s}{N_{s,TWc}} & 0 & 0 & 0 & 0 & 0 & \frac{p_s}{N_{s,TWc}} & \frac{p_s}{N_{s,TWc}} & 0 & \frac{p_e}{N_{e,TWc}} & 0 & 0 & 0 \\
TwC & 0 & 0 & 0 & 0 & 0 & 0 & 0 & 0 & 0 & 0 & 0 & 0 & 0 & 0 & 0 & 0 & 0 & 0 & 0 \\
tWC & 0 & 0 & 0 & 0 & 0 & 0 & 0 & 0 & 0 & 0 & 0 & 0 & 0 & 0 & 0 & 0 & 0 & 0 & 0 \\
TWC & 0 & 0 & 0 & 0 & 0 & 0 & 0 & 0 & 0 & 0 & 0 & 0 & 0 & 0 & 0 & 0 & 0 & 0 & 0
\end{array}\right)
$$

$$P_{TwC} =$$

| | $T$ | $W$ | $C$ | $Tw$ | $Tc$ | $Wt$ | $Wc$ | $Ct$ | $Cw$ | $TW$ | $TC$ | $WC$ | $Twc$ | $tWc$ | $twC$ | $TWc$ | $TwC$ | $tWC$ | $TWC$ |
|---|---|---|---|---|---|---|---|---|---|---|---|---|---|---|---|---|---|---|---|
| $T$ | 0 | 0 | 0 | 0 | 0 | 0 | 0 | 0 | 0 | 0 | 0 | 0 | 0 | 0 | 0 | 0 | 0 | 0 | 0 |
| $W$ | 0 | 0 | 0 | 0 | 0 | 0 | 0 | 0 | 0 | 0 | 0 | 0 | 0 | 0 | 0 | 0 | 0 | 0 | 0 |
| $C$ | 0 | 0 | 0 | 0 | 0 | 0 | 0 | 0 | 0 | 0 | 0 | 0 | 0 | 0 | 0 | 0 | 0 | 0 | 0 |
| $Tw$ | 0 | 0 | 0 | 0 | 0 | 0 | 0 | 0 | $\frac{p_b}{N_{b,TwC}}$ | 0 | 0 | 0 | 0 | 0 | 0 | 0 | $\frac{p_s}{N_{s,TwC}}$ | 0 | 0 |
| $Tc$ | 0 | 0 | 0 | 0 | 0 | 0 | 0 | 0 | 0 | 0 | 0 | 0 | 0 | 0 | 0 | 0 | 0 | 0 | 0 |
| $Wt$ | 0 | 0 | 0 | 0 | 0 | 0 | 0 | 0 | 0 | 0 | 0 | 0 | 0 | 0 | 0 | 0 | 0 | 0 | 0 |
| $Wc$ | 0 | 0 | 0 | 0 | 0 | 0 | 0 | 0 | 0 | 0 | 0 | 0 | 0 | 0 | 0 | 0 | 0 | 0 | 0 |
| $Ct$ | 0 | 0 | 0 | 0 | 0 | 0 | 0 | 0 | 0 | 0 | 0 | 0 | 0 | 0 | 0 | 0 | 0 | 0 | 0 |
| $Cw$ | 0 | 0 | 0 | $\frac{p_b}{N_{b,TwC}}$ | 0 | 0 | 0 | 0 | 0 | 0 | 0 | 0 | 0 | 0 | 0 | 0 | $\frac{p_s}{N_{s,TwC}}$ | 0 | 0 |
| $TW$ | 0 | 0 | 0 | 0 | 0 | 0 | 0 | 0 | 0 | 0 | 0 | 0 | 0 | 0 | 0 | 0 | 0 | 0 | 0 |
| $TC$ | 0 | 0 | 0 | 0 | 0 | 0 | 0 | 0 | 0 | 0 | 0 | 0 | 0 | 0 | 0 | 0 | 0 | 0 | 0 |
| $WC$ | 0 | 0 | 0 | 0 | 0 | 0 | 0 | 0 | 0 | 0 | 0 | 0 | 0 | 0 | 0 | 0 | 0 | 0 | 0 |
| $Twc$ | 0 | 0 | 0 | 0 | 0 | 0 | 0 | 0 | 0 | 0 | 0 | 0 | 0 | 0 | $\frac{p_b}{N_{b,TwC}}$ | 0 | $\frac{p_s}{N_{s,TwC}}$ | 0 | 0 |
| $tWc$ | 0 | 0 | 0 | 0 | 0 | 0 | 0 | 0 | 0 | 0 | 0 | 0 | 0 | 0 | 0 | 0 | 0 | 0 | 0 |
| $twC$ | 0 | 0 | 0 | 0 | 0 | 0 | 0 | 0 | 0 | 0 | 0 | 0 | $\frac{p_b}{N_{b,TwC}}$ | 0 | 0 | 0 | $\frac{p_s}{N_{s,TwC}}$ | 0 | 0 |
| $TWc$ | 0 | 0 | 0 | 0 | 0 | 0 | 0 | 0 | 0 | 0 | 0 | 0 | 0 | 0 | 0 | 0 | 0 | 0 | 0 |
| $TwC$ | 0 | 0 | 0 | $\frac{p_s}{N_{s,TwC}}$ | 0 | 0 | 0 | 0 | $\frac{p_s}{N_{s,TwC}}$ | 0 | 0 | 0 | $\frac{p_s}{N_{s,TwC}}$ | 0 | $\frac{p_s}{N_{s,TwC}}$ | 0 | $\frac{p_e}{N_{e,TwC}}$ | 0 | 0 |
| $tWC$ | 0 | 0 | 0 | 0 | 0 | 0 | 0 | 0 | 0 | 0 | 0 | 0 | 0 | 0 | 0 | 0 | 0 | 0 | 0 |
| $TWC$ | 0 | 0 | 0 | 0 | 0 | 0 | 0 | 0 | 0 | 0 | 0 | 0 | 0 | 0 | 0 | 0 | 0 | 0 | 0 |

$$
P_{tWC} = \left(\begin{array}{c|ccccccccccccccccccc}
 & T & W & C & Tw & Tc & Wt & Wc & Ct & Cw & TW & TC & WC & Twc & tWc & twC & TWc & TwC & tWC & TWC \\
\hline
T & 0 & 0 & 0 & 0 & 0 & 0 & 0 & 0 & 0 & 0 & 0 & 0 & 0 & 0 & 0 & 0 & 0 & 0 & 0 \\
W & 0 & 0 & 0 & 0 & 0 & 0 & 0 & 0 & 0 & 0 & 0 & 0 & 0 & 0 & 0 & 0 & 0 & 0 & 0 \\
C & 0 & 0 & 0 & 0 & 0 & 0 & 0 & 0 & 0 & 0 & 0 & 0 & 0 & 0 & 0 & 0 & 0 & 0 & 0 \\
Tw & 0 & 0 & 0 & 0 & 0 & 0 & 0 & 0 & 0 & 0 & 0 & 0 & 0 & 0 & 0 & 0 & 0 & \frac{p_s}{N_{s,tWC}} & 0 \\
Tc & 0 & 0 & 0 & 0 & 0 & 0 & 0 & 0 & 0 & 0 & 0 & 0 & 0 & 0 & 0 & 0 & 0 & 0 & 0 \\
Wt & 0 & 0 & 0 & 0 & 0 & 0 & 0 & \frac{p_b}{N_{b,tWC}} & 0 & 0 & 0 & 0 & 0 & 0 & 0 & 0 & 0 & \frac{p_s}{N_{s,tWC}} & 0 \\
Wc & 0 & 0 & 0 & 0 & 0 & 0 & 0 & 0 & 0 & 0 & 0 & 0 & 0 & 0 & 0 & 0 & 0 & 0 & 0 \\
Ct & 0 & 0 & 0 & 0 & 0 & \frac{p_b}{N_{b,tWC}} & 0 & 0 & 0 & 0 & 0 & 0 & 0 & 0 & 0 & 0 & 0 & 0 & 0 \\
Cw & 0 & 0 & 0 & 0 & 0 & 0 & 0 & 0 & 0 & 0 & 0 & 0 & 0 & 0 & 0 & 0 & 0 & 0 & 0 \\
TW & 0 & 0 & 0 & 0 & 0 & 0 & 0 & 0 & 0 & 0 & 0 & 0 & 0 & 0 & 0 & 0 & 0 & 0 & 0 \\
TC & 0 & 0 & 0 & 0 & 0 & 0 & 0 & 0 & 0 & 0 & 0 & 0 & 0 & 0 & 0 & 0 & 0 & 0 & 0 \\
WC & 0 & 0 & 0 & 0 & 0 & 0 & 0 & 0 & 0 & 0 & 0 & 0 & 0 & 0 & 0 & 0 & 0 & 0 & 0 \\
Twc & 0 & 0 & 0 & 0 & 0 & 0 & 0 & 0 & 0 & 0 & 0 & 0 & 0 & 0 & 0 & 0 & 0 & 0 & 0 \\
tWc & 0 & 0 & 0 & 0 & 0 & 0 & 0 & 0 & 0 & 0 & 0 & 0 & 0 & 0 & \frac{p_b}{N_{b,tWC}} & 0 & 0 & \frac{p_s}{N_{s,tWC}} & 0 \\
twC & 0 & 0 & 0 & 0 & 0 & 0 & 0 & 0 & 0 & 0 & 0 & 0 & 0 & \frac{p_b}{N_{b,tWC}} & 0 & 0 & 0 & \frac{p_s}{N_{s,tWC}} & 0 \\
TWc & 0 & 0 & 0 & 0 & 0 & 0 & 0 & 0 & 0 & 0 & 0 & 0 & 0 & 0 & 0 & 0 & 0 & 0 & 0 \\
TwC & 0 & 0 & 0 & 0 & 0 & 0 & 0 & 0 & 0 & 0 & 0 & 0 & 0 & 0 & 0 & 0 & 0 & 0 & 0 \\
tWC & 0 & 0 & 0 & \frac{p_s}{N_{s,tWC}} & 0 & \frac{p_s}{N_{s,tWC}} & 0 & 0 & 0 & 0 & 0 & 0 & 0 & \frac{p_s}{N_{s,tWC}} & \frac{p_s}{N_{s,tWC}} & 0 & 0 & \frac{p_e}{N_{e,tWC}} & 0 \\
TWC & 0 & 0 & 0 & 0 & 0 & 0 & 0 & 0 & 0 & 0 & 0 & 0 & 0 & 0 & 0 & 0 & 0 & 0 & 0
\end{array}\right)
$$

$$
P_{TWC} = \left(\begin{array}{c|ccccccccccccccccccc}
 & T & W & C & Tw & Tc & Wt & Wc & Ct & Cw & TW & TC & WC & Twc & tWc & twC & TWc & TwC & tWC & TWC \\
\hline
T & 0 & 0 & 0 & 0 & 0 & 0 & 0 & 0 & 0 & 0 & 0 & \frac{p_b}{N_{b,TWC}} & 0 & 0 & 0 & 0 & 0 & 0 & \frac{p_s}{N_{s,TWC}} \\
W & 0 & 0 & 0 & 0 & 0 & 0 & 0 & 0 & 0 & 0 & \frac{p_b}{N_{b,TWC}} & 0 & 0 & 0 & 0 & 0 & 0 & 0 & \frac{p_s}{N_{s,TWC}} \\
C & 0 & 0 & 0 & 0 & 0 & 0 & 0 & 0 & 0 & \frac{p_b}{N_{b,TWC}} & 0 & 0 & 0 & 0 & 0 & 0 & 0 & 0 & \frac{p_s}{N_{s,TWC}} \\
Tw & 0 & 0 & 0 & 0 & 0 & 0 & 0 & 0 & 0 & 0 & 0 & 0 & 0 & 0 & 0 & 0 & 0 & 0 & 0 \\
Tc & 0 & 0 & 0 & 0 & 0 & 0 & 0 & 0 & 0 & 0 & 0 & 0 & 0 & 0 & 0 & 0 & 0 & 0 & 0 \\
Wt & 0 & 0 & 0 & 0 & 0 & 0 & 0 & 0 & 0 & 0 & 0 & 0 & 0 & 0 & 0 & 0 & 0 & 0 & 0 \\
Wc & 0 & 0 & 0 & 0 & 0 & 0 & 0 & 0 & 0 & 0 & 0 & 0 & 0 & 0 & 0 & 0 & 0 & 0 & 0 \\
Ct & 0 & 0 & 0 & 0 & 0 & 0 & 0 & 0 & 0 & 0 & 0 & 0 & 0 & 0 & 0 & 0 & 0 & 0 & 0 \\
Cw & 0 & 0 & 0 & 0 & 0 & 0 & 0 & 0 & 0 & 0 & 0 & 0 & 0 & 0 & 0 & 0 & 0 & 0 & 0 \\
TW & 0 & 0 & \frac{p_b}{N_{b,TWC}} & 0 & 0 & 0 & 0 & 0 & 0 & 0 & 0 & 0 & 0 & 0 & 0 & 0 & 0 & 0 & \frac{p_s}{N_{s,TWC}} \\
TC & 0 & \frac{p_b}{N_{b,TWC}} & 0 & 0 & 0 & 0 & 0 & 0 & 0 & 0 & 0 & 0 & 0 & 0 & 0 & 0 & 0 & 0 & \frac{p_s}{N_{s,TWC}} \\
WC & \frac{p_b}{N_{b,TWC}} & 0 & 0 & 0 & 0 & 0 & 0 & 0 & 0 & 0 & 0 & 0 & 0 & 0 & 0 & 0 & 0 & 0 & \frac{p_s}{N_{s,TWC}} \\
Twc & 0 & 0 & 0 & 0 & 0 & 0 & 0 & 0 & 0 & 0 & 0 & 0 & 0 & 0 & 0 & 0 & 0 & \frac{p_b}{N_{b,TWC}} & \frac{p_s}{N_{s,TWC}} \\
tWc & 0 & 0 & 0 & 0 & 0 & 0 & 0 & 0 & 0 & 0 & 0 & 0 & 0 & 0 & 0 & 0 & \frac{p_b}{N_{b,TWC}} & 0 & \frac{p_s}{N_{s,TWC}} \\
twC & 0 & 0 & 0 & 0 & 0 & 0 & 0 & 0 & 0 & 0 & 0 & 0 & 0 & 0 & 0 & \frac{p_b}{N_{b,TWC}} & 0 & 0 & \frac{p_s}{N_{s,TWC}} \\
TWc & 0 & 0 & 0 & 0 & 0 & 0 & 0 & 0 & 0 & 0 & 0 & 0 & 0 & 0 & \frac{p_b}{N_{b,TWC}} & 0 & 0 & 0 & \frac{p_s}{N_{s,TWC}} \\
TwC & 0 & 0 & 0 & 0 & 0 & 0 & 0 & 0 & 0 & 0 & 0 & 0 & 0 & \frac{p_b}{N_{b,TWC}} & 0 & 0 & 0 & 0 & \frac{p_s}{N_{s,TWC}} \\
tWC & 0 & 0 & 0 & 0 & 0 & 0 & 0 & 0 & 0 & 0 & 0 & 0 & \frac{p_b}{N_{b,TWC}} & 0 & 0 & 0 & 0 & 0 & \frac{p_s}{N_{s,TWC}} \\
TWC & \frac{p_s}{N_{s,TWC}} & \frac{p_s}{N_{s,TWC}} & \frac{p_s}{N_{s,TWC}} & 0 & 0 & 0 & 0 & 0 & 0 & \frac{p_s}{N_{s,TWC}} & \frac{p_s}{N_{s,TWC}} & \frac{p_s}{N_{s,TWC}} & \frac{p_s}{N_{s,TWC}} & \frac{p_s}{N_{s,TWC}} & \frac{p_s}{N_{s,TWC}} & \frac{p_s}{N_{s,TWC}} & \frac{p_s}{N_{s,TWC}} & \frac{p_s}{N_{s,TWC}} & \frac{p_e}{N_{e,TWC}}
\end{array}\right)
$$

### Supplement 2A: *Viburnum* Included Affinity Data

**Conservative Included Enabled Affinity Scoring** Viburnum species well outside of the normal climate range, judged by Köppen-Geiger climate zones. We scored warm temperate species as having an enabled affinity for cold temperate biomes if they are found to be free-living outdoors within an arboretum or Botanical garden within a continental climate zone. We chose this scoring as continental climate zones must experience average temperatures below freezing in the coldest month, amply sufficient to be deemed as cold temperate. For some of these species we confirmed the validity of the BGCI records through correspondence with the arboretum/botanical garden in question. Other species we confirmed by identifying the species as present on the institutions collection page. Note we exclude variety-specific and cultivar-specific records in species from this conservative scoring.

**Conservative included cold temperate enabled affinity.** 4 species are included here that have a warm temperate established affinity and cold temperate enabled affinity, from 4 major clades. ! species is included here from with a tropical established affinity and warm temperate included affinity (*V. punctatum*)

*V. punctatum* (*Punctata*) Bot. Gard. of South Australia (Adelaide, Australia) (Confirmed through online collection database, Accession numbers W920631*A/W920631*B and locations M30.3/M30.4); Kunming Bot. Gard, United States National Arb. (Washington D.C., USA); Xishuanbanna Tropical Bot. Gard. (Yunnan, China)

*V. obovatum* (Lentago) Chicago Bot. Gard. (Chicago, USA) (Confirmed per. comm. but shrubbier growth than in the wild); Missouri Bot. Gard. (St. Louis, MO, USA)

*V. hanceanum* (*Lutscentia*) The New York Bot.Gard. (NY, NY, USA)(Confirmed through online collections site, Accession ID: 685/56*A, B and D located in garden with plant finder); The Dawes Arb. (Dawes, OH, USA)

*V. henryi* (Solenotinus) Chicago Bot. Gard. (Chicago, IL, USA) (Confirmed per. comm.); The Morris Arb. (Philadelphia, PA, USA)

*V. davidii* (Tinus)Bot. Gard. of Tartu University (Tartu, Estonia); The New York Bot.Gard. (NY, NY, USA)(confirmed through online collections site, Accession ID: 73/2004*A, 773/2004*B, 773/2004*C); Conservatoire et Jardin botaniques de la Ville de Genave (Geneva, Switzerland); Oklahoma City Zoo and Bot. Gard. (Oklahoma City, OK, USA)

**Bold Included Enabled Affinity Scoring**

Viburnum species in cultivation outside of their normal climate (judging by temperature) – this takes the records from gardens/arboreta more or less at face value – We looked for cases that seemed wrong – e.g., a cold-loving species in a garden climate with little or no freezing; warm-loving species in a garden climate with prolonged freezing in the winter.

UC Berkekely seems like a special case – little freezing, but mild summers. We have omitted species that are only out of place in UC Berkeley (no other supporting records).

Note that these scores give us multiple reservations for several reasons: (1) the species IDs could be wrong (not infrequent); (2) the plant might be in a greenhouse or otherwise protected; (3) plants might have grown there for a short time but then died (and the records are not up to date).

**Bold included cold temperate enabled affinity.** 11 species are included here that have a warm temperate established affinity and cold temperate enabled affinity, from 7 major clades

*V. obovatum* (Lentago) Chicago Bot. Gard. (Chicago, USA) (Confirmed per. comm. but shrubbier growth than in the wild); Missouri Bot. Gard. (St. Louis, MO, USA)

*V. hanceanum* (*Lutscentia*) The New York Bot.Gard. (NY, NY, USA)(Confirmed through online collections site, Accession ID: 685/56*A, B and D located in garden with plant finder); The Dawes Arb. (Dawes, OH, USA)

*V. henryi* (Solenotinus) Chicago Bot. Gard. (Chicago, IL, USA) (Confirmed per. comm.); The Morris Arb. (Philadelphia, PA, USA)

*V. odoratissimum* (Solenotinus) The Linnaean Gardens of Uppsala (Uppsala, Sweden); Missouri Bot. Gard. (St. Louis, MO, USA)

*V. suspensum* (Solenotinus) Bot. Gard. of Tartu University (Tartu, Estonia)

*V. davidii* (Tinus)Bot. Gard. of Tartu University (Tartu, Estonia); The New York Bot.Gard. (NY, NY, USA)(confirmed through online collections site, Accession ID: 73/2004*A, 773/2004*B, 773/2004*C); Conservatoire et Jardin botaniques de la Ville de Genave (Geneva, Switzerland); Oklahoma City Zoo and Bot. Gard. (Oklahoma City, OK, USA)

*V. tinus* (Tinus) Oklahoma City Zoo and Bot. Gard. (Oklahoma City, OK, USA); The Linnaean Gardens of Uppsala (Uppsala, Sweden); The New York Bot.Gard. (NY, NY, USA)

*V. corylifolium* (*Sucotinus*) Kunming Bot. Gard. (Kumming, China); University of California Bot. Gard. at Berkeley (Berkley, CA, USA); Pukekura Park (Plymouth New Zealand)

*V. foetidum var. rectangulatum* (Succotinus) The Morris Arb. (Philadelphia, PA, USA)

*V. japonicum* (Succotinus) Missouri Bot. Gard. (St. Louis, MO, USA)

*V. luzonicum* (Succotinus) Missouri Bot. Gard. (St. Louis, MO, USA)

**Bold included warm temperate enabled affinity.** 13 species are included here that have a cold temperate established affinity (or tropical for *V. punctatum*) and warm temperate enabled affinity, from 7 major clades

*V. punctatum* (*Punctata*) Bot. Gard. of South Australia (Adelaide, Australia) (Confirmed through online collection database, Accession numbers W920631*A/W920631*B and locations M30.3/M30.4); Kunming Bot. Gard, United States National Arb. (Washington D.C., USA); Xishuanbanna Tropical Bot. Gard. (Yunnan, China)

*V. lentago* (Lentago) Sarah P. Duke Gardens (Durham, NC, USA); Jerusalem Bot. Gard. (Jerusalem, Israel); University of California Bot. Gard. at Berkeley (Berkley, CA, USA)

*V. glomeratum* (*Euviburnum*) Xishuanbanna Tropical Bot. Gard. (Yunnan, China)

*V. rhytidophyllum* (*Euviburnum*) Auckland Bot. Gard. (Auckland, New Zealand), Xishuanbanna Tropical Bot. Gard. (Yunnan, China)

*V. schensianum* (*Euviburnum*) Jerusalem Bot. Gard. (Jerusalem, Israel)

*V. utile* (*Euviburnum*) Kunming Bot. Gard. (Kumming, China

*V. plicatum* (*Lutscentia*) Auckland Bot. Gard. (Auckland, New Zealand); Pakekura Park (Plymouth New Zealand); Kunming Bot. Gard. (Kumming, China)

*V. opulus* (Opulus) Jerusalem Bot. Gard. (Jerusalem, Israel); University of California Bot. Gard. at Berkeley (Berkley, CA, USA); Jardín Botánico Carlos Thays (Buenos Aires, Argentina); Auckland Bot. Gard. (Auckland, New Zealand)

*V. sargentii* (Opulus) Auckland Bot. Gard. (Auckland, New Zealand); Atlanta Bot. Gard. (Atlanta, USA); University of California Bot. Gard. at Berkeley (Berkley, CA, USA)

*V. trilobum* (Opulus) Auckland Bot. Gard. (Auckland, New Zealand)

*V. betulifolium* (*Sucotinus*) Kunming Bot. Gard. (Kumming, China); University of California Bot. Gard. at Berkeley (Berkley, CA, USA); Pukekura Park (Plymouth New Zealand)

*V. foetidum var. rectangulatum* (Succotinus) The Morris Arb. (Philadelphia, PA, USA)

*V. setigerum* (*Sucotinus*) Auckland Bot. Gard. (Auckland, New Zealand)

#### Supplement 2B: *Viburnum* Excluded Affinity Data

**Explanation/Metadata for Table S**

**Column A.** Species names correspond to those in Landis et al. (2021). Donoghue et al. (2022) made a number of taxonomic and nomenclatural changes in their analysis of the neotropical *Oreinotinus* clade; for details see Donoghue et al., 2022, Supplementary Tables 1 and

2 for details). All *Oreinotinus* species are scored as occupying cloud forests and the minor taxonomic changes made by Donoghue et al. (2022), and the small differences in tree topology that they obtained for this clade, do not influence the results reported here.

**Column B.** Major clade names from Clement et al. (2014) correspond to those in Landis et al. (2021). These are used in some analyses here to extrapolate from species that have been studied in detail.

**Column C.** Biome assignments (4 states) were obtained from Landis et al. (2021; also see Edwards et al., 2017), but updated as follows based on newer information: (1) *V. lantana* and *V. maculatum* are now 1,3 based on its geographic/climatic range as documented in Kollmann and Grubb (2002) and Park and Donoghue (2021); (2) *V. rufidulum* is now scored 1,3 based on its geographic/climatic range as documented in Spriggs et al. (2019a)

O=tropical rainforest; 1=evergreen warm temperate (lucidophyllous) forest; 2=cloud forest; 3=cool temperate forest; ?=uncertain. Note that some species are scored with two states (i.e., 0,1 or 1,3).

In our dataset there are 163 species, which is a "complete" sample with the exception of several recent taxonomic changes within in *Oreinotinus* (Donoghue et al., 2022). Of these, 12 species are scored 0, 47 are scored 1, 36 are scored 2, 56 are scored 3, 2 are scored 0,1 (1 uncertain, ?), and 10 are scored 1,3 (6 uncertain, ?).

**Column D.** Biome assignments (3 states) with evergreen warm temperate (lucidophyllous) forest and cloud forests lumped together as extratropical warm forests without prolonged freezing.

O=tropical rainforest; 1= extratropical warm forest; 2=cool temperate forest. Uncertainty=(?)

In our dataset there are 83 species scored 1 (47 scored in column C as 1 plus 36 scored in column C as 2).

**Column E.** Seed dormancy/germination type. *Viburnum* species differ with respect to germination requirements (Baskin and Baskin, 1998; Baskin et al., 2006; Baskin et al., 2009). Each fruit contains a single seed that is enclosed inside of a hardened endocarp formed by the inner wall of the ovary. Each seed contains one tiny and underdeveloped embryo embedded in copious endosperm at its apical end. To germinate, the triggered embryo has to grow a relatively long distance through the endosperm before the radicle (embryonic root) can emerge. Then, typically, it must overcome a physiological barrier before the epicotyl (embryonic shoot) can emerge. This pattern of development is known as morpho-physiological dormancy (MPD), and in *Viburnum* the phases of growth are mediated by the temperatures experienced. To date, germination has been studied to varying degrees in 27 *Viburnum* species, and it has been found that these have one of the four different levels of MPD described below. Four other species of the *Oreinotinus* clade have been studied by MJD and colleagues at Yale (bringing the total to 31 species), but these results have not been published.

We note that MPD has been documented in several species of *Sambucus* (Hidayati et al., 2000), as well as in *Adoxa* (Baskin et al., 2006). Together these two form the sister clade to *Viburnum*. MPD is very likely to have been ancestral in the Dipsacales judging by its presence in related Campanulids based on the size of the underdeveloped embryo relative to the volume of endosperm (Baskin et al., 2006). Within the Dipsacales MPD or morphological dormancy (MD) has also been reported in the early-diverging lineages of the Caprifoliaceae s.l. (e.g., in *Weigela*, *Diervilla*, *Heptacodium*, *Lonicera*, *Triosteum*, *Symphoricarpos*, *Linnaea*, and *Kolkwitzia*). MPD/MD appears to have been lost within the Dipsacales in the large herbaceous Valerina clade in which the embryo is well developed and the endosperm is reduced or absent (Baskin et al., 2006).

1. Deep simple epicotyl MPD

This requires a prolonged warm period for embryo growth through the endosperm and radicle emergence and a prolonged cold period before epicotyl emergence. This germination behavior has often been described as "double dormancy" and it is associated with a very elongate process in nature in which seeds are distributed in the fall of a first year and remain ungerminated through the winter, then the radicle emerges during the prolonged warm period of the following growing season and the seed overwinters in this half-germinated condition, and finally the epicotyl emerges after the prolonged cold period during the following winter. In other words, in nature, the completion of germination can require 1.5 years, spanning two winters. This can be sped up under controlled conditions in the greenhouse, or sometimes given unusual climatic conditions in nature, but in any case germination often still require 7-9 months.

Deep simple epicotyl MPD has been confirmed in the following 17 species, representing 7 different major clades (clade name in parentheses), all native to cold climates and generally subjected to prolonged periods of freezing in the wild:

*V. furcatum* (Pseudotinus) (Phartyal et al., 2014)

*V. lantanoides* (Pseudotinus) (Gill and Pogge, 1974)

*V. cassinoides* (Lentago) (Gill and Pogge, 1974)

*V. lentago* (Lentago) (Giersbach,1937; Baskin et al., 2008)

*V. prunifolium* (Lentago) (Giersbach,1937; Baskin et al., 2008)

*V. rufidulum* (Lentago) (Giersbach,1937; Baskin et al., 2008) ***Limited freezing in the southeastern USA

*V. rafinesquianum* (*Mollotinus*) (Gill and Pogge, 1974)

*V. dentatum* (Dentata) (Giersbach,1937; Baskin et al., 2008) ***Exact ID uncertain

*V. recognitum* (?? as *V. pubescens*) (Dentata) (Giersbach,1937)

*V. edule* (Opulus) (Baah-Acheamfour and Sobze 2022)

*V. opulus* (Opulus) (Giersbach,1937; Walck et al., 2012)

*V. sargentii* (Opulus) (Walck et al., 2012; Zhang et al., 2019)

*V. trilobum* (Opulus) (Walck et al., 2012; Fedec and Knowles, 1973; Knowles and Zalik, 1958)

*V. acerifolium* (Lobatas) (Giersbach, 1937; Hidayati et al., 2005)

*V. dilatatum* (Succotinus) (Giersbach,1937; Baskin et al., 2008)

*V. betulifolium* (Succotinus) (Chien et al. 2011)

*V. parvifolium* (Succotinus) (Chien et al. 2011)

*V. mullaha* (Succotinus) (Pharswan et al., 2019)

2 Non-deep simple epicotyl MPD

This requires a short warm period for root emergence and an additional warm period for shoot emergence. The following 7 species have been documented in this category, all native to warm climates without prolonged freezing:

*V. nudum* (Lentago) (Giersbach, 1937; Baskin et al., 2008)

*V. odoratissimum* (Solenotinus) (Baskin et al., 2008)

*V. tinus* (Tinus) (Karlsson et al., 2005)

*V. treleasei* (Tinus) (Moura and Silva, 2010)

*V. propinquum* (Tinus) (Chien and Chen, 2012)

*V. scabrellum* (Dentata) (Giersbach, 1937; Baskin et al., 2008)

*V. formosanum/luzonicum* (Succotinus) (Baskin et al., 2009): there is taxonomic confusion in this case; we currently recognize two species and have scored them both as having this form of dormancy.

Based on unpublished germination studies at Yale, four neotropical cloud forest species appear to be in this category. Specifically, we found that seeds of *V. lautum*, V. *jucundum*, and *V. hartwegii* from Chiapas, Mexico could be germinated without a cold period. Seedling emergence took 7-10 months, and was unreliable. More synchronous germination was achieved when we interjected a cold stratification period of up to four months, but a cold period was not necessary for germination. The same results were obtained when we attempted to germinate seeds generated from a manual cross between *V. caudatum* (maternal parent) from eastern Mexico and *V. recognitum* (paternal parent) from northeastern North America.

3. Deep complex MPD

In this case germination can proceed at cool temperatures, with no warm period necessary, though there is still a delay between embryo growth through radicle emergence and growth of the epicotyl. Only 1 species has been confirmed to date, from a cool mountainous region in central Taiwan: *V. plicatum var. formosanum* (Lutescentia) (Chen et al., 2021). We note that this is a case in which there is taxonomic confusion, and the studied plants may instead correspond to *V. hanceanum*.

4. Non-deep simple MPD

Here there is no epicotyl dormancy, and therefore no delay between root and shoot emergence. That is, it appears that there is no physiological barrier and the embryo can grow continuously until it fully emerges. Only 1 species has been confirmed to date: *V. lantana* (Euviburnum) (Santiago et al., 2014; also see Bezděčková et al., 2009; Moghimifam Haghighi, 2022)

**Column F.** Reference to the study/studies documenting seed dormancy/germination type.

Baah-Acheamfour, M., Sobze, J. M. 2022. Exposing *Viburnum edule* seeds to a sequence of temperatures affects the germination characteristics. Seed Science and Technology *50*: 235-239.

Baskin, C. C., Chen, S. Y., Chien, C. T., Baskin, J. M. 2009. Overview of seed dormancy in *Viburnum* (Caprifoliaceae). Propagation of Ornamental Plants 9: 115-121.

Baskin C C., Chien C.-T., Chen S.-Y., Baskin J. M. 2008. Germination of *Viburnum odoratissimum* seeds: a new level of morpho-physiological dormancy. Seed Science Research 18: 179-184.

Baskin J. M., Hidayati S. N., Baskin C. C., Walck J. L., Huang Z. Y., Chien C. T. 2006. Evolutionary considerations of the presence of both morphophysiological and physiological seed dormancy in the highly advanced euasterids II order Dipsacales Seed Science Research 16: 233-242.

Bezděčková, L., Řezníčková, J., & Procházková, Z. 2009. Germination of stratified seeds and emergence of non-stratified seeds and fruits of *Viburnum lantana, Euonymus europaeus* and *Staphylea pinnata.* Zprávy Lesnickeho Výzkumu 54: 275-285.

Chen, S. Y., Liu, C. P., Baskin, C. C., Chien, C. T. 2021. Deep complex morphophysiological dormancy in seeds of *Viburnum plicatum* var. *formosanum* (Adoxaceae) from subtropical mountains. Seed Science Research 31: 236-242.

Chien, C. T., Chen, S. Y. 2012. Seed germination and dormancy in the woody plant *Viburnum propinquum* (Caprifoliaceae) from a temperate mountain in Taiwan. In II International Symposium on Woody Ornamentals of the Temperate Zone 990: 451-456.

Chien, C. T., Chen, S. Y., Tsai, C. C., Baskin, J. M., Baskin, C. C., Kuo-Huang, L. L. 2011. Deep simple epicotyl morphophysiological dormancy in seeds of two *Viburnum* species, with special reference to shoot growth and development inside the seed. Annals of Botany 108: 13-22.

Fedec P., Knowles R. H. 1973. After ripening and germination of seeds of American highbush cranberry (*Viburnum trilobum*). Canadian Journal of Botany 51: 1761-1764.

Giersbach J. 1937. Germination and seedling production of species of *Viburnum.* Contributions from Boyce Thompson Institute 9: 79-90.

Gill J. D., Pogge F. L. 1974. *Viburnum* L. *In:* C.S. Schopmeyer (Tech. coord.). Seeds of woody plants in the United States. Agriculture Handbook No. 450. USDA Forest Service: 844-850.

Hidayati S. N., Baskin J. M., Baskin C. C. 2000. Morphophysiological dormancy in seeds of two North American and one Eurasian species of *Sambucus* (Caprifoliaceae) with underdeveloped spatulate embryos. American Journal of Botany 87: 1669-1678.

Hidayati S. N., Baskin J. M., Baskin C. C. 2005. Epicotyl dormancy in *Viburnum acerifolium* (Caprifoliaceae). American Naturalist 153: 232-244.

Karlsson l. M., Hidayati S. N., Walck J. l., Milberg P. 2005. Complex combination of seed dormancy and seedling development determine emergence of *Viburnum tinus* (Caprifoliaceae). Annals of Botany 95: 323-330.

Knowles R. H., Zalik S. 1958. Effects of temperature treatment and of a native inhibitor on seed dormancy and of cotyledon removal on epicotyl growth in *Viburnum trilobum* Marsh. Canadian Journal of Botany 36: 561-566.

Lamichhane, G., Poudel, P., Pandeya, P. R., Adhikari, M., Sharma, G. (2023). *Viburnum* spp. (*Viburnum erubescens* Wall., *Viburnum mullaha* Buch.-Ham. ex D. Don). Pp. 459-465 in Himalayan Fruits and Berries. Academic Press.

Moghimifam, R., Haghighi, A. R. 2022. Evaluation of dormancy breaking treatments for enhanced germination in *Cotinus coggygria*, *Cornus mas* and *Viburnum lantana* seeds. Seed Science and Technology 50: 323-328.

Moura, M., Silva, L. 2010. Seed germination of *Viburnum treleasei* Gand., an Azorean endemic with high ornamental potential. Propagation of Ornamental Plants 10: 129-135.

Pharswan, D. S., Maikhuri, R. K., Bhandari, B. S., Jugran, A. K., Rawat, L. S., Negi, V. S. 2019. Conservation strategies and seed germination of *Viburnum mullaha* (Buch.-Ham ex D. Don (Indian cranberry) growing in western Himalaya, India. Himalayan Ecology *27*: 29.

Phartyal, S. S., Kondo, T., Fuji, A., Hidayati, S. N., Walck, J. L. 2014. A comprehensive view of epicotyl dormancy in *Viburnum furcatum*: combining field studies with laboratory studies using temperature sequences. Seed Science Research 24: 281-292.

Santiago, A., Ferrandis, P., & Herranz, J. M. 2015. Non-deep simple morphophysiological dormancy in seeds of *Viburnum lantana* (Caprifoliaceae), a new dormancy level in the genus *Viburnum*. Seed Science Research 25: 46-56.

Walck, J. L., Karlsson, L. M., Milberg, P., Hidayati, S. N., Kondo, T. 2012. Seed germination and seedling development ecology in world-wide populations of a circumboreal Tertiary relict. AoB PLANTS, pls007.

Zhang, Q., Li, Y., Sun, X., Xing, S., Cong, W., & Liu, X. 2019. Study on dormancy mechanism and breaking dormancy method of *Viburnum sargentii* seeds. American Journal of Plant Sciences 10: 65-78.

**Column G.** Biome non-affinity (4 states) based on seed germination type and expert knowledge.

The key observation for present purposes is that some *Viburnum* species – those listed above under 1 with deep simple epicotyl MPD – require a prolonged cold period before germination can be completed. In nature this would correspond with the northern winter during which the seeds in the soil would experience prolonged periods of freezing. Under greenhouse conditions continued growth of the epicotyl can be induced with low temperatures that are above freezing – a stratification temperature of 5C is typical in the experiments that have been conducted. Though it appears not to require a prolonged warm period for germination, *V. plicatum* in group 3, with deep complex MPD, falls in this category, as a prolonged cold period appears necessary.

At the other extreme are those listed above under 4 with non-deep simple epicotyl MPD. These do not require cold for germination and, judging by their consistent association with warm biomes, would be unlikely to survive prolonged cold. Though it can withstand cold temperatures, *V. lantana* in group 3, with non-deep simple MPD, falls for in this category from the standpoint of not requiring cold.

From the above we infer that species with the need for prolonged cold for germination, which currently occupy cool temperate forests (biome 3), would be prohibited from successfully occupying climates where there is not such a cold period – this would include tropical forests without freezing (biome 0), warm temperate forests with no or very little cold (biome 1), and cloud forests that experience some nights with freezing, but not a prolonged cold period (biome

2). Thus, is appears to be an evolutionary ratchet whereby once a species has double dormancy, and requires prolonged cold for germination, it would be difficult to return to a climate that lacks sufficient cold, as it would be prevented from ever becoming established in such a climate.

At the other extreme are seeds that do not require cold, and most likely could not withstand prolonged cold. We note that this has not yet been rigorously tested by subjecting populations of seeds with non-deep simple epicotyl MPD to prolonged freezing to assess their survival rate. In the meantime, however, for present purposes we will infer that their germination in a cold climate would be hindered and that they would therefore be unlike to become established in a biome with prolonged freezing during the winter (biome 3). On the other hand we assume that these species could freely move between biomes 0 and 1, and probably also biome 2. The latter assertion is also untested, in the sense that we do not know how sensitive seeds in this category would be to harsh freezes of one or a few days.

*In summary, based on the reasoning outlined above we have scored species in the first category (germination types 1 and 3) as having a non-affinity for biomes 0, 1, and 2, and we have scored species in the second category (with germination types 2 and 4) as having a non-affinity for biome 3.* We stress however that unlikely does not mean impossible, and it is evident that some unlikely transitions surely must have occurred to yield the observed phylogenetic distribution of the biome states. Also, there are some cases of variation within species (e.g., *V. lantana* and *V. maculatum* in Europe occupy both warm temperate and cool temperate forests; likewise *V. rufidulum* in eastern North America). There are also cases of close relatives occupying both biomes (e.g., *V. nudum* in warm forests and *V. cassinoides* in cool forests in eastern North America; *V. taiwanianum* in warm forests in Taiwan and *V. urceolatum* in cool forests in Japan; the widespread Asian species *V. odoratissimum* in warm forests and its close relative *V. sieboldii* in cool forests in Japan). These observations highlight evolutionary lability in germination and/or biome occupancy.

**Column H.** Biome non-affinity (3 states) based on seed germination type and expert knowledge.

Biome assignments with evergreen warm temperate (lucidophyllous) forest and cloud forests lumped together as extratropical warm forests without prolonged freezing.

O=tropical rainforest; 1= extratropical warm forest; 2=cool temperate forest. Uncertainty=(?)

In our dataset there are 83 species scored 1 (47 scored in column C as 1 plus 36 scored in column C as 2).

**Column I.** Unlikely biome(s) (4 states) with a conservative phylogenetic extension based on known germination and expert knowledge.

In cases where germination has been studied in at least one species of a major clade, and where there is only one germination type known in the clade, and where all species in the clade occupy a similar climatic range, we extended that scoring of the known species to the unsampled species in that clade. This applied to the following major clades: *Pseudotinus*, *Opulus*, and *Lobata*, and with germination type 1, and *Tinus* and *Oreinotinus* with germination type 2.

In cases where there were any differences among the studied species in either the germination type or the climatic range, where possible we extended scorings based on phylogenetic relatedness and expert knowledge of the species. In the Lentago clade, we scored the unstudied *V. obovatum* of the southeastern United States, and *V. elatum* of Mexico, the same as *V. nudum* (germination type 2) based on Spriggs et al. (2019a, b). In Euviburnum we extended the scoring of *V. lantana* only to its sister species, *V. maculatum.* In this conservative scoring, we have not scored any other species in this clade. In Lutescentia, *V. plicatum* is the only species that occupies cool temperate forests, so we have extended its scoring only to its sister species, *V. hanceanum*, which occupies somewhat warmer forests at lower latitudes (Park and Donoghue, 2021). In Solenotinus, we extended the scoring of *V. odoratissium* only to its very similar sister species, *V. awabuki.* Climatic ranges in in Succotinus are too variable to extend the scorings based on the studied species. In Mollotinus we extended the scoring of *V. rafinesquianum* to its close relatives *V. molle* and V. *bracteatum*, but not to the climatically distinct *V. australe* and *V. ellipticum.*

**Column J.** Unlikely biome(s) (3 states) with a conservative phylogenetic extension based on known germination and expert knowledge.

See Column I for details. Biome assignments with evergreen warm temperate (lucidophyllous) forest and cloud forests lumped together as extratropical warm forests without prolonged freezing. O=tropical rainforest; 1= extratropical warm forest; 2=cool temperate forest. Uncertainty=(?)

**Column K.** Unlikely biome(s) (4 states) with a bold phylogenetic extension based on known germination type and expert knowledge.

Under this scoring, values are filled in for as many species as possible, based in most cases on the experience of MJD with these species in the field; those that are too poorly known in the field are left unscored here. For species scored in Column I, the scores here are the same. Additional scores were made as follows: (1) all tropical species (scored 0 in column C and D) are inferred to be unlikely to occupy cool temperate forests (biome 3). (2) Except as noted below, species scored as occupying cool temperate forests (biome 3) in Column C are inferred to be unlikely to occupy tropical forests. (3) Based on Park and Donoghue (2021), species in the Euviburnum clade that occupy the coolest habitats (*V. mongolicum and V. burejaeticum, and* the clade containing *V. schensianum, V. utile, V. congestum, V. carlesii, V. bitchiuense*) are inferred to be unlikely to occupy biomes 0, 1, or 2; the remaining species are scored the same as the studied species, *V. lantana.* (4) Species of Lutescentia other than *V. plicatum* and *V. hanceanum* occupy tropical or lucidophyllous forests in Asia, and these we infer to be unlikely to occupy cool temperate forests (biome 3). (5) The several species of Solenotinus that occupy notably cooler climates (*V. farreri, V. grandiflorum, V. foetens*) are inferred to be unlikely to occupy biomes 0, 1, and 2; and the cool temperate *V. sieboldii* is inferred to be unlikely to occupy tropical forests (biome 0). The remaining species, which are known through personal experience to occupy lucidophyllus forests (and are scored 1 in column C) are scored as unlikely to occupy cool temperate forests (biome 3): these are *V. taitoense, V. suspensum, V. henryi, and V. brachybotryum.* The remaining species in this clade are poorly understood and are left unscored. (6) The two remaining species of the Tinus clade are too poorly understood, and are left unscored. (7) All species of Coriacea and Sambucina occupy topical or lucidophyllus forests in southeastern Asia (Clement and Donoghue, 2011) and these we infer to be unlikely to occupy cool temperate forests (biome 3). (8) The climate ranges of Succotinus species are variable and in some cases too poorly understood to be scored here with confidence. Those species known to occupy cool temperate forests are here scored as being unlikely to occupy tropical forests (biome

0): *V. melanocarpum, V. lobophyllum, V. hupehense, V. wrightii , V. ichangense, V. setigerum, V. phlebotrichum, V. erosum,* and *V. brachyandrum.* Those known to occupy warmer temperate forests are scored here as being unlikely to occupy cool temperate forests (biome 3): *V. foetidum, V. integrifolium, V. japonicum, V. hainanense, V. adenophorum, V. tashiroi,* and *V. sempervirens.* (9) The *Mollotinus* species *V. australe* (northeastern Mexico and western Texas) and *V. ellipticum* (northwestern North America) occupy warmer forests and are here scored as unlikely to occupy tropical forests. (10) All species of *Oreinotinus* occupy cloud forests in the neotropics, and are scored here as unlikely to occupy cool temperate forests (biome 3).

**Column L.** Unlikely biome(s) (3 states) with a bold phylogenetic extension based on known germination type and expert knowledge.

See Column K for details. Biome assignments with evergreen warm temperate (lucidophyllous) forest and cloud forests lumped together as extratropical warm forests without prolonged freezing. O=tropical rainforest; 1= extratropical warm forest; 2=cool temperate forest. Uncertainty=(?)

**Column M.** Leafing Habit (2 states). E=evergreen, or leaf exchanging; D=deciduous, or semi-deciduous. Based on Edwards et al. (2017), Yang and Malecot (2011), and expert knowledge.

In our dataset there 88 species scored as evergreen (E) and 72 scored as deciduous (D); we are uncertain about the remaining 3 species.

Evergreen viburnums are unlikely to survive prolonged freezing. The evidence for this is two-fold. First, in nature, evergreen species only occupy warmer climates where freezes do not occur or are rare and not prolonged. There are five exceptions, all of them Chinese species of Euviburnum. *Viburnum rhytidophyllum, V. buddleifolium*, *V. chinshanense*, *V. utile*, and *V. congestum* can tolerate freezing temperatures and are evergreen to semievergreen in the wild and in cultivation. Some of these are observed to experience significant leaf damage and stem die-back during cold winter and behave more like leaf-exchangers (adding new leaves at the same time that the senescing leaves of the past season are dropping). In cultivation some of these can be grown, for example, in the northern New England states (e.g., *V. rhytidophyllum*), but others (e.g., *V. utile*) are not found in cultivation much north of Washington DC (roughly beyond USDA hardiness zone 6b).

Second, there is direct evidence that evergreen species cannot tolerate cold. For example, a cultivated plant of the Bornean species *V. clemensiae* did not survive a cold-snap in a Yale Marsh Garden greenhouse. Neotropical cloud forest species can grow well in temperate greenhouses that experience occasional cold temperatures, but our attempts to grow these outdoors in Connecticut have failed. Specifically, *V. lautum* and *V. jucundum* have been unable to survive prolonged winter freezing. In contrast, these same species have been grown outdoors for multiple years in the San Francisoco area (e.g., in Golden Gate Park and in the UC Berkeley Botanical Garden). Plants of *V. tinus* and *V. davidii* have survived several seasons in places such as the New York Botanical Garden, but only in protected areas, and then not for long before they succumb to low winter temperatures. Other members of the Tinus clade have been cultivated successfully in the vicinity Washington DC (e.g., *V. propinquum*). The same is true of several evergreen species of the Solenotinus clade; we note, for example, that *V. odoratissimum* is thriving in the US National Arboretum. Several other evergreen species of Tinus (e.g., *V. cinnamomifolium*) and Solenotinus (e.g., *V. henryi*) have also been successfully cultivated in the coastal northwest of North America, such as in the University of Washington Arboretum in Seattle. Other evergreen species are known in cultivation only in warmer more southern regions. For example, *V. suspensum* (Solenotinus), which is narrowly endemic to the Ryuku Islands, has been widely cultivated across the southern United States and in Mexico (and elsewhere), but does not survive in the eastern United States north of the Carolinas. Surprisingly, it can tolerate much drier conditions than other viburnums and even thrives under desert conditions with proper irrigation (e.g., in Southern Arizona).

How do cold-adapted deciduous viburnums fare in warm climates? Assuming that they can germinate without prolonged cold (see above), deciduous species can survive continuous warm temperatures and become established. We note, first, that a few deciduous species grow in nature in warm/tropical climates (e.g., *V. luzonicum* in the Philippines), so this can be successful. Second, we know this from growing temperate species under warm greenhouse conditions for multiple seasons and through successful flowering and fruiting. At the Yale Marsh Garden greenhouses we have grown plants of multiple temperate species, especially from the Dentata, Lentago, and Succotinus clades. We have observed, however, that they grow far less well after several years under such conditions, and that their growth is promoted by shifting them into cooler greenhouse conditions during the winter where they form well-developed winter buds and become dormant for a period of time. Overall, we infer that viburnums adapted to cold climates can be grown in continuously warm climates, but that long-term survival in such climates would likely entail evolutionary adjustments in phenology.

As documented especially by Edwards et al. (2017), there have been multiple (~15) evolutionary shifts from the evergreen to the deciduous habit, and some (but many fewer) shifts in the other direction. These shifts have been tightly correlated with the evolution of leaf teeth (Schmerler et al., 2012), with seasonal heteroblasty (Spriggs et al., 2018), and with shifts into colder climates with prolonged freezes during the winter (Edwards et al., 2017). These observations support the view that these traits are highly evolutionarily labile and might be expected to evolve *in situ* in response to gradual climate change. Regarding causal factors, Edwards et al. (2017) showed a very tight correlation in *Viburnum* species in Taiwan and Japan between the deciduous habit and mean monthly minimum temperatures that fall below freezing during the winter in December, January, and February. Evergreen species in these areas are not found under such conditions.

Collectively, these observations lead us to the following conclusions. Evergreen species would have difficulty becoming established in climates with prolonged winter freezing. Here we have scored them as having an affinity for tropical (0), warm temperate (1), and cloud forests (2), but a non-affinity for cold temperate forests (3). deciduous species would have an easier time becoming established in warm forests, and would stand a better chance of adapting their phenology to these climates. In one scoring, we have not specified non-affinities for deciduous species. In another scoring we recognize a non-affinity of deciduous species only for the most tropical forests (0); this supposes that they would fare better in climates with a somewhat colder winter (warm temperate forests, 1) or with diurnal fluctuation dipping below freezing (cloud forests, 2).

**Column N.** Unlikely biome(s) (4 states) based on leafing habit (conservative scoring)

O=tropical rainforest; 1=evergreen warm temperate (lucidophyllous) forest; 2=cloud forest; 3=cool temperate forest.

We infer based on the evidence above that evergreen species would have difficulty becoming established in climates with prolonged winter freezing. Here we have scored them as having an affinity for tropical (0), warm temperate (1), and cloud forests (2), but a non-affinity for cold temperate forests (3).

deciduous species would have an easier time becoming established in warm forests, and would stand a better chance of adapting their phenology to these climates. Under this conservative scoring we score no non-affinities for deciduous species.

**Column O.** Unlikely biome(s) (4 states) based on leafing habit (bold scoring)

O=tropical rainforest; 1=evergreen warm temperate (lucidophyllous) forest; 2=cloud forest; 3=cool temperate forest.

As in Column N for evergreen species. For deciduous species we here recognize a non-affinity of deciduous species only for tropical forests (0); this supposes that they would fare better in climates with a somewhat colder winters (warm temperate forests, 1) or with diurnal fluctuation dipping below freezing (cloud forests, 2).

**Column P.** Unlikely biome(s) (3 states) based on leafing habit (conservative scoring)

O=tropical rainforest; 1= extratropical warm forest; 2=cool temperate forest.

Evergreen species would have difficulty becoming established in climates with prolonged winter freezing. Here we have scored them as having an affinity for tropical (0) and warm temperate (1), but a non-affinity for cold temperate forests (2).

deciduous species would have an easier time becoming established in warm forests, and would stand a better chance of adapting their phenology to these climates. Under this conservative scoring we score no non-affinities for deciduous species.

**Column Q.** Unlikely biome(s) (3 states) based on leafing habit (bold scoring)

O=tropical rainforest; 1= extratropical warm forest; 2=cool temperate forest.

As in Column P for evergreen species. For deciduous species we here recognize a non-affinity of deciduous species only for the most tropical forests (0); this supposes that they would fare better in climates with a somewhat colder winters, or with diurnal fluctuation dipping below freezing (1).

**References for leafing:**

Include Egolf papers and Dirr book.

Marjorie Weber paper too

**Column R.** Cultivated somewhere outdoors

Viburnum species in cultivation. In total, 87 of the 163 Viburnum species recognized here (i.e., more than half) have been or are currently in cultivation outdoors somewhere in the world. This includes representatives of all 16 clades listed in Column B except for the more (sub)tropical Lutescentia, Coriaceae, and Sambucina clades from Southeast Asia, and the cloud forest species of Orienotinus from the neotropics. Also not in cultivation are the two deeply branching tropical species from Borneo, *V. clemensiae* and *V. amplificatum.* These scores are based primarily on Dirr (2007) but with additions based on personal observations by MJD as follows: *V. urceolatum* has been observed in cultivation in the US National Arboretum, *V. puncatum* in the Kunming Botanical Garden, *V. maculatum* in the Arnold Arboretum, *V. hanceanum* in the US National Arboretum, *V. lutescens* in the Bogor Botanical Garden, *V. sambucinum* in the Bali Botanical Garden, *V. melanocarpum* in the Arnold Arboretum, *V. brachyandrum* in the Tokyo Botanical Garden, *V. lautum* in the UC Berkeley Botanical Garden and the Strybing Arboretum, *V. jucundum* in the Strybing Arboretum, and *V. hartwegii* in the Strybing Arboretum. Additional scores for *V. congestum*, *V. taitoense*, *V. oliganthum*, and *V. parvifolium* are based on on-line searches of commercial nurseries; these sites are deemed reliable by virtue of providing a trust-worthy description and/or photograph. *V. scabrellum* is not specifically listed by Dirr (2007); we assume that this is included in his broad circumscription of *V. dentatum* and we are confident that plants from the southern part of the range of *V. dentatum* s. l. have been cultivated. Two species are scored "?" to signify that it is uncertain whether they are in cultivation. Of these, *V. kansuense* is listed in Dirr (2007), but is scored here as "?" because his photograph does not show this species, and from his description it is unclear what material he observed. *V. orientale* and *V. nervosum* are similarly problematical in Dirr (2007), but are scored here as being in cultivation based on other sources. *Viburnum fordiae* was not listed by Dirr (2007); it may be in cultivation but is scored here as "?". In almost all cases, species that have been brought into cultivation have been grown outside of their native region/range (e.g., many species were introduced into western horticulture from wild collections obtained in China and Japan). However, as far as we are aware the following species are cultivated only within their native region/range: *V. punctatum*, *V. lutescens* and *V. sambucinum* in Southeast Asia, and *V. lantanoides* in northeastern North America.

**Column S.** USDA Hardiness Zones from Dirr (2007)

Here we give the USDA Hardiness Zones indicated by Dirr (2007). USDA Hardiness Zones are based on the average annual extreme minimum temperature during the winter across the United States. Zones 1-6 experience minimum temperature below freezing (0 degrees F; -15 degrees C); zones 7-13 do not experience average minimum temperatures below freezing. All *Viburnum* species in cultivation fall in the range of zones from 2 (-50 to -40 F; *V. edule*) to 11 (+40 to +50; *V. suspensum*, *V. odoratissimum*). The majority of Viburnum species in cultivation span the freezing divide (i.e., they include at least zones 6 and 7). These are species that can tolerate some (variable) degree of prolonged winter freezing.

The following three species are listed in Zones 2 and 6 (i.e., having average minimum temperatures below freezing): *V. lantanoides*, *V. edule*, and *V. recognitum.* Several others are listed as having a range starting from zone 3 and sometimes extending to zone 7 or 8: *V. cassinoides*, *V. lentago*, *V. burejaeticum*, *V. trilobum*, *V. sargentii*, *V. opulus*, *V. acerifolium*, *V. rafinesquianum*, and *V. dentatum.* In Zones 7 and 8 these plants would occasionally (but not consistently) experience winter temperatures below freezing. Collectively, these 12 species are considered the most "cold-loving" of all *Viburnum* species, and it is noteworthy that all of them have deciduous leaves with prominent lobes and/or toothed margins.

In contrast, the following two species are listed only in zone 8 and higher: *V. odoratissimum* and *V. suspensum*. The following 14 species are listed only in zone 7 and higher (not experiencing average minimum temperatures below freezing): *V. awabuki*, *V. henryi*, *V. tinus*, *V. rigidum*, *V. davidii*, *V. propinquum*, *V. calvum*, *V. atrocyaneum*, *V. cylindricum*, *V. foetidum*, *V. mullaha*, *V. japonicum*, *V. luzonicum*, and *V. sempervirens*. Collectively, these 16 species are considered to be the most "warm-loving" (or the most freezing intolerant) *Viburnum* species currently in cultivation. It is noteworthy that all of these are species whose ranges are in subtropical to tropical biomes, and that they all have evergreen leaves with entire margins (except *V. suspensum* with crenulate margins)

References cited regarding Viburnum USDA Hardiness Zones:

Dirr, M. A. 2007. Viburnums. Flowering Shrubs for Every Season. Timber Press, Portland, Oregon USA

Edwards, E. J., D. S. Chatelet, B-C. Chen, J. Y. Ong, S. Tagane, H. Kanemitsu, K. Tagawa, L. Teramoto, B. Park, K-F. Chung, J-M. Hu, T. Yahara, and M. J. Donoghue. 2017. Convergence, consilience, and the evolution of the temperate deciduous forests. Amer. Nat. 190: S87-S104.

Park, B. and M. J. Donoghue. 2021. Phylogenomic insights into the independent origins of sterile marginal flowers in Viburnum. 2021. Int. J. Plant Sci. 182: 591-608.

Schmerler, S., W. Clement, J. Beaulieu, D. Chatelet, L. Sack, M. J. Donoghue, and E. Edwards. 2012. Evolution of leaf form correlates with tropical-temperate transitions in *Viburnum* (Adoxaceae). Proc. Royal Soc. B: Biological Sciences. 279: 3905-3913.

Spriggs, E. L., W. L. Clement, P. W. Sweeney, S. Madriñán, E. J. Edwards, and M. J. Donoghue. 2015. Temperate radiations and dying embers of a tropical past: Evidence from *Viburnum* diversification. New Phytologist 207: 340-354.

Spriggs, E. L., S. B. Schmerler, E. J. Edwards, and M. J. Donoghue. 2018. Leaf form evolution in *Viburnum* parallels variation within individual plants. Amer. Nat. 191: 235-249.

**Column T.** USDA Zones from other sources

Our analyses utilize the zones assigned by Dirr (2007) except as noted below. These are considered the most reliable based on Dirr's expert knowledge of *Viburnum* horticulture. However, because the assignment of species to zones varies from source to source, here we include zones assignments from on-line sources that were considered reliable enough based on the included descriptions and photographs. Searches were carried out using the species name and "USDA Hardiness Zone", and the selected web-site is given in Column U. Although the Dirr scores are not always identical to the online sources, they generally overlap considerable; in no case is there a major discrepancy. "?" indicates that a search was made but that information was not found or was deemed unreliable.

For those few species where Dirr (2007) did not list hardiness zones, we used an on-line source that we considered reliable. This applies for the following species: *V. urceolatum*, *V. glomeratum*, *V. hanceanum*, *V. taitoense*, and *V. oliganthum*.

**Column U.** URLs for USDA Zones from other sources

The source web-site is given for each entry in Column T. Searches were carried out using the species name and "USDA Hardiness Zone". We used only sources that were considered reliable based on the included descriptions and photographs.

**Column V.** Unlikely biome (4-states) based on USDA hardiness zones (freezing tolerance)

O=tropical rainforest; 1=evergreen warm temperate (lucidophyllous) forest; 2=cloud forest; 3=cool temperate forest.

Considering only those cultivated species for which we have reliable data on USDA hardiness zones, we use the hardiness assignments to constrain biome transitions based on the following reasoning. The 15 species assigned above under Column S to the "warm-loving" (freezing intolerant) category are considered unlikely to transition into a cold temperate biome (3) owing to their inability to persist in such climates long enough to evolve freeze-tolerance mechanisms. They can, however, transition into tropical (0), warm temperate (1), and cloud forests (2). Conversely, the 12 species assigned above to the "cold-loving" category are considered unlikely to transition into a tropical biome (0), in which average winter minimum temperatures are high (generally not less than 10-15 degrees F) and no freezing occurs. They could, however, possibly transition into warm temperate (1) and cloud forests (2). All other species, which variously span the freezing mark as described above, are here considered unconstrained in transitioning between biomes.

**Column W.** Unlikely biome (3-states) based on USDA hardiness zones (freezing tolerance)

O=tropical rainforest; 1= extratropical warm forest; 2=cool temperate forest.

Based on the reasoning above (Column U), species in the "warm-loving" (freezing intolerant) category are considered unlikely to transition into a cool temperate biome (2) owing to their inability to persist in such climates long enough to evolve freeze-tolerance mechanisms. They can, however, transition into extratropical warm forests (1). And, species in the "cold-loving" category are considered unlikely to transition into a tropical biome (0), but they could possibly transition into extratropical warm forests (1).

### Supplement 3: *RFBS* Supplementary Analyses

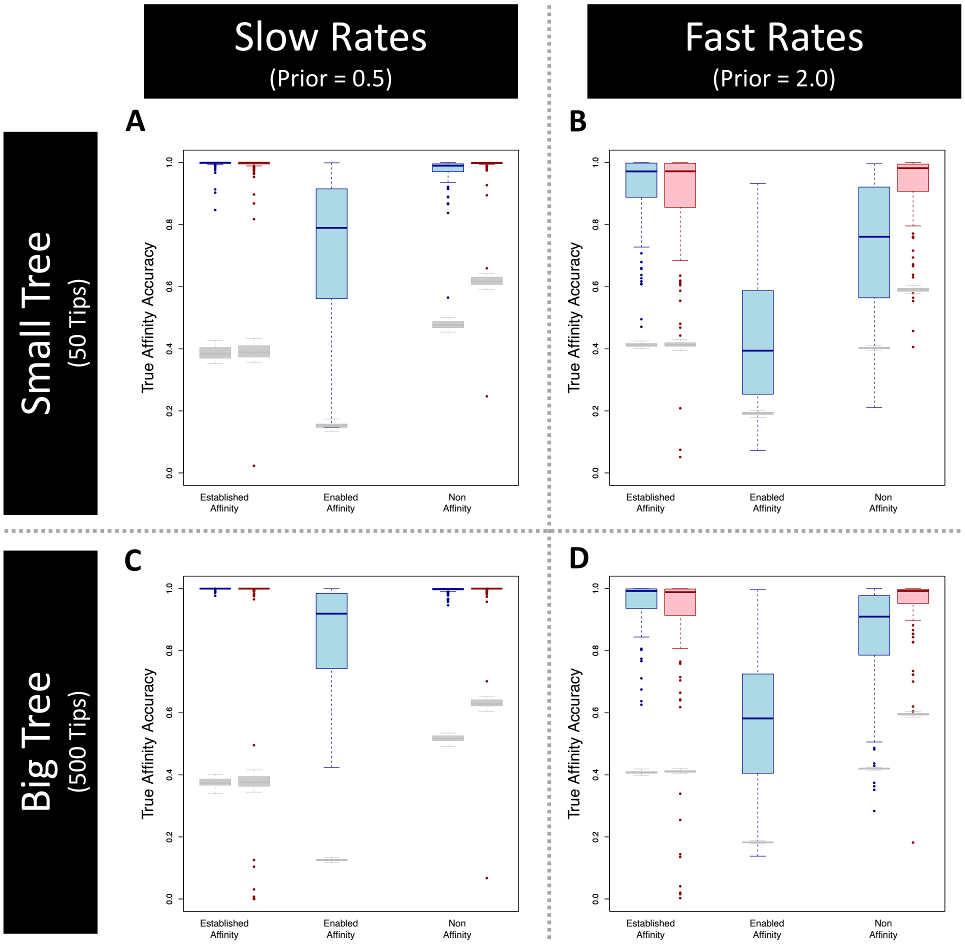


Figure S1: Ancestral affinity accuracy for *RFBS* (blue boxes) and *DEC* (red boxes) simulation analyses .Boxplots are plotting the distribution of support for the true affinity for each biome at each node/corner of the ancestral state reconstructions. Columns of each plot include support for established affinity, enabled affinity, and non-affinity. Four simulation treatments of 100 datasets are shown for two treatments of tree size (Small Tree= 50 tips, Large Tree=500 tips) and exponential prior scaling parameter(Slow=0.5, Fast=2.0). A) Small tree and slow rate treatment, B) Small tree and fast rate treatment, C) Large tree and slow rate treatment, D) Large tree and fast rate treatment.

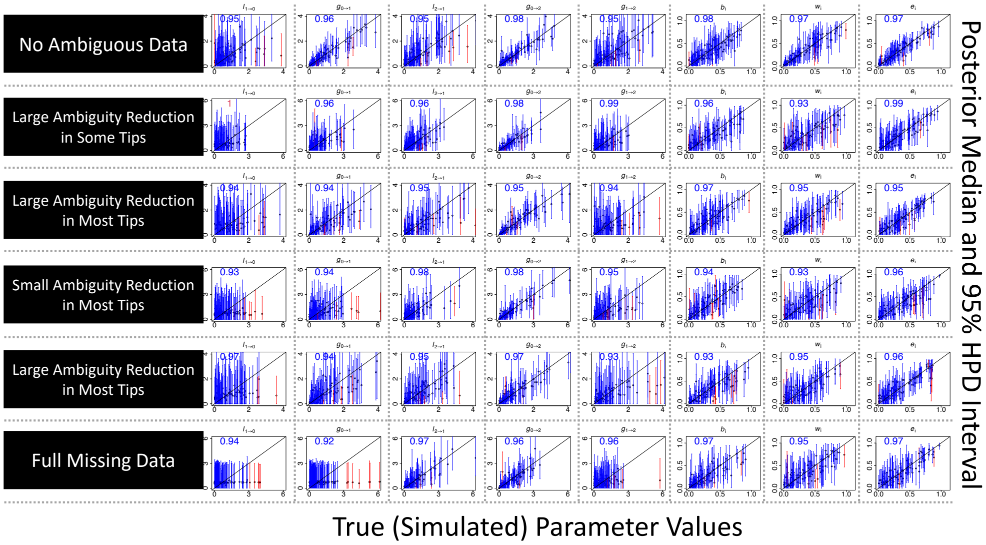


Figure S2: Missing unestablished (enabled or non-affinity) biome affinity data simulation results for model parameters. We resolved either 33% or 66% of ambiguous states that were not the true state, equivalent to removing at most either two ambiguous states (Large Ambiguity Reduction) or one ambiguous state (Small Ambiguity Reduction), respectively, over 25% (Some Tips), or 75% (Most Tips) of tips in the tree, for a total of 4 partially ambiguous treatments, and two additional treatments where all tip states are entirely unambiguous or entirely ambiguous.

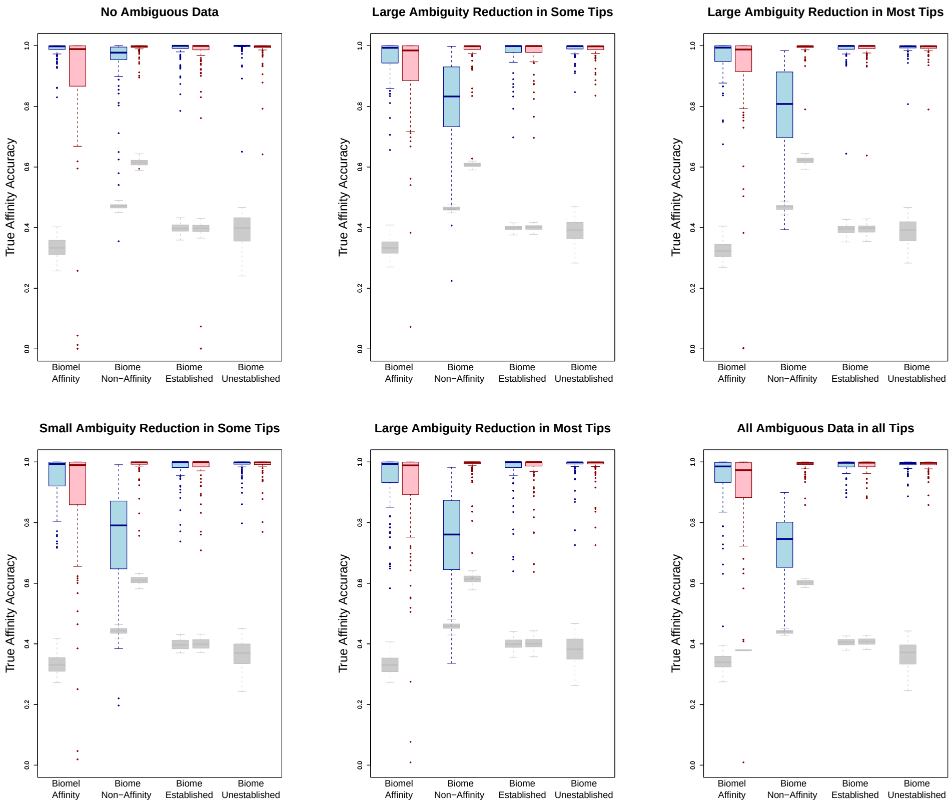


Figure S3: Missing unestablished (enabled or non-affinity) biome affinity data simulation results for *RFBS* (blue boxes) and *DEC* (red boxes). We resolved either 33% or 66% of ambiguous states that were not the true state, equivalent to removing at most either two ambiguous states (Large Ambiguity Reduction) or one ambiguous state (Small Ambiguity Reduction), respectively, over 25% (Some Tips), or 75% (Most Tips) of tips in the tree, for a total of 4 partially ambiguous treatments, and two additional treatments where all tip states are entirely unambiguous or entirely ambiguous. Columns of each plot include support for biome affinity (established/enabled) non affinity, biome occupancy (established affinity), biome non occupancy (enabled/non-affinity).

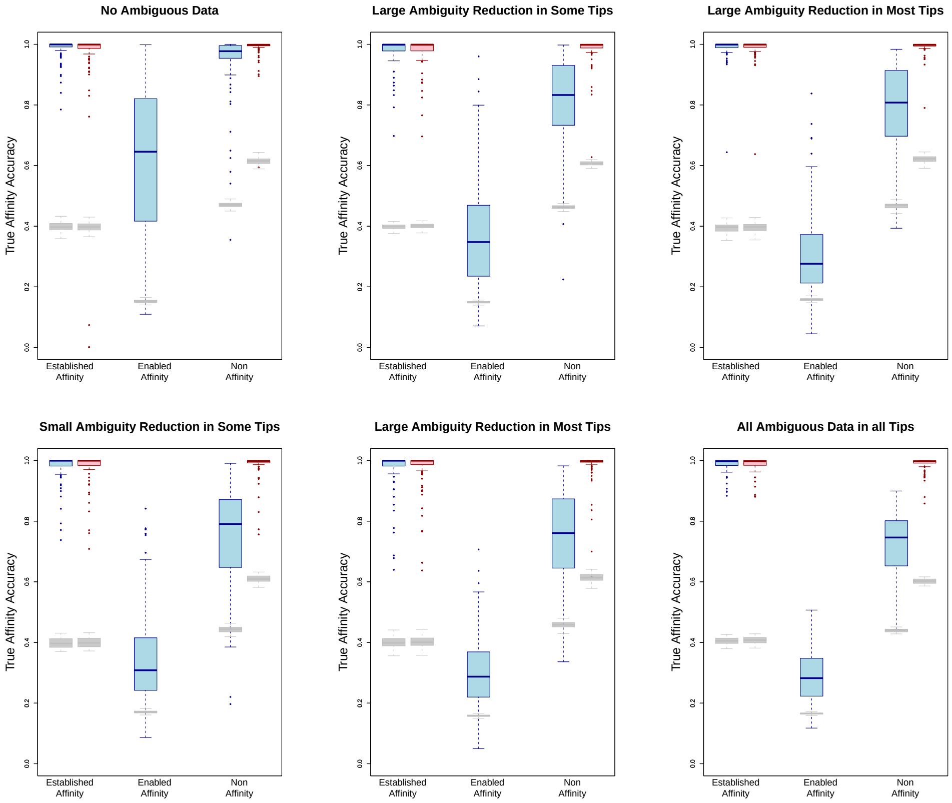


Figure S4: Missing unestablished (enabled or non-affinity) biome affinity data simulation results for *RFBS* (blue boxes) and *DEC* (red boxes) . We resolved either 33% or 66% of ambiguous states that were not the true state, equivalent to removing at most either two ambiguous states (Large Ambiguity Reduction) or one ambiguous state (Small Ambiguity Reduction), respectively, over 25% (Some Tips), or 75% (Most Tips) of tips in the tree, for a total of 4 partially ambiguous treatments, and two additional treatments where all tip states are entirely unambiguous or entirely ambiguous. Columns of each plot include support for established affinity, enabled affinity, and non-affinity.

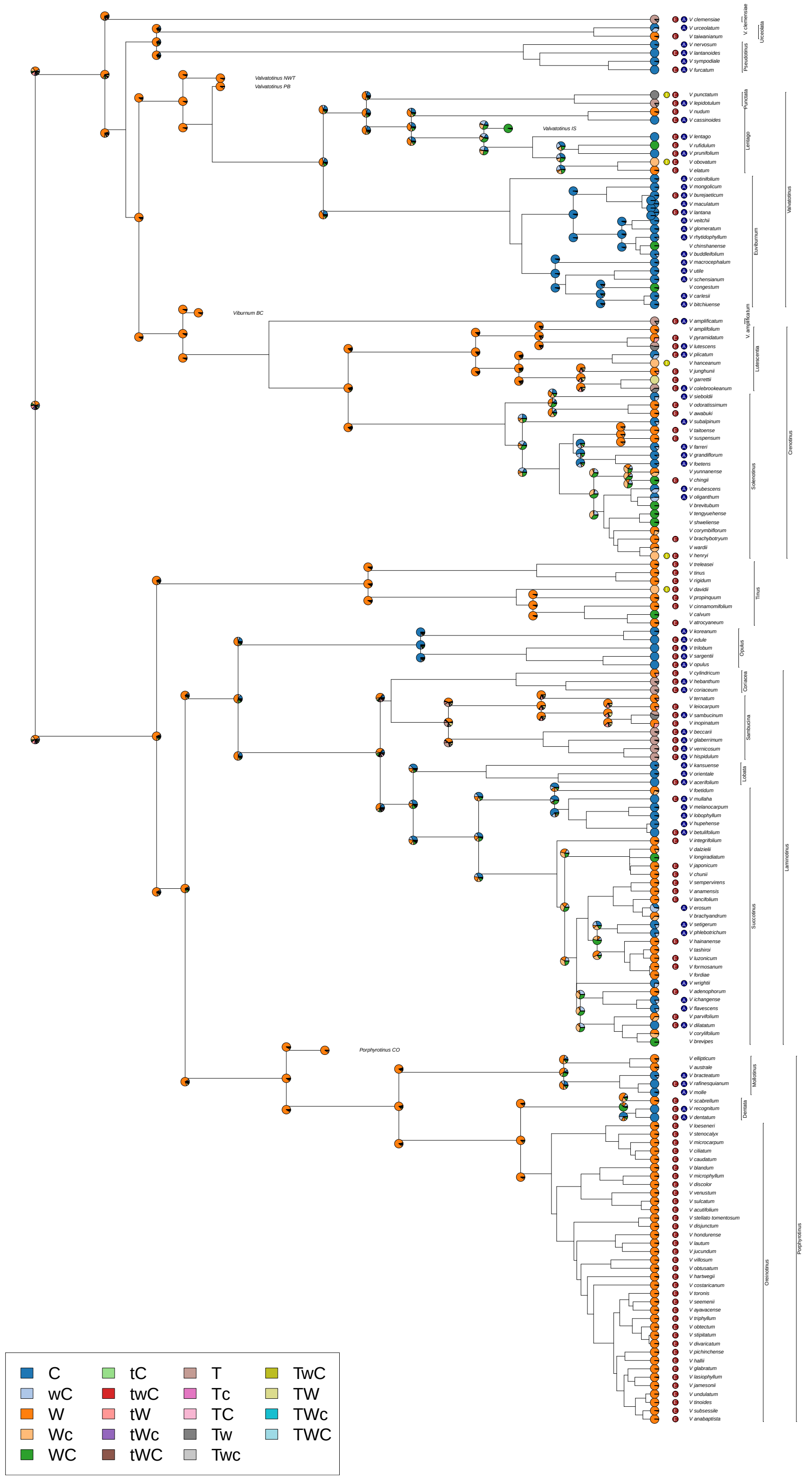


Figure S5: Ancestral state reconstruction of *Viburnum* biome affinities from *RFBS* using conservative included and excluded enabled affinity tip data. Colored circles between affinity probability pie charts and tip labels denote information used to reduce tip ambiguity. Dark red "E": enabled biome affinity excluded; dark yellow "I": established biome affinity included; dark blue "A": climatic adjacency rule applied (i.e., a species cannot have an affinity for cold temperate and tropical biomes with no affinity for intermediary warm temperate biomes).

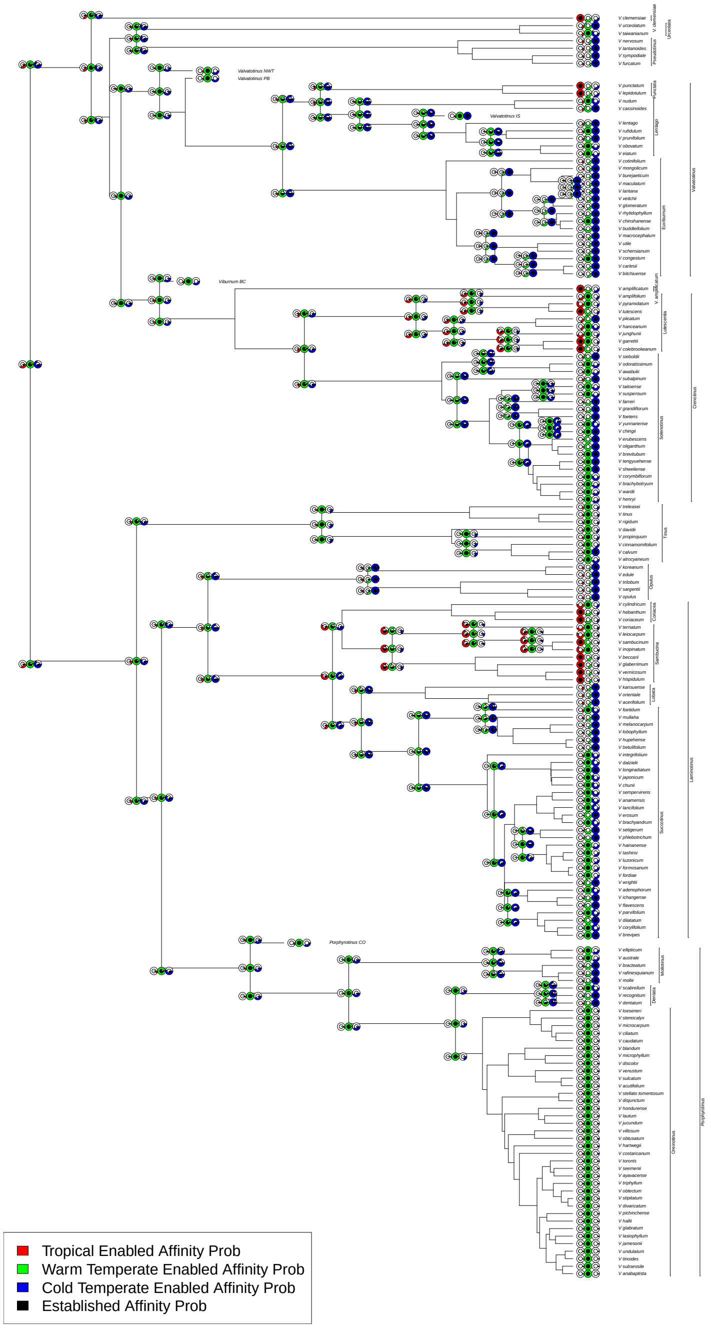


Figure S6: Ancestral state reconstruction of *Viburnum* biome affinities from *RFBS* without using any included and excluded enabled affinity tip data. Node and corner pie chart triplets represent the probability for each biome established/enabled affinity given the probabilities for each state. Outer colored circles represent posterior probability support for enabled affinities (red=tropical, green=warm temperate, blue=cold temperate) while black inner circles represent posterior probability for established biome affinities.

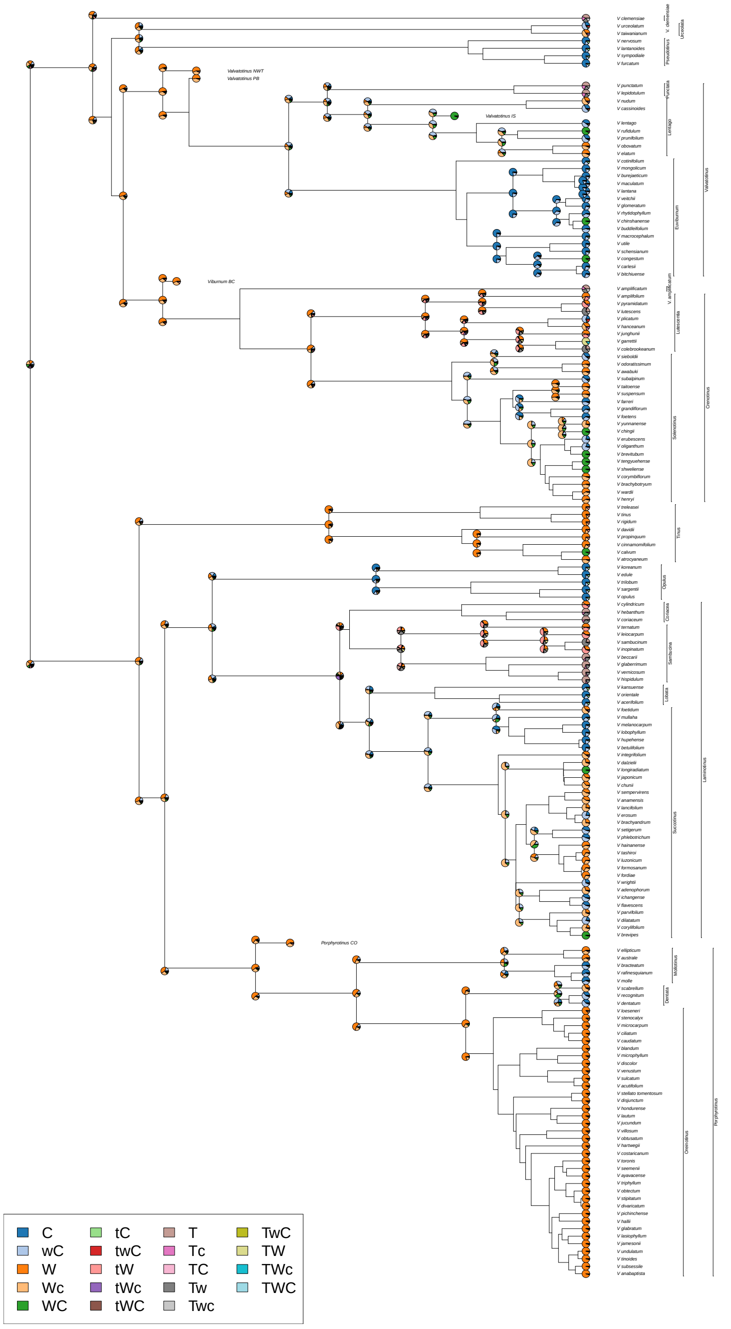


Figure S7: Ancestral state reconstruction of *Viburnum* biome affinities from *RFBS* without using any included and excluded enabled affinity tip data. Legend explains color coding of states, (T/t=Tropical, W/w= warm temperate, C/c=cold temperate). Upper case denotes established and enabled affinity while lower case denote only an enabled affinity, no letter for a biome denotes non-affinity

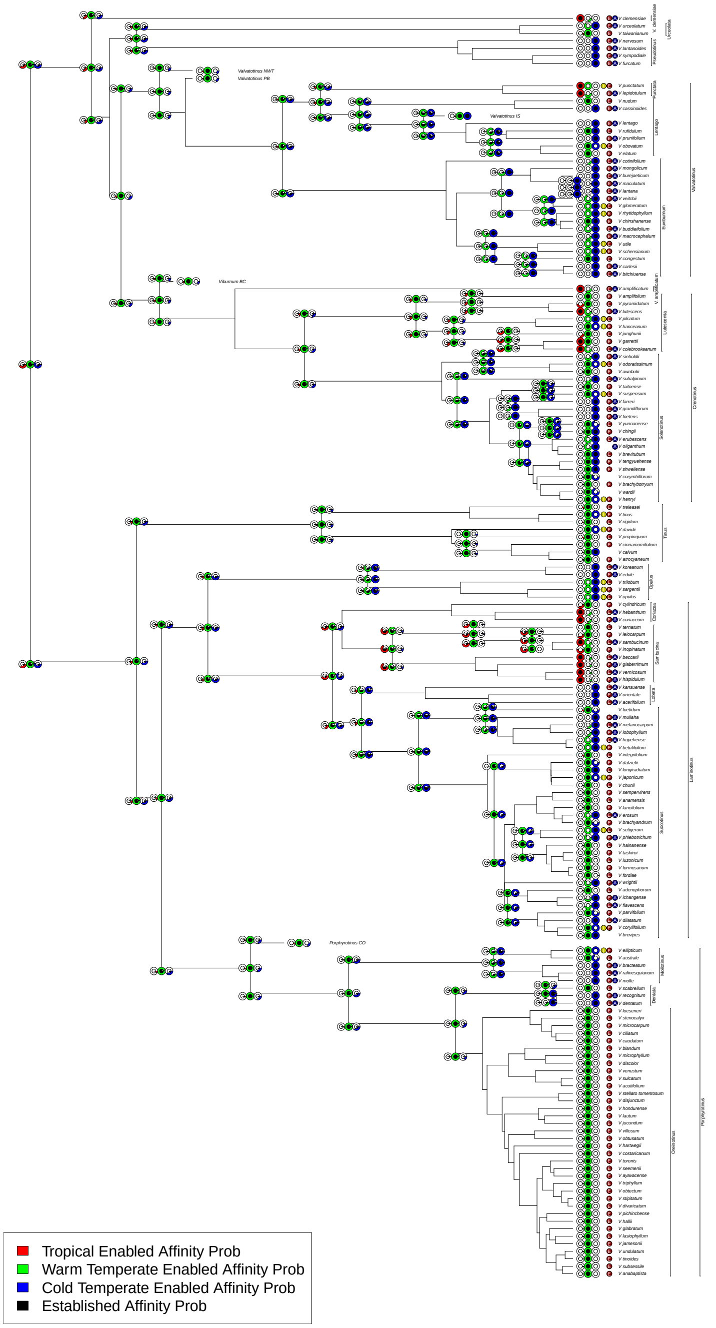


Figure S8: Ancestral state reconstruction of *Viburnum* biome affinities from *RFBS* using bold included and bold excluded enabled affinity tip data. Node and corner pie chart triplets represent the probability for each biome established/enabled affinity given the probabilities for each state. Outer colored circles represent posterior probability support for enabled affinities (red=tropical, green=warm temperate, blue=cold temperate) while black inner circles represent posterior probability for established biome affinities.Colored circles between affinity probability pie charts and tip labels denote information used to reduce tip ambiguity. Dark red "E": enabled biome affinity excluded; dark yellow "I": established biome affinity included; dark blue "A": climatic adjacency rule applied (i.e., a species cannot have an affinity for cold temperate and tropical biomes with no affinity for intermediary warm temperate biomes).

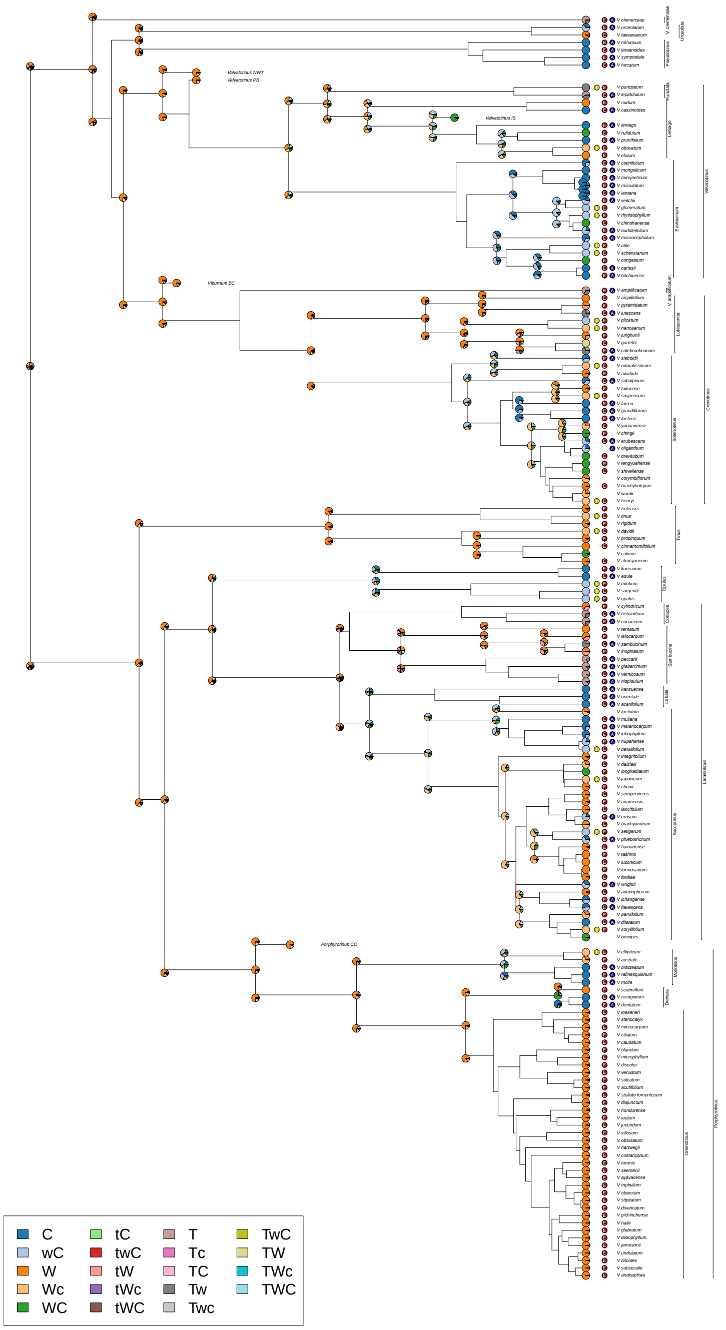


Figure S9: Ancestral state reconstruction of *Viburnum* biome affinities using bold included and bold excluded enabled affinity tip data. Legend explains color coding of states, (T/t=Tropical, W/w= warm temperate, C/c=cold temperate). Upper case denotes established and enabled affinity while lower case denote only an enabled affinity, no letter for a biome denotes non-affinity.Colored circles between affinity probability pie charts and tip labels denote information used to reduce tip ambiguity. Dark red "E": enabled biome affinity excluded; dark yellow "I": established biome affinity included; dark blue "A": climatic adjacency rule applied (i.e., a species cannot have an affinity for cold temperate and tropical biomes with no affinity for intermediary warm temperate biomes).
